# Learning from tandem mass spectra at scale with a self-supervised foundation model for proteomics

**DOI:** 10.64898/2026.09.03.747733

**Authors:** Mechiel Nieuwoudt, Marco Reverenna, Divanisha Patel, Rachel Catzel, Isaac H.J. Houngue, Jemma Daniel, Kevin Eloff, Alberto Santos, Nicolas Lopez Carranza, Timothy P. Jenkins, Jeroen Van Goey, Konstantinos Kalogeropoulos

**Author notes:** Shared correspondence and joint supervision. These authors contributed equally to this work (Mechiel Nieuwoudt, Marco Reverenna, Divanisha Patel). Bindbridge, Barn 4, Park Farm, Villa Road, Impington, Cambridgeshire CB24 9NZ, United Kingdom (Mechiel Nieuwoudt).

## Abstract

Mass spectrometry-based proteomics increasingly relies on machine learning, yet existing models are trained for defined supervised tasks such as peptide identification, *de novo* sequencing or fragment intensity prediction, limiting transfer across datasets, instruments and acquisition methods. Here we present InstaNovo-FM, a self-supervised foundation model for bottom-up proteomics trained to reconstruct masked regions of tandem mass spectra. We assemble a diverse training corpus spanning 1.63 billion MS/MS spectra and 184.6 million high-confidence annotations. We train an encoder-only transformer on the annotated tier using a physics-aware masked reconstruction objective. We demonstrate that the InstaNovo-FM embeddings encode fundamental experimental and biological properties, including fragmentation method, sequence properties and post-translational modifications, without requiring peptide labels. Furthermore, this foundation model directly enables diverse downstream applications, including *de novo* peptide sequencing, database-free identification and analytical run classification. InstaNovo-FM establishes a unified representation space for peptide fragmentation spectra, enabling robust transferability across the proteomics ecosystem.

## Main

Mass spectrometry (MS)-based proteomics has transformed system-wide protein analysis by enabling the study of proteome composition and regulation across diverse biological contexts, including disease, signalling and cellular state [1, 2]. This technology drives central applications ranging from immunopeptidomics [3] and clinical biomarker discovery to the characterisation of organisms across the breadth of biology [4–14]. Public repositories such as PRIDE and MassIVE now hold petabytes of raw mass spectrometry data, a cumulative archive spanning hundreds of thousands of experiments and trillions of individual spectra [15, 16]. Despite this scale, current analytical pipelines remain annotation-dependent. Peptide identification is achieved by matching spectra against a reference database under fixed enzyme rules and a limited modification vocabulary, leaving spectra from unknown peptides, non-standard modifications, and novel sequence contexts unintegrated. The bottleneck is therefore not data but method, as no analytical framework in proteomics learns directly from raw spectra without requiring annotation at every step.

Machine learning already serves as a cornerstone of state-of-the-art proteomics analyses tools [17, 18]. Methods such as Percolator [19], DIA-NN [20] and MSBooster [21] demonstrate substantial improvements in the identification and quantification of peptides. The abundance of MS data has facilitated the training of large task-specific models for retention time and fragment ion intensity prediction [22–24], and *de novo* peptide sequencing [25–29]. Recent large-scale pretraining efforts further highlight the potential of this approach. pUniFind [30] achieves unified peptide-spectrum scoring supporting over 1,300 modifications, DIA-BERT and its data-dependent counterpart DDA-BERT improve acquisition-specific identification through large-scale supervised pretraining [31, 32], and XuanjiNovo scales supervised *de novo* sequencing to large corpora [33]. Despite their scale, these approaches either focus on proteomics datasets of limited diversity, or remain optimised for a single supervised objective.

Foundation models — large-scale models trained on broad and diverse data with self-supervised objectives [34, 35] — have achieved transformative results across biological domains. In the protein sequence domain, this paradigm has yielded models for generative protein design [36, 37], protein language models [38, 39], and structure prediction [40, 41]. These models learn representations without task-specific supervision, and utilize them across a wide variety of downstream applications. In the small-molecule domain, self-supervised models such as DreaMS [42] and LSM-MS2 [43] learn rich molecular representations directly from metabolomics tandem mass spectra without relying on annotated libraries. Recent work has begun to explore analogous approaches in MS/MS-based proteomics [44–46], yet existing pretraining strategies in this domain continue to rely on peptide sequence annotations, constraining learned representations to the chemical space and task objectives of supervised *de novo* sequencing. No existing model learns general spectral representations from peptide fragmentation spectra without incorporating peptide sequence annotations as supervision signals during pretraining.

A general, mass spectrometry-based proteomics foundation model would address these limitations across multiple levels of analysis. At the fundamental level, it would learn the physical relationships between peaks providing a shared spectral context, learned without any sequence annotations. Downstream models can reuse these representations, including for spectra and experimental conditions that annotation-dependent encoders never observe. At the application level, learned embeddings can serve as features for rescoring database search results, initialise *de novo* sequencing decoders with far less labelled training data, and enable retrieval-based identification through embedding-level similarity without explicit database queries [47]. Recent work demonstrates that deep learned representations of mass spectra improve peptide identification across spectral library, database, and hybrid search modes [48], and that end-to-end deep learning database search with contrastive spectrum–peptide objectives outperforms heuristic scoring engines across diverse species [49]. However, these methods require sequence-annotated training data, limiting their scope to the portion of the spectral universe already covered by existing identification pipelines. Crucially, training without sequence annotations avoids the systematic biases inherent to supervised pretraining. Because sequence labels derive from database searches on well-characterised organisms, supervised approaches concentrate signal on the fraction of proteome space that existing databases already cover well, and generalise less reliably beyond it. Here we present InstaNovo-FM, a self-supervised foundation model that learns transferable representations from tandem mass spectra in bottom-up proteomics without using peptide sequence annotations as supervision signals at any pretraining stage. We assemble a diverse training corpus by harmonising metadata using a large language model (LLM)-assisted curation pipeline. Using the reprocessed data, we develop an encoder-only transformer that learns to predict masked regions of spectra and underlying mass spectrometry properties. We demonstrate that the resulting embeddings generalise effectively across downstream tasks, and we release model weights, embeddings, and code publicly to support future development across the proteomics community.

## A large, diverse proteomics pretraining corpus

To build a training corpus reflecting real-world proteomics diversity, we assembled and uniformly reprocessed a massive collection of public mass spectrometry data. Because a self-supervised model’s representations are bounded by the breadth and cleanliness of its pretraining data, we prioritised broad, uniformly-reprocessed coverage over the reuse of any single existing dataset, yielding a corpus that also serves as a community resource. Starting from 26,603 PRIDE submissions deposited before February 2024 [15], we used GPT-4 to extract structured metadata (organism, tissue, instrument model, fragmentation method, acquisition mode, quantification strategy and disease context) from unstructured, free-text sample and data processing descriptions (Fig. 1A). This process produced a queryable catalog from which we selected 92 projects, prioritising biological and clinical diversity across orthogonal parameters. The selected projects span a wide range of biological sources(Fig. 1B), alongside diverse clinical contexts (Fig. 1C). The resulting corpus spans 72 organisms across all major domains of life (Fig. 1D; Supplementary Fig. S1A) and diverse acquisition settings (Supplementary Fig. S1B).

**Figure 1:**
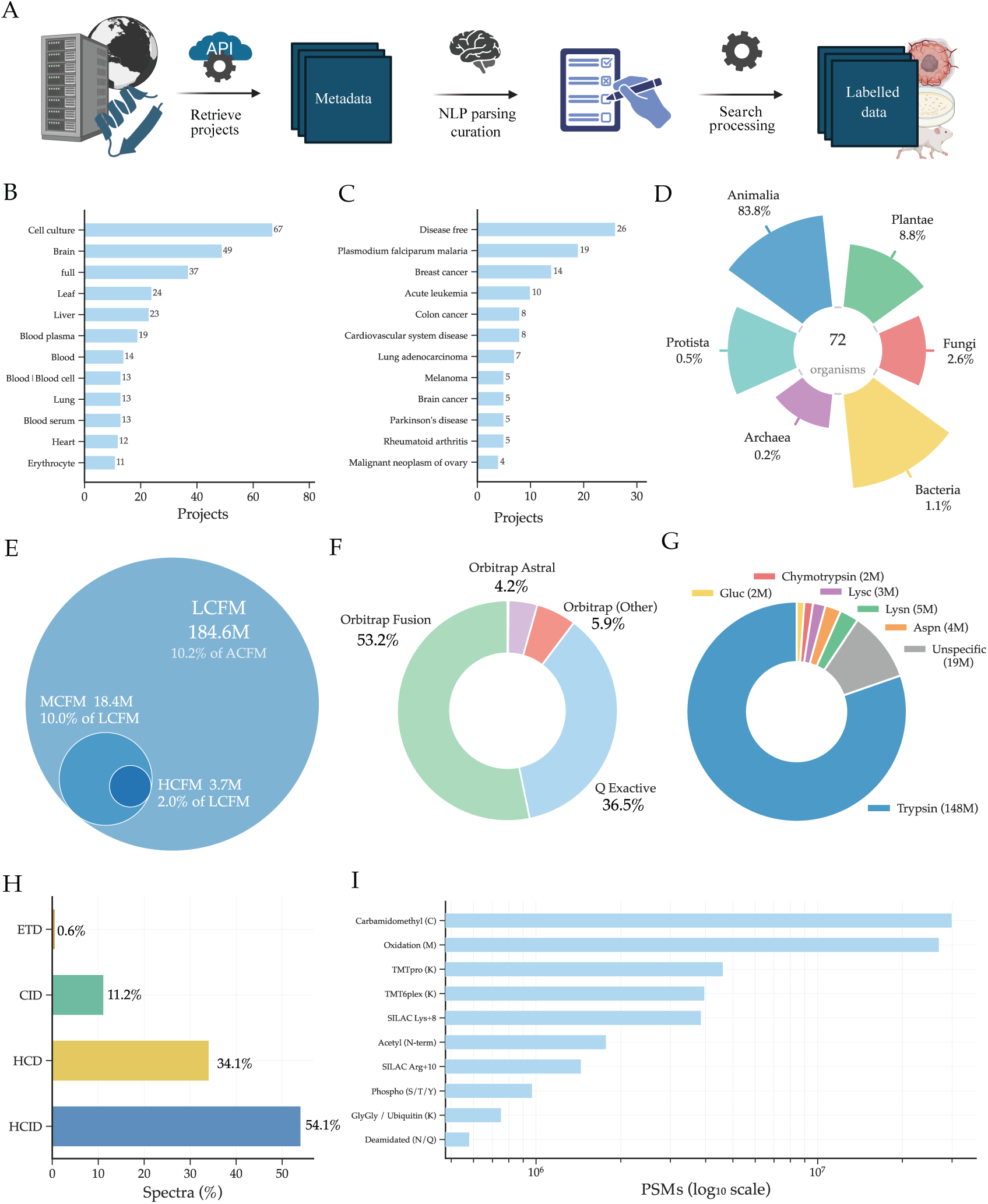
Construction and composition of the InstaNovo-FM pretraining corpus. **A.** The LLM- assisted curation pipeline retrieves projects from public repositories through their APIs, parses and curates the free-text metadata with a natural-language-processing step and processes the raw files into labelled, peptide-annotated fragment data. **B.** Number of projects annotated with each of the twelve most frequent tissue or sample-source terms. **C.** Number of projects annotated with each of the twelve most frequent disease terms. **D.** Kingdom-level taxonomic composition across the organism-annotated projects, spanning 72 organisms. **E.** Nested spectrum counts for the confidence tiers derived from ACFM: LCFM, MCFM and HCFM. **F.** Share of LCFM spectra by mass-analyser family. **G.** Share of LCFM spectra by digestion protease. **H.** Fragmentation method as a percentage of LCFM spectra. **I.** The ten most frequent post-translational and chemical modifications ranked by PSM count.

We reprocessed all raw files through a unified pipeline using FragPipe [21] with 177 experiment-specific workflow configurations. The complete set of 1,625,276,573 MS/MS scans from these files (about 1.6 billion spectra), the vast majority without any peptide annotation, constitutes the All Confidence Foundational Model (ACFM) dataset (Supplementary Fig. S2). Peptide identification at run-specific 1% false discovery rate (FDR) yielded 184,607,213 high-confidence peptide-spectrum matches (PSMs), the labelled subset we term the Low Confidence Foundational Model (LCFM) dataset (Fig. 1E); “low confidence” here denotes the broadest, least-stringently-filtered labelled tier from which more selectively curated higher-confidence subsets (MCFM and HCFM; Methods) are derived for later analyses, preserving parameter distributions across tiers (Supplementary Fig. S3). To prevent data leakage, we implemented peptide-centric splits for training and evaluation. The annotated LCFM tier spans 82 projects and provides a broad distribution of technical and chemical modalities (Fig. 1F). Enzymatic digestion within this tier is expectedly dominated by trypsin (148 million spectra), but we deliberately integrated alternative proteases, including LysN, AspN, and LysC, alongside unspecific cleavage products (19 million spectra) to maximize sequence and cleavage-site diversity (Fig. 1G).

Furthermore, LCFM captures highly heterogeneous fragmentation regimes (Fig. 1H; Supplementary Fig. S1B). At the chemical level, this tier robustly samples the modification space (Fig. 1I). At the peptide level, the labelled LCFM tier comprises 6.52 million unique peptide sequences (4.78 million distinct unmodified backbones). Unlike processed identification tables, tandem mass spectra represent direct measurements inextricably bound to matrix effects, instrument configurations, data acquisition parameters, analyte physicochemical properties and fragmentation dynamics. Capturing this multi-modal heterogeneity is therefore essential for a spectrum-level foundation model, directly guiding our tokenization, embedding strategies and neural architecture.

## Designing a foundation model for proteomics tandem mass spectra

Foundation models for natural language or vision learn from discrete tokens or pixels arranged in structured grids [50, 51]. Tandem mass spectra offer neither. They are unordered sets of continuous-valued (*m/z*, intensity) pairs, with no fixed vocabulary, highly variable length, and a noise floor that dominates peak counts. Building a masked self-supervised objective on this domain requires resolving a sequence of domain-specific problems that have no counterpart in text or image modelling (Fig. 2A; Supplementary Fig. S4).

**Figure 2:**
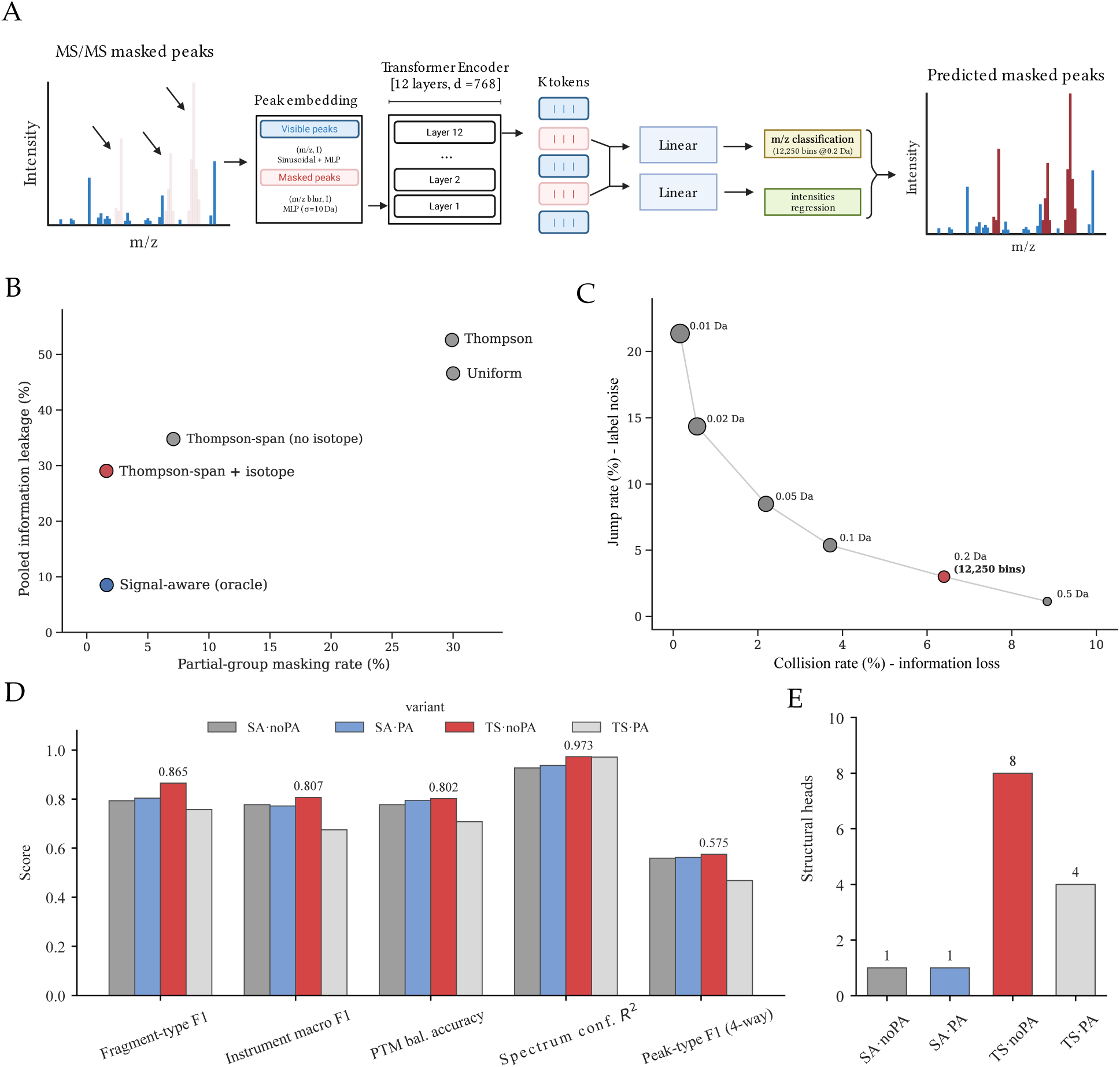
Self-supervised pretraining architecture, masking design and reconstruction objective. **A.** A 768-dimensional encoder-only transformer maps a partially masked MS/MS spectrum to a latent token and per-peak tokens. Four linear heads predict the bin group, within-group offset and *δ*-Da of each masked peak together with its intensity, reconstructing the masked peaks. **B.** Partial-group masking rate versus pooled information leakage across masking strategies. Thompson-span masking with isotope co-masking (red) approaches the signal-aware oracle (blue) without annotations. **C.** Jump rate (repeat observations of a fragment ion falling outside its most common bin) versus collision rate (peaks sharing a bin within a spectrum) across binning resolutions from 0.01 to 0.5 Da; the 0.2 Da grid (12,250 bins, red) gives the best trade-off. **D.** Frozen-embedding probe scores for the four pretraining variants on fragment-type F1, instrument macro-F1, PTM balanced accuracy, spectrum-confidence R^2^ and four-way peak-type F1. **E.** Number of structurally enriched attention heads out of twelve per variant.

Because *m/z* spans three orders of magnitude with chemically meaningful structure at multiple scales, from amino acid mass differences (*∼*57–186 Da) down to isotope spacing (1.003 Da/*z*), a linear embedding would collapse this resolution. We encode each peak using a multi-scale sinusoidal encoder with learnable frequencies at multiple log-spaced scales [52–54], enabling simultaneous resolution of coarse and fine spectral structure. The resulting embedding is concatenated with the peak’s intensity and projected through a second MLP, giving a single 768-dimensional peak token.

In masked language modelling [50], token identity and sequence position are independent. In a mass spectrum, the most natural positional signal is the *m/z* coordinate, which is also the masked prediction target. Encoding *m/z* as position would give the model direct access to the answer through the positional channel, making the self-supervised task degenerate. To resolve this, we encode masked peak positions using a Gaussian-blurred *m/z* value (*σ* = 10 Da) combined with the original visible peak intensity, providing approximate spatial identity for masked positions without revealing the *m/z* reconstruction target. We chose the blur width to be an order of magnitude larger than isotope spacing (*∼*1 Da/*z*), so within-group isotope identity is hidden by the blur while neighbourhood context remains available (Supplementary Fig. S5). We do not apply any learned positional encoding to the peak sequence, with peaks being processed as an unordered set by the transformer.

Across the HCFM dataset, the most highly curated subset of our data corpus, approximately 64% of peaks cannot be attributed to known fragment ions, rising to *∼*77% after subtracting the fraction of peaks expected to match a theoretical fragment ion by chance alone. Yet, these peaks carry only *∼*47% of the total ion current.

Because most of these peaks carry little predictable relationship to spectral context, a uniform masking strategy would spend the majority of the training gradient on them. We therefore designed Thompson-span (TS) masking with isotope co-masking (Supplementary Fig. S6A,B), an annotation-free strategy that targets contiguous *m/z*-sorted spans of 3–4 peaks selected via intensity-weighted Thompson sampling [55], biasing anchor selection toward high-intensity (chemically informative) fragments (Supplementary Fig. S7; Methods, section Thompson masking with isotope co-masking). Isotope co-masking is critical, since if the base ion is masked but any member of its isotope cluster remains visible, the model can reconstruct the base via a trivial arithmetic operation rather than learning fragmentation chemistry.

We benchmark TS masking against two references: naive uniform masking, which masks individual peaks at random, and an annotation-driven oracle that uses ground-truth peptide annotations to mask each theoretical fragment group (Supplementary Fig. S7). Partial-group masking is a local shortcut, as a still-visible isotope or loss sibling reveals a partly-masked group. TS masking with isotope co-masking cuts this from *∼*30% under uniform masking to 1.6%, matching the annotation-driven oracle without using any annotation (Fig. 2B). Pooled information leakage is global, as a surviving complementary or neutral-loss partner elsewhere in the spectrum reveals a masked fragment. Here, TS masking trails the oracle (29.1% vs. 8.5%), as the oracle also masks theoretical neutral-loss children whereas TS co-masks only isotope peers. This residual leakage, intrinsic to the physical redundancy of fragmentation spectra, did not prevent downstream gains. Isotope co-masking is retained in all subsequent models, so “TS masking” hereafter refers to Thompson-span masking with isotope co-masking, the configuration used for every TS·noPA and TS·PA result reported below.

Predicting *m/z* by regression allows the model to satisfy the mean-squared error objective by memorising the *m/z* intensity-weighted mean, without learning peak localisation. We instead frame reconstruction as classification over a discrete *m/z* grid at fixed 0.2 Da resolution. We chose this resolution from twelve candidate strategies spanning fixed-width, fixed-ppm and adaptive families, each scored on two quantities that trade off against one another. The jump rate is the fraction of repeat observations of the same fragment ion that fall outside that ion’s most common bin, i.e. the label noise seen by the classification head; the collision rate is the fraction of peaks within a spectrum that share a bin with another peak. Narrow bins lower collisions but raise jumps, and wide bins do the reverse. The 0.2 Da grid gives the best balance of the two, and is the width whose bin edges align with the instrument mass-error envelope (Fig. 2C; Supplementary Fig. S6C; Methods, section Hierarchical *m/z* binning and prediction head; full twelve-strategy evaluation in section Note 3: Binning strategy evaluation).

With all design choices in place, we asked which peaks the model actually relies on using integrated-gradients attribution [56]. Whereas attention shows only which peaks the model can attend to, integrated gradients score how much each visible peak drives the reconstruction of a masked fragment. For 85% of masked fragment peaks, the ion-ladder neighbour (an ion peak displaced from the masked ion by a single amino-acid residue mass) was present in the spectrum, and therefore available as evidence.

We observe reconstruction signal concentrates on annotated structure. The ion-ladder neighbour emerges as the single most-attributed peak for 34% of masked fragments and appears among the top five for 55.0%, confirming that the model reconstructs real fragment peaks rather than memorising the *m/z* distribution or noise patterns. Masked groups are reconstructed to the correct *m/z* bin with 70.1% accuracy overall. Backbone y-ions are reconstructed more accurately than b-ions (74.3% versus 59.2% bin accuracy; median reconstruction error 89.8 versus 196.4 ppm, both within the 0.2 Da target-bin width), consistent with the dominance of y-ions in HCD tryptic spectra and with their cleaner separation in the peak-type probe. Unsurprisingly, performance degrades above *m/z* 1500, consistent with data sparsity at high mass rather than model capacity. Finally, within a spectrum the model’s confidence ranks annotated fragment ions above unannotated peaks (mean per-spectrum AUROC 0.658), making high-confidence predictions particularly reliable for downstream use (Supplementary Fig. S8).

## Masked peak reconstruction drives broad structural specialisation

To disentangle the contributions of masking strategy (TS or signal-aware, SA) and pairwise attention bias (PA-bias) [57], we trained four variants of InstaNovo-FM (SA·noPA, SA·PA, TS·noPA, and TS·PA) on the LCFM corpus under identical conditions. Each variant was evaluated on a battery of frozen-embedding probes described in the Methods (section Factorial ablation of masking and attention bias), computed on the LCFM validation split.

Across the factorial (full results in Supplementary Table S4), TS·noPA leads on the largest share of structural and spectral probes, achieving the highest fragment type F1 (0.865), instrument identification macro F1 (0.807), spectrum confidence R² (0.973), and PTM balanced accuracy (0.802) (Fig. 2D). Notably, TS·noPA develops 8 out of 12 structurally-enriched attention heads, against four for TS·PA and one for each signal-aware variant (Fig. 2E; Methods, section Factorial ablation of masking and attention bias). Its isotope-spacing enrichment is also the highest, at 1.99*×*: attention between peaks one isotope spacing apart is twice what a uniform spread of the same attention mass would give. Our findings show that TS masking, by forcing the encoder to reconstruct contiguous *m/z*-sorted spans, drives structural specialisation to distribute broadly across the full attention stack (8 of 12 heads on the ablation split, 9 of 12 for the deployed model; Supplementary Fig. S9), whereas signal-aware masking concentrates it in a single specialist head.

Nevertheless, we still observe architectural tradeoffs. SA·PA leads on metrics tied to precursor identity and CID-specific fragmentation patterns: precursor *m/z* R² (0.940), modification class macro F1 (0.631), and CID Recall@1 (0.473), the fraction of CID query spectra whose nearest neighbour in the embedding is another spectrum of the same peptide. Signal-aware masking directly targets annotated fragment groups, and the PA token anchors the embedding to spectral identity by encoding pairwise *m/z* differences between visible peaks. Together, these biases favour peptide-identity representations over broad spectral interpretability. SA·noPA leads on the same retrieval measure computed over all spectra (Recall@1 0.406) and on cross-spectrum ion-identity AUROC (0.874), which measures whether the same fragment ion, seen in different spectra of one peptide, embeds more similarly than a different ion at the same approximate *m/z*. The effective rank of SA·noPA embeddings is also the highest (60.5 versus 53.1 for TS·noPA), making it most advantageous for retrieval tasks requiring broad diversity coverage across a large spectrum pool.

TS·PA is the weakest variant, underperforming on 9 of the 12 evaluated metrics. PA-bias encodes pairwise *m/z* differences from the visible context and is therefore structurally redundant when TS masking already forces the encoder to learn contiguous spectral context. The annotation-free TS·noPA is competitive with signal-aware SA·noPA on structural and spectral probes while not requiring any peptide annotation at training time. SA·noPA has a slight advantage on retrieval fidelity and cross-spectrum ion identity, while TS·noPA leads on structural interpretability, instrument identification, and overall spectral generalisation. However, SA·noPA demands the presence of peptide labels for embedded spectra without a major performance advantage. Therefore, we proceeded with TS·noPA for subsequent development and analysis.

## Corpus scale trades retrieval fidelity for broader linear decodability

Next, we investigated the effect of training dataset size and quality for our self-supervised model. We compare a model pretrained on the curated MCFM tier against the deployed model pretrained on the full LCFM corpus, with both models using identical Thompson-span (TS·noPA) masking. MCFM is a high-confidence, annotation-rich subset of LCFM (*∼*10% of its spectra). LCFM is roughly an order of magnitude larger but retains the lower-confidence peptide-spectrum matches that curation discards, with a larger fraction of peaks left unassigned by the search database (comparison controls and evaluation protocol in Methods, section Corpus-scale evaluation). We observe that training on the full corpus improves the linear decodability of spectral and metadata properties. The deployed LCFM model leads on instrument identification, fragmentation method, hydrophobicity, PTM presence, and precursor-mass and precursor-*m/z* regression, with relative gains of roughly 4–17% across these probes (Supplementary Table S5). The deployed model also leads on precursor charge (macro-F1 0.650 versus 0.602), while spectrum-confidence regression is effectively tied. Conversely, the curated-subset baseline retains an edge on retrieval. It achieves higher duplicate-spectrum Recall@1 and mAP@20 (mean average precision over each query’s 20 nearest neighbours) over the same held-out LCFM pool (Supplementary Table S5). Training on the smaller, higher-confidence subset sharpens the nearest-neighbour structure for same-peptide retrieval, whereas training on the larger, less-curated full corpus yields representations from which spectral and metadata attributes are more linearly decodable. Therefore, both regimes produce strong representations, and the preferable regime depends on the downstream objective: broad property decodability or retrieval fidelity.

Together, these results indicate that a reliable, broadly informative representation can be learned from the larger LCFM corpus. Training on the full low-confidence tier does not so much degrade representation quality as reshape it, improving the linear decodability of spectral and acquisition properties at a modest cost to same-peptide retrieval fidelity relative to the cleaner MCFM subset.

## Embedding space encodes a hierarchy of proteomics knowledge

If InstaNovo-FM has learned the physics and biology encoded in mass spectra, the organisation of its embedding space should reflect this learning not through explicit labelling, but as an emergent property of training. To examine this, we computed UMAP embeddings of 100,000 spectra sampled from the LCFM test set and progressively interrogated the structure at decreasing scales, asking at each level what information has been implicitly encoded (Fig. 3).

**Figure 3:**
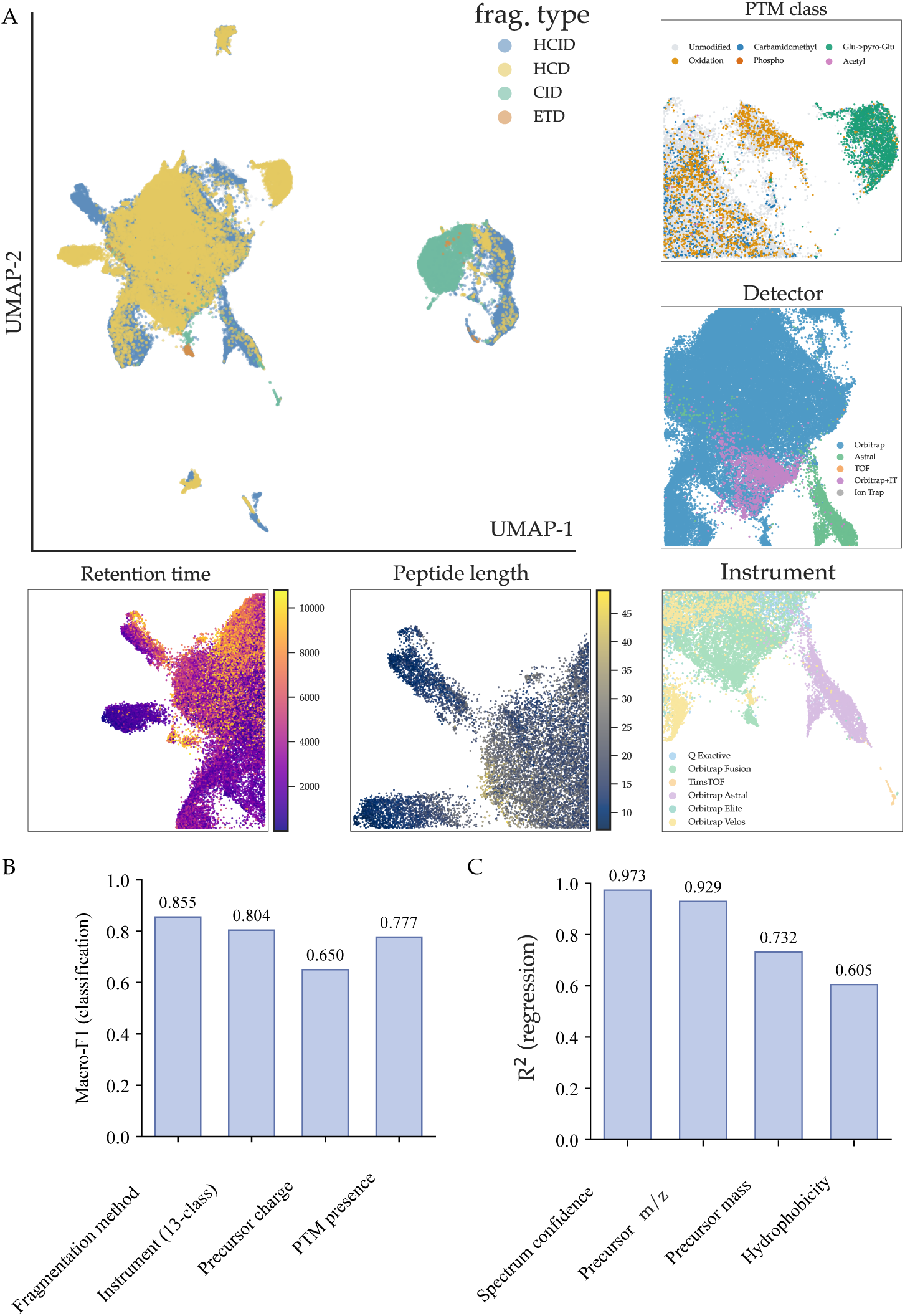
Frozen InstaNovo-FM embeddings encode a hierarchy of proteomic structure without labels. **A.** UMAP of *∼*100,000 held-out LCFM spectra computed from frozen mean-pooled peak-token embeddings and coloured by fragmentation type (HCID, HCD, CID, ETD). Embeddings separate by fragmentation type at the coarsest scale and resolve instrument, labelling, modification and enzyme structure within. **B.** Macro-F1 of selected linear classification probes trained on the frozen embeddings. **C.** R^2^ of selected linear regression probes.

At the broadest scale, the embedding space separates spectra by fragmentation method into three well-defined lobes corresponding to HCD/HCID, CID, and ETD acquisition (Fig. 3A). This result is consistent with each fragmentation method producing a characteristically different ion population, and the model learning these fingerprints. Zooming into the HCD lobe reveals sub-structure corresponding to instrument family (Fig. 3B). Orbitrap Fusion, Q-Exactive, and Orbitrap Astral spectra occupy distinct regions, an organisation the model acquires without instrument labels. This axis, however, largely reflects project-level acquisition context rather than instrument physics. Instrument identity is linearly decodable at macro-F1 0.804 when projects are shared between the probe’s training and test splits, but falls to 0.110 when projects are held disjoint, even though every instrument in the disjoint test split is represented by thousands of spectra in probe training. Within instrument sub-clusters, further organisation emerges by biological and experimental context (Fig. 3C), with embeddings projecting consistently across acquisition modes and interleaving data-dependent acquisition (DDA) and data-independent acquisition (DIA) spectra of identical peptides (Supplementary Fig. S10). Samples labelled with isobaric tags (TMT 6-/10-plex and TMTpro 18-plex) form compact, reproducible sub-clusters irrespective of biological origin. We speculate that reporter and complementary ion signatures imprint spectra strongly enough to override biological variation. This layered structure mirrors the hierarchy of variance components in a typical multi-lab proteomics dataset, and emerges without any of these labels being provided at training time.

The embedding is also organised at the peptide level: replicates of the same peptide are frequently among each other’s nearest neighbours (Recall@1 0.215; Supplementary Fig. S11).

To quantify the information encoded at each level, we trained linear classifiers on frozen mean-pooled peak-token embeddings for a battery of classification tasks spanning technical and biological dimensions (Supplementary Note 6). Fragmentation method is classified with a macro F1 of 0.855, instrument family with 0.804, and precursor charge with 0.650, all far above their majority-class baselines (0.171, 0.025 and 0.104 respectively) and all from a linear probe on frozen embeddings (Fig. 3B; full results in Supplementary Fig. S12).

Together, these probes confirm that there is an embedding hierarchy in spectrum representations yielded by our foundational model, and these are linearly decodable when downstream models are applied. The categories that emerge are physical (fragmentation method, instrument type), experimental (labelling strategy, solvents used), and chemical (peptide identity, modification state). The model has learned, from the spectra alone, which of these factors are the primary sources of variance in the data.

## Peak-level representations reveal learned fragmentation chemistry

The spectrum-level mean-pooled embedding encodes a hierarchy of experimental and biological properties. To understand what information drives this organisation, we descended to the individual peak and questioned whether the model’s peak-level representations encode ion type, and whether its attention mechanism aligns with the physical relationships between ions. We trained a linear classifier on frozen peak embeddings (Methods, section Evaluation framework) to assign each peak to one of four categories: b-ions, y-ions, precursor ions, or unannotated peaks. As a baseline we used the peak encoder output taken before any self-attention layer, which describes each peak by its own *m/z* and intensity alone; on these embeddings four-way classification accuracy was 49.1%, close to the majority-class rate. After the full twelve-layer transformer, four-way accuracy rose to 77.5% (peak-type macro F1 0.575) for TS·noPA (Supplementary Fig. S13). This improvement is attributable entirely to contextual self-attention. The model uses information from surrounding peaks to determine the type of a given peak. Unannotated peaks were separated from annotated ions most cleanly of all (precision 0.950, F1 0.840), confirming that the model has learned a signal-versus-noise distinction without explicit noise labels. Among backbone ions, y-ions were classified more reliably (F1 0.734) than b-ions (F1 0.531), consistent with their dominant presentation in HCD tryptic spectra [28, 58].

To ask whether the model encodes ion identity independently of *m/z* position, we measured the cosine similarity between embeddings of the same fragment ion across different spectra of the same peptide (Methods, section Evaluation framework). TS·noPA achieves a cross-spectrum AUROC of 0.837 for same-ion versus *m/z*-matched discrimination (Supplementary Fig. S14). The mean same-ion cosine similarity was 0.921, compared to 0.830 for *m/z*-matched controls (different ions at the same approximate *m/z*) and 0.697 for random cross-spectrum pairs. We confirmed the same structure with within-spectrum pairwise similarity analysis. Isotopic peaks embed nearly identically to their monoisotopic parents (mean cosine similarity 0.927), neutral-loss peaks remain close to their non-loss parents (0.834), and both exceed the within-spectrum random baseline (0.759). Therefore, the model does not merely encode *m/z*, but also ion identity.

We then turned to the attention mechanism itself. When the model attempts to reconstruct a masked fragment ion, which peaks does it attend to? Masked isotope peaks draw disproportionate attention to their monoisotopic parent and to the same ion in adjacent charge states. Masked neutral-loss ions show enriched attention toward their non-loss counterparts (Supplementary Fig. S9). These relationships emerge without explicit supervision, suggesting that the model has inferred local fragmentation regularities from spectral context alone. To quantify these relationships directly, we measured per-head attention enrichment as the ratio of observed attention between physically related peaks to that expected under a random peak-pair baseline (Methods, section Factorial ablation of masking and attention bias). Attention concentrates on related peaks at mean enrichment factors of 2.18*×* for parent–child (isotope) pairs, 2.13*×* for isotope-spacing pairs (maximum 3.05*×*), 1.88*×* for neutral-loss pairs, and 1.82*×* for backbone fragments attending to their immonium-ion region (Supplementary Fig. S9), consistent with the broad structural specialisation established previously (section Masked peak reconstruction drives broad structural specialisation).

We next examined the unannotated peaks (those outside the standard b/y fragment annotation) that the model relies on, using integrated-gradients attribution on individual spectra (Fig. 4A). Across approximately 42,950 such unannotated peaks, 11.2% match a defined off-database fragment species within 10 ppm, a 2.1-fold enrichment over a sequence-scrambled decoy library and an 18-fold enrichment over *m/z*-shifted nulls, confirming the matches are not coincidental (Supplementary Fig. S14). The attribution, matching, and null-model methodology is detailed in Supplementary Note 5. The dominant category is internal fragments (matching 10.3% of unannotated peaks, products of cleavage at two backbone positions that single-cleavage b/y annotation omits), followed by side-chain d- and w-ions, side-chain neutral losses, and combined precursor losses. Critically, the model assigns these chemically interpretable unannotated peaks systematically higher confidence than unmatched unannotated peaks (median joint confidence 0.26 versus 0.13; AUROC 0.684); Supplementary Fig. S8, Supplementary Fig. S14D), indicating it treats them as predictable signal. For example, in the test-split spectrum of the peptide TGVELGKPTHFTVNAK the model places elevated confidence on unannotated peaks that resolve to the internal fragments int(b,6–7,GK) (*m/z* 186.124, 1.1 ppm) and int(b,2–3,GV) (*m/z* 157.097, 0.9 ppm), both carried at model confidence above 0.99. Internal fragments are written int(ion type, residue range, subsequence), so int(b,6–7,GK) is the b-type internal fragment spanning residues 6–7 of the peptide, GK.

**Figure 4:**
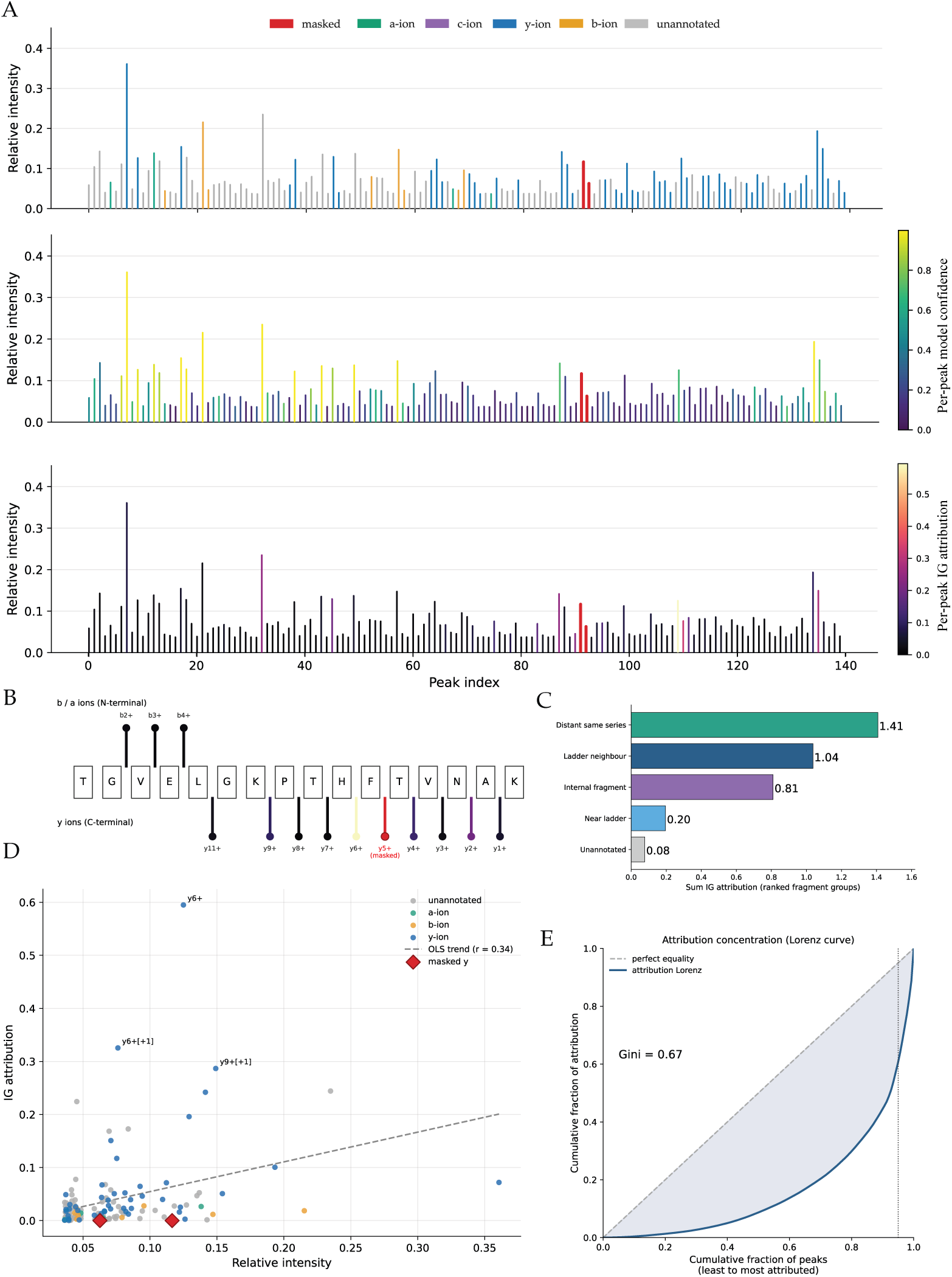
Peak-level attribution reveals learned fragmentation chemistry. **A.** Integrated-gradients (IG) analysis of one masked reconstruction (peptide TGVELGKPTHFTVNAK; HCD, Q Exactive, precursor charge 3+). The same spectrum is illustrated coloured by ion type (top), by per-peak IG attribution to the masked group (middle) and by per-peak model confidence (bottom). **B.** Fragment-ion ladder for the same peptide, with b-ions (N-terminal, above) and y ions (C-terminal, below) coloured by IG attribution. The masked *y*^+^ ion is red and its ladder neighbour 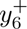 carries the highest neighbour attribution. **C.** Summed IG attribution by ranked fragment-group category. **D.** Per-peak IG attribution versus relative intensity coloured by ion type (OLS trend *r* = 0.34); the masked ions are marked (red diamonds) and the highest-attribution peaks 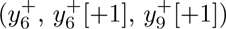 are labelled. **E.** Lorenz curve of attribution across peaks (Gini = 0.67), showing attribution concentrated on a small fraction of peaks.

These patterns are borne out in individual integrated-gradients attributions, in which every peak in a spectrum is scored by its contribution to reconstructing one masked fragment group. In another representative test case, where the model correctly reconstructs the masked *y*_5_ ion of TGVELGKPTHFTVNAK (HCD, Q Exactive, precursor charge 3+), attribution concentrates first on the adjacent ladder ion *y*_6_ (the single phenylalanine step of 147.07 Da, resolved from sequence), together with more distant members of the same *y*-series (the singly- and doubly-charged *y*_9_), reproducing the residue-step arithmetic of the fragment ladder (Fig. 4A,B). The same reconstruction also draws on unannotated peaks that resolve to defined internal fragments (int(b,6–7,GK) at *m/z* 186.124 (1.1 ppm) and int(b,6–10,GKPTH) at 521.283 (0.02 ppm)), showing the model recruiting canonical b/y-ladder chemistry and chemically-defined off-database ions. Across the test split, the top-ranked attributed peak is a ladder neighbour as often as it is an as-yet-unannotated peak (each *∼*28% of masked groups; Fig. 4C). Attribution is only weakly correlated with peak intensity (*r* = 0.34; Fig. 4D) and is concentrated on a small minority of peaks, with the most-attributed 20% carrying most of the total (Gini coefficient 0.67; Fig. 4E). The model is therefore not simply attending to the most intense peaks. Both measures are defined in Supplementary Note 5.

In sum, our results demonstrate that our model, without being told what these peaks are, has learned that they carry structural information, leveraging real fragmentation chemistry (Supplementary Table S7, Supplementary Fig. S14). This extends to unreported or unannotated fragment ions, which can be of potential utility for peptide identification strategies and fragmentation model prediction development.

## Frozen InstaNovo-FM embeddings encode spectral and metadata structure learned without labels

To compare the representations of InstaNovo-FM to representations of models with alternative training strategies, we benchmark with a uniform battery of linear probes across four frozen embedding spaces: InstaNovo v1.2 (IN v1.2) [28], Casanovo v5.0.0 [59, 60], XuanjiNovo (100M MassNet) [33], and InstaNovo-FM. We provide an example of the model embedding spaces with fragmentation type highlighted in Fig. 5A. In each case, a single linear classifier, or ridge regressor for continuous targets, was trained on the frozen model embeddings (see Methods, section External benchmark implementations, for more details).

**Figure 5:**
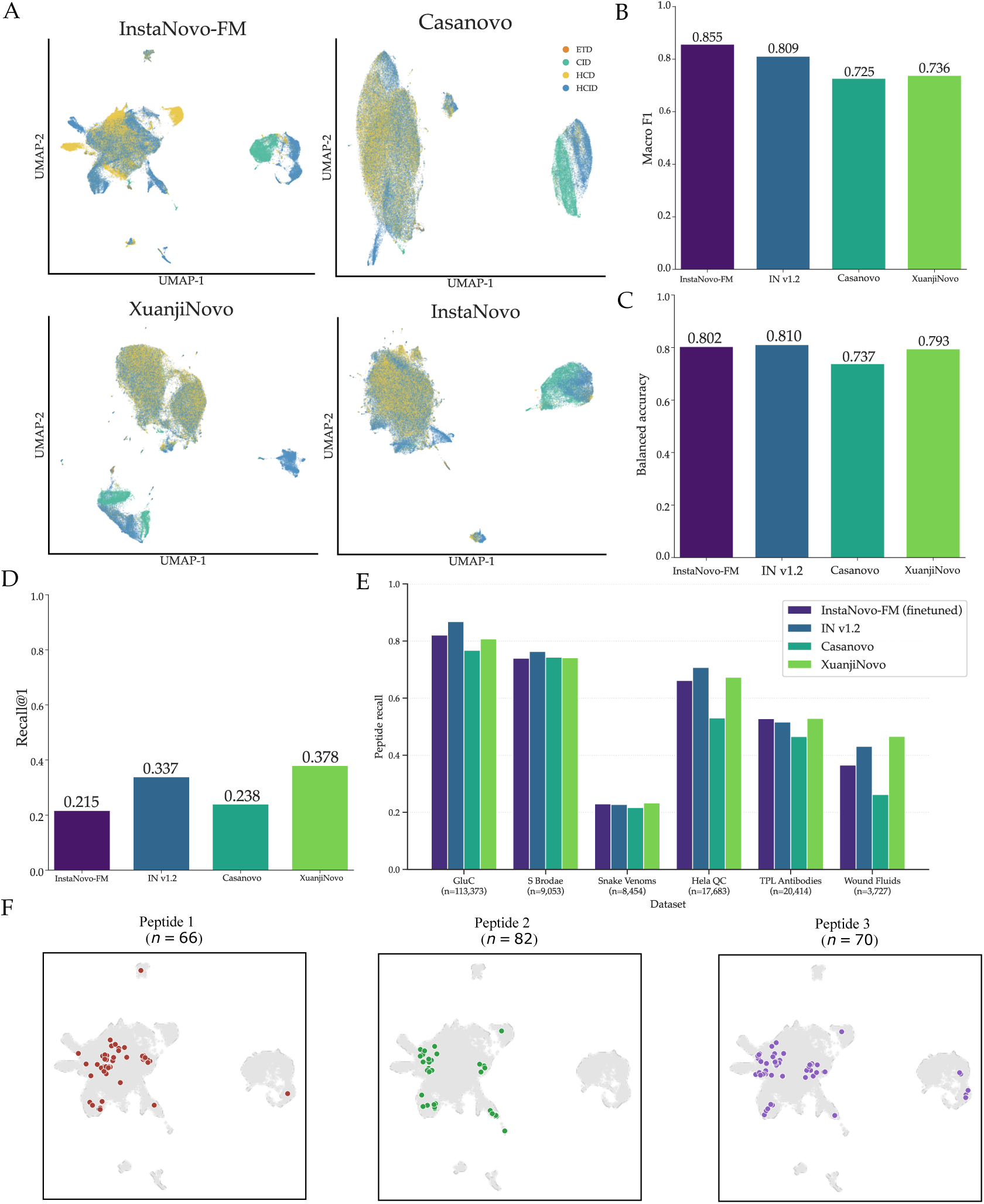
Uniform linear-probe and *de novo* benchmarking across embedding spaces. **A.** UMAP of the same held-out spectra embedded by four encoders (InstaNovo-FM, Casanovo, Xuanji- Novo and InstaNovo v1.2), coloured by fragmentation type. **B.** Duplicate-retrieval Recall@1 per en- coder results. **C.** Linear probe fragmentation type classification macro-F1 results. **D.** Linear probe PTM-presence balanced accuracy results. **E.** Peptide recall for the four models across six held- out biological datasets, with InstaNovo-FM fine-tuned. **F.** Global InstaNovo-FM UMAP highlight- ing the spectra of three example peptides, whose replicate spectra co-locate: Peptide 1, TYFPHFDL- SHGSAQVK (*n* = 66); Peptide 2, EFNAETFTFHADIC[UNIMOD:4]TLSEK (*n* = 82); Peptide 3, AYHE- QLSVAEITNAC[UNIMOD:4]FEPANQMVK (*n* = 70).

Across the linear probe tests, no single encoder dominates, with InstaNovo-FM strongest on acquisition- and fragmentation-related probes and the IN v1.2 strongest on peptide-property and modification probes. InstaNovo-FM outperforms the other models on instrument identification, fragmentation method (Fig. 5C), precursor charge, precursor *m/z* and precursor mass, whereas IN v1.2 leads on PTM presence (Fig. 5D) and modification class (detailed results in Supplementary Table S10). XuanjiNovo leads hydrophobicity and the two duplicate retrieval metrics (Fig. 5B, see Methods, section External benchmark implementations, for details on XuanjiNovo precursor probe exclusions), while Casanovo is ahead in instrument classification. It is important to note that the IN v1.2, Casanovo and XuanjiNovo encoders were all trained with peptide-sequence labels, whereas InstaNovo-FM reaches comparable or improved linear decodability with none.

Our results demonstrate that InstaNovo-FM reaches representations of comparable order without using any peptide-sequence labels. This label-free parity, together with the model’s extension to spectral and run-level properties for which no annotation pipeline exists (instrument, fragmentation, experimental conditions), demonstrate the value of this self-supervised approach.

To further validate the depth of the embeddings from InstaNovo-FM, we benchmarked against IN v1.2, Casanovo, and XuanjiNovo on the task of *de novo* sequencing (see Methods, section External benchmark implementations, for benchmark model implementations, and Methods, section Adapting InstaNovo-FM to *de novo* sequencing, for details on how InstaNovo-FM was adapted to the task of *de novo* sequencing, and model selection).

We compare the peptide recall across the six held-out biological validation datasets on these models (Fig. 5E, Supplementary Fig. S15; detailed results in Supplementary Table S12). Peptide recall varies strongly with dataset difficulty, ranging from *∼*0.23 on the Snake Venoms proteome to *∼*0.87 on GluC. No individual model dominates with IN v1.2 achieving the highest recall on GluC, S Brodae and Hela QC, while XuanjiNovo leads on Snake Venoms, TPL Antibodies (nanobodies) and Wound Fluids. The fine-tuned InstaNovo-FM is competitive throughout, ranking second on GluC, Snake Venoms and TPL Antibodies, trailing the best model by 0.003 or less on the latter two, and outperforming Casanovo on five of the six datasets. We find that spectra originating from the same peptide cluster closely in the InstaNovo-FM embedding space, when not separated by properties such as detector and fragmentation method (Fig. 5F). Additional exploration was done on the impact of training dataset size and model size on model performance, and is detailed in Methods (see section Impact of training dataset and model size on *de novo* sequencing performance).

Overall, these results demonstrate that, after fine-tuning, the self-supervised InstaNovo-FM representations support *de novo* sequencing on out-of-distribution biological samples at a level comparable to the benchmark models.

## Frozen embeddings transfer to downstream modification tasks

To test whether the frozen representations transfer to real biological problems, we applied InstaNovo-FM to three downstream tasks spanning database-free identification, modification analysis, and run-level classification (Fig. 6).

**Figure 6:**
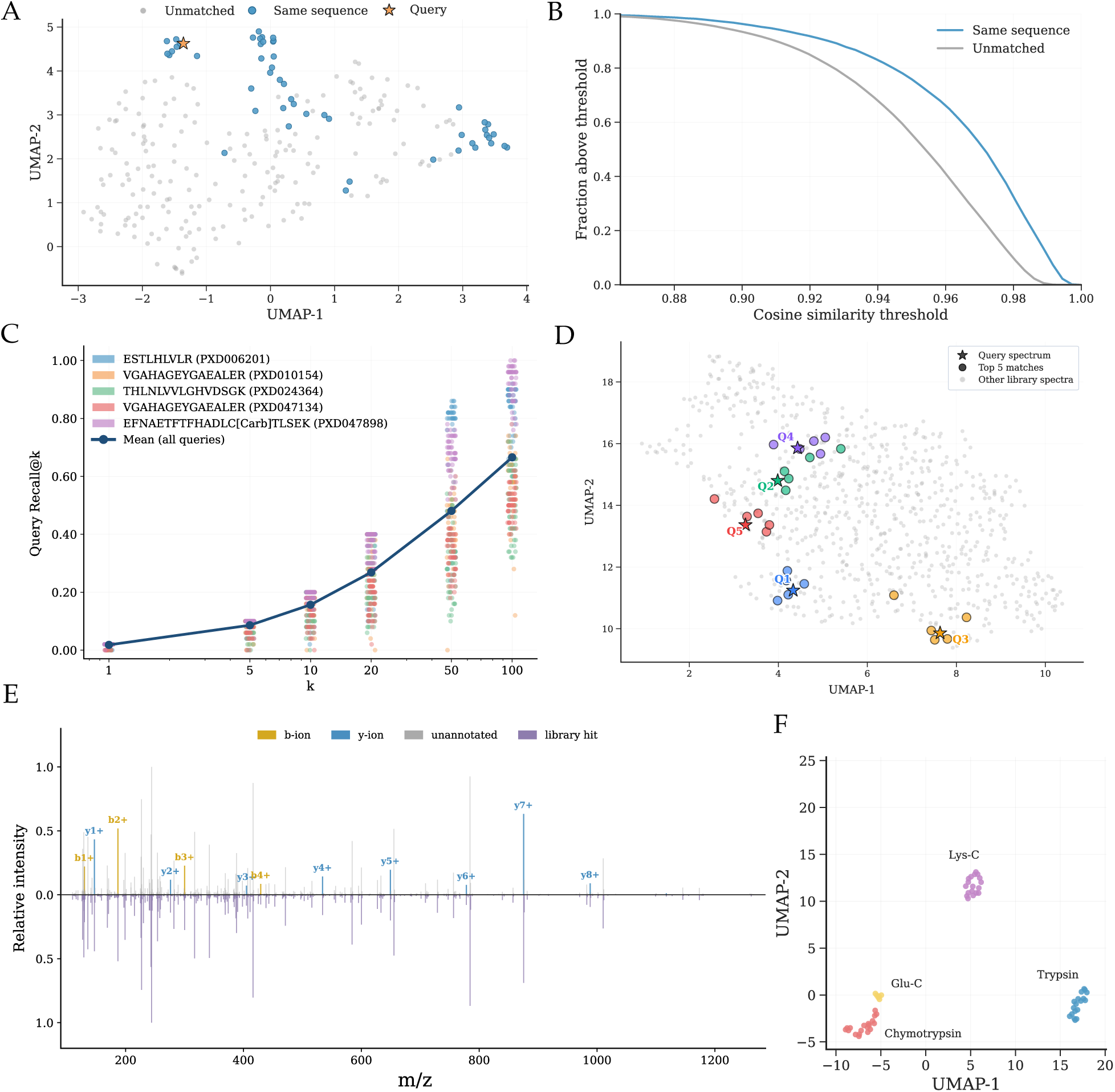
Frozen InstaNovo-FM embeddings enable database-free retrieval, rescue and run-level classification. **A.** UMAP of a controlled retrieval panel: a query spectrum (star) with its 50 same-sequence replicates (blue) among 200 spectra of other peptides (grey), showing same-sequence spectra clustering around the query. **B.** Complementary cumulative distribution of pairwise cosine similarity for same-sequence pairs (blue) versus unmatched pairs (grey). **C.** Recall@*k*: for each query, the proportion of its 50 same-sequence spectra (sharing its exact peptide sequence) found within the top-*k* nearest neighbours by embedding cosine similarity; capped at *k/*50 for *k <* 50. Points show all 250 queries (50 per peptide, across five peptide/project pairs; coloured, sequence and project in the legend), with the mean across all queries in navy. **D.** UMAP of frozen embeddings for a held-out project unseen during pretraining. Five query spectra (stars) with their top-5 nearest library spectra by embedding cosine similarity (matching-coloured circles); remaining reference-library spectra shown in grey. **E.** Mirror plot of a query spectrum (up) against its retrieved library hit (down), with b- and y-ions annotated and shared library-hit peaks highlighted. **F.** UMAP of run-level embeddings for the digestion-enzyme cohort coloured by enzyme (Trypsin, Lys-C, Glu-C, Chymotrypsin), which separate into distinct clusters.

First, we used frozen InstaNovo-FM embeddings as a retrieval index over annotated reference spectra. Given a query spectrum, we defined the top-*k* nearest embedded neighbours with known peptide identifications as candidate matches. This provides a database-free alternative to traditional search, identifying spectra from peptides absent from the local search database as long as they have been observed elsewhere in the corpus. On the held-out LCFM evaluation set, retrieval over frozen TS·noPA embeddings recovers a true same-peptide replicate as the nearest neighbour with Recall@1 of 0.215 overall (0.269 on CID and 0.231 on HCD/Orbitrap spectra) and a mean average precision MAP@20 of 0.076 (Supplementary Table S10).

To visualise the embedding geometry underlying this retrieval, we constructed a controlled panel of five peptide sequences, each with 50 same-peptide replicates searched against up to 200 spectra from other peptides. For a representative query, VGAHAGEYGAEALER, a subset of same-sequence replicates clusters tightly around the query, while the remainder form two additional subclusters at greater UMAP distance (Fig. 6A). These distant subclusters are explained by acquisition context: triply charged precursors (z=3), associated with later retention times and lower spectral quality, separate from the tight cluster of doubly charged (z=2) spectra surrounding the z=2 query. The complementary cumulative distribution of cosine similarity (fraction of pairs above each threshold; Fig. 6B) indicates an anisotropic embedding geometry, with matched and unmatched pairs confined to a narrow high-similarity band; same-sequence pairs are nonetheless shifted toward higher similarity, and retrieval errors arise where the curves overlap. Binary recall@*k* increases from 0.91 at *k* = 1 to *∼*1.0 by *k* = 20 across the 250 reference queries (Supplementary Fig. S16D). When the search depth is sufficient to recover multiple replicates, mean proportional recall@*k* rises from 0.48 at *k* = 50 to *∼*0.67 at *k* = 100 (Fig. 6C).

Applying the same retrieval mechanism to unassigned spectra would enable their rescue. Our approach transfers peptide identifications from labelled anchors to unannotated queries via embedding proximity, validating transfers through physical spectrum-match metrics. To evaluate this, we applied frozen embeddings to a held-out project the model never encountered during pretraining, treating unidentified spectra as queries and database-identified spectra as anchors (Supplementary Table S9). Fig. 6D shows five representative rescues: each query spectrum (star) retrieves its nearest neighbours (circles) from the anchor library, landing tightly around the anchor in embedding space. For each query, the top-1 anchor represents the top-ranked rescued match (embedding similarity *≥* 0.998), while top 2-5 neighbours cluster within 0.985 *−* 0.995. Physical validation confirms top-1 transfers are genuine: across the five rescued pairs, raw experimental spectrum cosine ranges from 0.960 to 0.996 (spectral angles 0.836 *−* 0.933), peak alignment recovers 59 *−* 78% of observed query peaks (*±*0.05 Da), and matched *y*-ion series cover 60 *−* 100% of backbone cleavage bonds (35 *−* 50% of predicted *b/y* ions). The mirror plot (Fig. 6E) illustrates the highest-confidence case for peptide EGLELPEDEEEK (embedding similarity 0.999, raw spectrum similarity 0.994, 369 of 475 query peaks matched, 9 consecutive *y*-ions covering 82% of backbone bonds), where anchor and query fragment ladders overlap nearly peak-for-peak. Embedding proximity, raw spectrum similarity, and fragment-ion evidence thus converge on the same peptide identity, indicating that rescued calls reflect real peptide signal rather than embedding space artefacts.

To evaluate the performance of our representations in exposing specific PTMs present in spectra directly, we train linear probes on frozen InstaNovo-FM embeddings. For individual modifications, we evaluated detection on a stratified set in which phosphorylation- and glycosylation-bearing spectra are enriched to *∼*5,000 positives each per split (at their natural prevalence of *∼*0.3% and *∼*0.1%, a standard 100,000-spectrum probe set would contain only *∼*300 and *∼*100 positives respectively, too few to train and test a probe, so positives are oversampled). Frozen embeddings separate both modifications with high fidelity: phosphorylation is detected with an AUROC of 0.988 (balanced accuracy 0.93) and glycosylation with an AUROC of 0.9999 (balanced accuracy 0.996) (Supplementary Fig. S17A). We interpret this result as glycans imposing a large precursor-mass shift that the embeddings encode, whereas phosphorylation, a smaller though still substantial +80 Da modification, is a more stringent test and is still detected with near-perfect accuracy, indicating that modification-diagnostic structure is linearly accessible in the label-free representation. Beyond binary detection, we asked whether the embeddings resolve glycan composition: a five-way linear probe trained to classify N-glycan family (high-mannose, complex/hybrid, fucosylated, sialylated, and paucimannose) from frozen embeddings reaches a macro-AUROC of 0.915 and a macro-F1 of 0.628 (balanced accuracy 0.677; 71.4% accuracy against a 32.0% majority-class baseline; Supplementary Fig. S17B). Discrimination is strongest for the families carrying the most distinctive mass signatures, sialylated (*F*_1_ = 0.91) and high-mannose (*F*_1_ = 0.79), and weakest for the rare, low-mass paucimannose class (*F*_1_ = 0.29, *n* = 502). Therefore, our representations encode not merely the presence of a glycan but coarse glycan composition. We also probed deamidation of asparagine, a +0.98 Da shift which is near-isobaric to the M+1 precursor isotope. On the MassiveKB dataset, using a within-backbone paired test (2,000 peptide backbones each observed both with and without the modification, isolating the mass shift from sequence context), the frozen embeddings detect it at an AUROC of 0.67, which is above chance but far below the values reached for the large-mass-shift modifications above. These results show that detectability tracks the magnitude of the mass shift a modification imposes and is dependent on spectrum resolution and binning strategy.

Beyond peptide-level tasks, we asked whether frozen, run-aggregated spectrum embeddings encode the technical and biological characteristics of an entire LC-MS/MS run. We hypothesized that runs acquired under the same experimental conditions would yield representations that would cause them to cluster together. We aggregated per-spectrum embeddings into a single run-level representation using three pooling strategies (unweighted mean, the top-1,000 most confident spectra, and a confidence-weighted mean) and evaluated on a technical cohort in which runs differ by digestion enzyme (Trypsin, Lys-C, Glu-C, Chymotrypsin; *n* = 61 runs, with PRIDE identifier PXD065289). Run-level embeddings cluster and classify the digestion enzyme with 100% accuracy under all three pooling strategies, the top-1,000 strategy giving the cleanest separation with 0.779 as silhouette score (Fig. 6F; Supplementary Fig. S16A). Two further cohorts that vary in biological rather than technical conditions, namely stress treatment (PXD074422) and PGM1 knockdown (PXD074720), similarly cluster by condition (Supplementary Fig. S16B,C). Run-aggregated embeddings therefore resolve both technical and biological run conditions with no peptide or protein identifications, supporting database-free representation construction and run-condition classification.

Taken together, these results establish that a single self-supervised model, trained on raw tandem mass spectra without any peptide-sequence labels, learns a representation spanning the full hierarchy of proteomic information. That this representation transfers, with the encoder held frozen, to downstream tasks such as database-free identification, *de novo* sequencing and modification analysis demonstrates that annotation-free pretraining is a practical foundation for proteomics, one that extends naturally to the large fraction of spectra that current annotation-dependent pipelines leave unexploited.

## Discussion

The core finding of this work is that proteomics tandem mass spectra contain learnable chemical structure that self-supervised learning recovers at scale. The structure encoded is not only the instrument and experimental configuration that produced a spectrum, but the identity of individual fragment ions, the presence and position of post-translational modifications, and experimental and biological conditions under which the sample was acquired. The technical barriers to this result are non-trivial: approximately two thirds of peaks in a typical HCD spectrum are unpredictable noise, and the requirements for a masked reconstruction objective on continuous, unordered spectral data differ fundamentally from those for text or images. That frozen embeddings from a model trained without any sequence supervision encode this hierarchy, and support downstream probe and retrieval establishes that the information content of the raw mass spectrum has been systematically underexploited by annotation-dependent analysis pipelines.

A question that arose during design was whether the spectrum embedding would require explicit regularisation to prevent collapse toward a single quality axis, a degenerate solution in which embeddings are ordered only by overall signal-to-noise level and lose the finer chemical and experimental structure. In the final InstaNovo-FM TS·noPA model, well-dispersed embeddings organised by spectral and biological properties emerge without any explicit regularisation loss: the Wang and Isola uniformity metric [61] confirms that the embedding space is well-spread, yet no uniformity or VICReg [62] term appears in the training objective (Supplementary Note 2). The combination of Thompson-span masking with isotope co-masking, which forces the model to predict chemically structured peaks rather than trivially inferring noise level, appears sufficient to prevent degenerate or collapsing embeddings for downstream tasks.

Beyond validating that the model works, the attention analysis reveals what mass spectra encode. The model assigns elevated attention to peaks that do not appear in standard fragment ion annotation approaches, including recurring *m/z* values consistent with immonium ions, diagnostic PTM ions, and non-dominant ion series. The model was never told that these peaks exist or that they are informative. Instead, it learns their relevance from spectral context alone, because they co-vary systematically with the peaks it is trying to reconstruct. This finding suggests that current fragment ion matching approaches are an incomplete description of the spectral vocabulary, and that the learnable signal in mass spectra extends beyond what curated fragment annotation and search engine scoring functions capture. Annotation-free pretraining is therefore not only a way to use unlabelled data, it lets the model learn from chemistry that supervised objectives never reward, because those objectives optimise only against database-derived sequence labels and leave such peaks unmodelled.

At the run level, aggregating per-spectrum embeddings into a single experiment-level representation and classifying the experimental or biological condition of a run without protein identification represents a qualitatively new analysis mode for proteomics. The conventional workflow of peptide identification and quantification, protein rollup, and group comparison, discards any signal not assignable to a sequence match. Spectra from unknown organisms, non-canonical cleavage products, unconventional modifications, and contaminants (contaminants in the context of peptide identification, but informative signal nevertheless) are excluded. Run-level embedding comparison and classification could bypass this bottleneck, with downstream dissection of these differences by machine learning and mass spectrometry tools narrowing down on the discriminant factors.

InstaNovo-FM approaches leading *de novo* peptide sequencing performance without using sequence labels during pretraining, learning directly from fragmentation patterns rather than from sequence supervision; with the encoder held frozen it retains on average 85% of the fine-tuned peptide recall. This is conceptually distinct from peptide-centric *de novo* models, and is timely given recent evidence that supervised sequencing models can over-rely on learned sequence priors, producing peptides that are linguistically plausible but not faithful to the spectrum, or are limited by sequence ambiguity in incomplete spectra [63, 64]. Because InstaNovo-FM sees no sequence labels during pretraining, it cannot by construction encode such sequence-level priors, yet strong peptide-relevant structure still emerges from the fragmentation spectrum alone. Recently developed AI proteomics models that produce generalisable embeddings train encoders on sequence-labelled spectra and achieve strong performance on their target tasks. However, a model requiring high-quality sequence-annotated spectra for pretraining cannot be applied in settings where annotated data does not yet exist. Moreover, supervised pretraining inherits the systematic biases of the annotation pipelines that produced its labels, the same blind spots present in database search itself. An annotation-free objective avoids inheriting these biases through the labels. Just as protein language models derive their power from learning across the full diversity of sequence space rather than the small fraction with experimentally characterised functions, the value of annotation-free pretraining lies in its ability to generalise across conditions that supervised approaches cannot anticipate.

We note, however, that InstaNovo-FM’s current deployment still trains on the LCFM tier, whose membership is itself defined by database-search identifications, so it partially inherits the same corpus- selection bias. Fully realising the annotation-free advantage requires pretraining on the unlabelled ACFM tier, and because Thompson-span masking is annotation-free it applies to ACFM without modification. Whether the added scale of the unlabelled tier improves representations over the cleaner labelled subset, or whether its lower signal concentration offsets the gain, is an open empirical question we leave for future work. In addition, although DIA spectra are already in our corpus, the pretraining objective does not model their chimeric structure, in which wide isolation windows multiplex fragments from co-eluting precursors at unknown charge; adapting it to do so is a natural future direction, given the DIA datasets that increasingly dominate state-of-the-art proteomics.

The interpretability results reported here establish that specialisation is present at the level of individual attention heads, with attention concentrated on isotope, parent–child, neutral-loss and backbone–immonium relationships. What remains unresolved is the finer mechanism, namely how these head-level patterns compose across layers, and which directions in the embedding space encode which properties. Concept-level probing (i.e., identifying which embedding directions correspond to interpretable chemical properties such as peptide hydrophobicity, modification type, or precursor charge) would reveal the geometry of what the model has learned and make downstream predictions chemically auditable.

Several downstream applications are immediately within reach of the current frozen representations. The embedding space already encodes chromatographic behaviour, indicating that sequence-level physicochemical properties are linearly encoded in the spectrum embedding. Spectrum quality and peptide suitability prediction, assessing whether a given MS2 scan contains a peptide signal worth further analysis, is also a linear probe task with immediate value for real-time acquisition control and identification searches. PSM rescoring using InstaNovo-FM embeddings as additional features for a scoring engine could represent a direct complement to conventional database search, potentially recovering identifications that search scores alone miss. Beyond serving as input features, the peak relationships the model makes explicit (complementarity, isotope structure, and neutral losses) could also inform the design of downstream algorithms themselves, such as dynamic scoring functions or search heuristics that exploit this learned spectral context.

An interesting extension of this work would be a sequence-conditioned generation head built on the InstaNovo-FM backbone. Such a model could predict fragmentation spectra from peptide sequences, leveraging the physical chemistry the encoder has internalised to improve upon current spectrum prediction models. Another possible extension remains cross-modal integration, such as a joint embedding space aligning spectral and protein sequence representations, potentially enabling retrieval and inference that bridges measurement modality and sequence identity without database search. Finally, whether self-supervised spectral performance scales smoothly with data volume and model size, as observed in language and protein modelling, is a central question directly relevant to extensions of this work.

## Methods

### Dataset construction and curation

We retrieved metadata from 26,603 projects in the PRIDE repository [15] via the PRIDE API, applying a submission cutoff date of 1 February 2024. For each project, we sought to collect comprehensive details on organisms, instruments, experiment type, modifications used, acquisition methods, and study objectives. We employed a multi-stage selection strategy that combined automated filtering with LLM-assisted metadata parsing. We prioritised projects based on: (i) organism diversity across the tree of life; (ii) instrumentation variety (mass analyser, detector, fragmentation method); and (iii) experimental depth (PTMs, chemical labelling, acquisition mode). A GPT-4-based pipeline was used to parse unstructured project descriptions and extract structured metadata, yielding a final curated collection of 92 project accessions in the assembled corpus. We selected 46 large-scale studies via automated annotation [10, 65–106], 8 landmark studies manually included for data quality and community importance [12, 107–111], 5 datasets from the ProteomeTools project [23, 112–114], the 9 projects comprising the standard nine-species benchmark [115–124], and 24 complementary studies targeting specific metadata gaps to ensure broad chemical coverage [125–144]. Supplementary Table S1 lists all 92 accessions grouped by these categories. All raw files were individually annotated with instrument-specific and experiment-specific metadata.

### Spectral processing and peptide identification

We downloaded raw MS files from PRIDE via FTP using Filezilla (3.67.1), retaining only vendor-specific formats (.raw, .d, .wiff), totalling approximately 26 TB. We converted all raw files to mzML format using ProteoWizard MSconvert (v3.0.23211). We reprocessed all studies using FragPipe (v22.0) with MSFragger (v4.1), optimising search parameters per file according to acquisition mode (DDA or DIA), fragmentation method (HCD, CID, ETD), and relevant PTMs, using 110 reference proteomes and 177 distinct FragPipe workflow configurations. We applied false discovery rate (FDR) control at 1% at the PSM, peptide, and protein levels using MSBooster. We quantified DDA files with IonQuant (v1.10.27) and DIA files with DIA-NN [20] (v1.8.2). Peptide modification annotations in EncyclopeDIA notation were mapped to Proforma UNIMOD identifiers, comparing theoretical precursor mass-to-charge values against masses calculated from the annotated sequence to verify the translation. Precursor charge states were checked for consistency with acquisition mode: DDA spectra were required to carry non-zero charges, whereas DIA spectra were required to carry zero charge, reflecting the unknown charge state in wide-window acquisition; rows violating this rule were removed. Intensities were max-transformed per spectrum. Finally, all spectra were converted to Parquet format for efficient disk storage and fast data retrieval during model training.

### Confidence stratification

We constructed three labelled datasets at progressively stricter confidence thresholds from the complete PSM collection. The LCFM dataset comprises all spectra with PSMs at 1% FDR. The MCFM and HCFM datasets were derived as nested subsets of LCFM by ranking PSMs with a composite confidence score and applying global thresholds on the pooled score distribution.

For each PSM, four search engine-derived quantities were considered: inverted expectation value, search-engine probability, hyperscore, and, when reported, the difference between hyperscore and the runner-up score (nextscore). Because hyperscore and related discrimination metrics are length-dependent, each term was first converted to a percentile rank within equal-length groups, using average-rank ties and mapping ranks to the unit interval. The composite score was defined as the arithmetic mean of the three or four ranked components (three when nextscore was absent). Global thresholds were set as the 90th and 98th percentiles of the composite score over all finite-scored PSMs: MCFM retained PSMs strictly above the 90th percentile and HCFM retained PSMs strictly above the 98th percentile. Sequences containing glycosylation PTM annotations without a matching UNIMOD accession were excluded from scoring and thus the exported medium- and high-confidence subsets.

### Dataset splitting and preprocessing

Train, validation and test partitions were constructed on a peptide-centric basis with target proportions of 80%, 10% and 10%, respectively. Peptide identity was normalised prior to assignment by removing PTM annotations and mapping isoleucine to leucine. A central peptide registry recorded the split assignment for every normalised sequence; all spectra annotated with the same normalised peptide were assigned to the same partition, ensuring that no peptide appeared in more than one split and preventing leakage across confidence tiers. Rather than drawing a fresh 80/10/10 split on LCFM, we extended the registry of peptide assignments established for prior InstaNovo training corpora. Peptides already present in the registry retained their existing partition; only newly encountered peptides were assigned by deficit-proportional sampling toward 80/10/10 targets. This preserves compatibility with earlier InstaNovo [28] splits, enabling incremental addition of LCFM data for future model development. The registry was published alongside the model release to support reproducible extension of the training corpus. MCFM and HCFM inherited split assignments from LCFM. Before registry update or partition writing, spectra were retained only if they satisfied all of the following: retention time at most 10,800 seconds; lower isolation offset at most 300 Da; precursor charge between 0 and 7 inclusive; precursor *m/z* at most 2,000 Da; and no modification annotation that could not be resolved to a UNIMOD identifier. A precursor charge of 0 denotes a DIA spectrum, for which no single precursor charge state is assigned. DIA accounts for 3.6% of the LCFM training partition (Supplementary Table S15). Of the 184,607,213 LCFM PSMs, 181,777,591 (98.5%) satisfied these criteria. Retention time accounted for the large majority of exclusions, and the unresolved-modification criterion for most of the remainder. No PSM exceeded the charge or precursor-*m/z* bounds (Supplementary Table S14). The unresolved annotations are predominantly N-glycan compositions absent from UNIMOD; glycopeptides whose glycan carries a UNIMOD identifier are retained (Supplementary Tables S16 and S17).

Upon data loading, peaks below 1% of the base peak intensity were removed, and the top *N* = 200 peaks by intensity were retained. Intensities were square-root transformed and *ℓ*_2_-normalised per spectrum.

### Foundation model architecture

InstaNovo-FM is an encoder-only transformer with model dimension *d* = 768, 12 self-attention heads (head dimension 64), 12 transformer layers, and feedforward dimension 3072, totalling approximately 89.5M parameters (Supplementary Fig. S4; the full training configuration is given in Supplementary Table S8). The peak encoder and the treatment of positional information are detailed in section Note 1: Model architecture details. Dropout of 0.1 is applied to the attention-output and feedforward residual branches of every encoder layer, and within the peak-encoder MLP. Each peak is embedded using a multi-scale sinusoidal encoder that applies learnable frequencies at multiple scales to the *m/z* value, followed by an MLP projection. Masked peak positions are encoded using a Gaussian-blurred *m/z* value (*σ* = 10 Da) combined with the original visible peak intensity; this provides approximate spatial identity for masked positions without revealing the *m/z* reconstruction target. No learned positional encoding is applied to the peak sequence. A learnable latent token ([CLS]) is prepended to each input sequence. The fixed-dimensional spectrum embedding used for all downstream tasks is the mean of the final-layer hidden states over the non-padding peak tokens, excluding the [CLS] token; this uniform mean-pooled representation was selected over the [CLS]-token hidden state during evaluation. The deployed configuration passes no metadata tokens to the encoder: no instrument, detector, fragmentation-type, collision-energy or precursor feature is provided as input. The metadata recovered by the linear probes is therefore inferred from peak structure alone rather than read back from a conditioning token. Padding masks ensure that zero-padded positions do not contribute to attention or to the pooled embedding. Scaled dot-product attention (Flash Attention backend) [145] is used in all layers. No pairwise attention bias (PA-bias) or rotary position embedding is applied. PA-bias, as introduced by Lapin *et al.* [57], adds a learned scalar bias to each attention logit computed from the *m/z* difference between the query and key peaks, so that fragment pairs separated by chemically meaningful mass gaps (for example, a residue mass or a neutral-loss offset) can be preferentially attended to; we evaluate it as one axis of the four-variant ablation (section Masked peak reconstruction drives broad structural specialisation) and find it is not required by the deployed model.

### Hierarchical m/z binning and prediction head

Continuous *m/z* values are discretised for the classification objective using a fixed-resolution 0.2 Da binning strategy. The 0.2 Da bin width was selected by a safety-margin criterion: the empirical P95 mass-error envelope for CID ion-trap data is approximately 0.09 Da; a 0.2 Da floor guarantees classification targets remain stable under the worst-case mass errors encountered in the training corpus. Spanning *m/z ∈* [50, 2500] Da, this yields 12,250 bins. Predictions are decomposed hierarchically: a coarse group softmax predicts the bin group (245 groups of 50 bins each), followed by a fine offset softmax within the selected group. Group and offset losses are weighted equally at 0.5 each. Intensity is predicted as an auxiliary task: a two-layer MLP on the per-peak hidden states regresses the normalised peak intensity (sigmoid output), trained with a Huber loss (*δ* = 0.05) at weight *λ*_intensity_ = 0.2. This auxiliary head is used during pretraining only; its predictions are not read downstream (only the frozen embeddings are).

### Thompson masking with isotope co-masking

During pretraining, masking targets a 30% fragment-group budget rather than a fixed peak fraction; the per-peak mask ratio is not capped, so for Thompson-span masking with isotope co-masking this realises approximately 26% of peaks per spectrum (26.6% on LCFM). Anchor peaks are selected using intensity-weighted Thompson sampling [55], biasing selection toward high-intensity (chemically informative) peaks: each peak is scored by a draw from a Beta distribution whose parameters are set by its normalised intensity, and the highest-scoring peaks are taken as anchors. Sampling the score rather than ranking on intensity directly keeps anchor selection stochastic, so the same spectrum is masked differently across epochs and low-intensity peaks retain a non-zero selection probability. The name refers to this sampling scheme and not to the thomson (Th), the unit of *m/z*. Each selected anchor is expanded into a contiguous *m/z*-ordered span of 3–4 peaks. Isotope peaks of masked anchors are co-masked using the isotope spacing formula Δ(*m/z*) = 1.003355*/z* Da, where *z* is the fragment charge state, preventing trivial reconstruction from visible isotopic partners. Isotope peaks are co-masked regardless of whether they fall within the original span. Masked peak positions are encoded using a Gaussian-blurred *m/z* value (*σ* = 10 Da) combined with the original visible peak intensity, providing approximate spatial identity without leaking the reconstruction target; no learnable mask token is used. The model is trained to predict the group and offset bin indices of the original *m/z* values using cross-entropy loss.

### Training configuration

InstaNovo-FM was trained for *∼*230,000 steps on LCFM (*∼*117 million training spectra, approximately two epochs) with batch size 1,024 on four NVIDIA H100 GPUs using Hugging Face Accelerate with mixed precision (fp16) and gradient checkpointing. The learning rate schedule follows a cosine warmup-hold-decay pattern: linear warmup over 5% of steps to a peak rate of 10^−4^, hold for 30% of steps, then cosine decay to 10^−5^. Gradient clipping was applied at norm 1.0. In single-GPU training configurations, the model was compiled with torch.compile for execution efficiency. Model selection used median absolute error in PPM (median ae ppm) on the validation set, with checkpoints saved every 10,000 steps (top-3 retained).

For the corpus-scale comparison (section Corpus scale trades retrieval fidelity for broader linear decod- ability), we additionally trained a smaller MCFM baseline: a 9-layer Thompson-span (TS·noPA) model of ∼40 million parameters, trained for ∼90,000 steps on the curated MCFM subset. It shares the deployed model’s masking strategy and reconstruction objective, differing only in depth, parameter count, training budget, and training corpus. Both checkpoints were evaluated identically on the same held-out LCFM spectra: frozen logistic-regression and Ridge probes fit with cuML on 100,000/10,000/10,000 train/validation/test splits, and duplicate retrieval over 20,000 peptide groups (Supplementary Note 6).

### Theoretical fragment ion annotation

We annotated a subset of 200 HCFM validation spectra using the pyOpenMS [146] TheoreticalSpectrumGenerator class to characterise training signal composition and inform masking design. We generated b, y, and a ions (CID/HCD), c and z ions (ETD), at charge states 1^+^ through *z*_pre_ *−* 1^+^, including H_2_O and NH_3_ neutral losses and isotopic peaks up to M+4. Theoretical ions were matched to experimental peaks using a PPM-based greedy algorithm within a 10 ppm tolerance, with neutral losses and isotopic peaks matched conditionally on confirmed parent ion assignment. This analysis was conducted offline prior to model pretraining and does not use peptide annotations as supervision signals.

### Evaluation framework

All downstream evaluations except *de novo* sequencing used frozen mean-pooled peak-token embeddings without fine-tuning; the *de novo* sequencing benchmark fine-tunes the encoder weights (see below).

#### Evaluation data subsets

Supplementary Table S9 gives the corpus, split, and sample size for each evaluation. Unless otherwise stated, evaluations of the deployed InstaNovo-FM checkpoint use the held-out LCFM test split; the four-variant factorial ablation (section Factorial ablation of masking and attention bias) uses the LCFM validation split; and the signal-composition and theoretical-fragment analyses use HCFM. All LCFM splits are peptide-disjoint (80/10/10), so no peptide appears in more than one split. Spectra of all fragmentation types in the evaluation set are included; the taxonomy follows the b/y backbone series that dominates HCD/CID fragmentation, so the minority of ETD backbone ions (c/z) fall into the unannotated class. For the peak-level analyses, per-peak embeddings were extracted from the final encoder layer for 10,000 test spectra yielding *∼*411,000 individual peaks. Peaks were classified into four categories (b-ion, y-ion, precursor-related, unannotated) using theoretical fragment matching. A single-layer logistic regression classifier was trained on frozen peak embeddings with balanced class weights and early stopping (patience 10). Same-ion pairs (identical peptide, ion type, position, and charge state across independently acquired spectra) were compared against *m/z*-matched controls (distinct ions within 0.5 Da) and random controls. AUROC was computed for same-ion vs. *m/z*-matched discrimination. Within-spectrum cosine similarity was computed for isotope-parent pairs, neutral-loss-parent pairs, and random within-spectrum pairs.

### External benchmark implementations

We used InstaNovo v1.2, Casanovo, and XuanjiNovo as external benchmarks for the downstream linear probe and duplicate retrieval tasks. We used the publicly available InstaNovo v1.2 checkpoint, referred to as IN v1.2, Casanovo v5.0.0, and XuanjiNovo 100M MassNet checkpoints. At the time this work was written, this was the latest publicly available InstaNovo model. Casanovo embeddings and *de novo* predictions were produced with the casanovo package v5.1.2 loading the v5.0.0 release checkpoint (casanovo v5 0 0.ckpt). To extract embeddings from IN v1.2, we load the model checkpoint and pass the relevant data through the encoder component. Similarly, to obtain embeddings from Casanovo, we use the *casanovo* python package to load the model. We then pass the data through the encoder component and extract the embeddings. To extract embeddings from XuanjiNovo we load the spectrum encoder from the checkpoint and pass the data through. We note that XuanjiNovo feeds the precursor information directly into the encoder hence the precursor information is leaked into the embeddings. For this reason we exclude linear probe comparisons linked to precursor information (precursor mass, precursor *m/z*, precursor charge) as the model will be unfairly advantaged. For all embeddings we use the mean peaks embedding by averaging over the non-padding peak tokens. All encoders are probed on the same held-out LCFM spectra; this evaluation is in-distribution for InstaNovo-FM, but out-of-distribution for IN v1.2, Casanovo, and XuanjiNovo, which were pretrained on other corpora. We use these findings to demonstrate the depth of information encoded in the InstaNovo-FM embeddings, rather than as a head-to-head comparison of results. For the task of *de novo* sequencing, we benchmarked against the same checkpoints. The benchmark comparator models were run with a minimum peak-intensity threshold of 10^−6^, whereas InstaNovo-FM uses the 0.01 threshold at which it was trained; both thresholds are relative to the base-peak (maximum) intensity. Dropping peaks below 1% of the base peak removes only a tiny fraction of the peaks and does not appreciably shift the intensity distribution, so this difference does not materially affect the comparison. To implement the IN v1.2 and Casanovo models for *de novo* sequencing, we imported their respective code packages *instanovo* and *casanovo*, and ran their pipelines. To implement the XuanjiNovo model for *de novo* sequencing, we imported the code available on the MassNet-DDA GitHub and followed the standard evaluation pipeline documented on their repository.

### Linear probe and duplicate retrieval tasks

These probes underlie the uniform benchmarking reported in section Frozen InstaNovo-FM embeddings encode spectral and metadata structure learned without labels. In each case, a single linear classifier (or ridge regressor for continuous targets) was trained on the frozen embeddings across tasks spanning technical, physical, and biological dimensions of spectral information: fragmentation method and instrument identification, precursor charge, precursor *m/z* and mass, hydrophobicity, PTM presence and modification class, and peptide retrieval (Recall@1, mAP@20). Because the encoders differ in both capacity and training data, this is not a controlled comparison of architectures; holding the probe protocol uniform instead lets us read each embedding space for the depth of information it exposes. Embedding-extraction details, including the peak-intensity thresholds used per encoder, are given in the above section.

### Adapting InstaNovo-FM to *de novo* sequencing

To benchmark against IN 1.2, Casanovo, and XuanjiNovo on *de novo* sequencing, we adapted InstaNovo-FM. To do this, we removed the prediction heads and attached the InstaNovo model decoder layers, amino acid embedding layer, amino acid positional encoding layer, and amino acid prediction head.

To assess the impact of pretraining InstaNovo-FM, we ran three InstaNovo-FM *de novo* sequencing experiments with the foundation encoder being trained from scratch, left frozen, or fine-tuned. For the from scratch run, we trained the entire model from the first step, while for fine-tuning the decoder was trained from scratch and the encoder was trained with gradual unfreezing (see below for further details). For the frozen encoder run, we left the foundation model encoder weights frozen and only trained the added components. We trained all three models over 2.5 million steps, warming up over the first 100,000 steps to a learning rate of 5 *×* 10^−5^, using a batch size of 128. For the fine-tuning run of InstaNovo-FM, the peak encoder and encoder weights were unfrozen at step 100,000, while the other components (decoder, head, amino acid embedding, amino acid positional encoding) for *de novo* sequencing were trained from the start. We leave the InstaNovo-FM latent token frozen to preserve the information extracted from pretraining.

As shown in Supplementary Fig. S18, and further detailed below in a benchmarking experiment, we found the fine-tuned InstaNovo-FM to have the best performance of the three InstaNovo-FM variants, when taking peptide recall and convergence to steady state into account. Therefore, we use this variant for the *de novo* sequencing benchmark against IN v1.2, Casanovo and XuanjiNovo.

### Impact of training dataset and model size on *de novo* sequencing performance

To explore the impact of training dataset and model size on *de novo* sequencing performance, we trained two variants of the InstaNovo model, from scratch. To eliminate performance difference due to dataset, we trained on the same LCFM splits used to train InstaNovo-FM. For the first variant we used the publicly available model architecture, and for the second variant we used an adaption of the model with the encoder dimensions increased to match the sizes used for InstaNovo-FM, to investigate the effect of model size on performance. Hence, for the matched encoder size run, the number of encoder layers was increased from 9 to 12, the dimension of the feedforward layer was increased from 1024 to 3072, and the number of attention heads was decreased from 16 to 12.

We benchmarked these two InstaNovo variants against the three InstaNovo-FM variants detailed above, following the same training setup. To evaluate the *de novo* sequencing models we made use of the biological validation dataset featuring six smaller sets of data, as used in Fig. 5E. Note that we exclude the Immuno and Herceptin datasets used in the InstaNovo paper [28] due to the small amount of available data (*n <* 1000), since peptide recall would not be representative of model performance for these dataset sizes. For all models we performed decoding with beam search using *n_beams_* = 5.

The peptide recall results are displayed in Supplementary Fig. S18 and Supplementary Table S13. Pretraining combined with fine-tuning produces a *de novo* sequencing model that is competitive with a size-matched InstaNovo: the fine-tuned InstaNovo-FM matches or exceeds the size-matched baseline on four of the six datasets (Snake Venoms, *Hela QC*, TPL Antibodies and Wound Fluids) and trails it closely on GluC and S Brodae. The InstaNovo-FM variant trained from scratch performs almost identically (mean recall 0.556, versus 0.558 for the fine-tuned model), and the standard and size-matched InstaNovo baselines fall in the same band (0.552 and 0.554). Many of the differences among these four models are small and could be attributed to run variability. On this benchmark we therefore observe no measurable *de novo* advantage from foundational pretraining over training the same architecture directly, however there is a gain in training speed as the pretrained model was seen to converge to steady state faster. We see that the frozen InstaNovo-FM performs worse on all datasets, however the performance gap is modest given that the encoder is not adapted to the task, with representations being learnt through self-supervised masked *m/z* reconstruction. Across the six datasets the frozen model retains on average 85% of the fine-tuned recall (the mean of the six dataset peptide recall ratios of frozen to fine-tuned), and as much as 94% and 98% on GluC and S Brodae. The fact that so much sequencing performance is achieved without adapting the encoder shows that the self-supervised embeddings already encode a large portion of the information required for peptide identification. The largest performance gaps occur on the most out-of-distribution samples (Snake Venoms and Wound Fluids), where task-specific adaptation through fine-tuning contributes most.

### Factorial ablation of masking and attention bias

To isolate the contributions of masking strategy and pairwise attention bias (PA-bias) [57], we trained four variants of InstaNovo-FM on the LCFM corpus under identical conditions — 12-layer transformer, *∼*230,000 steps on *∼*117 million training spectra — differing only in masking strategy (signal-aware fragment masking, SA, which uses theoretical fragment annotations to mask complete annotated fragment groups, versus Thompson-span masking with isotope co-masking, TS) and the presence or absence of PA-bias (PA versus noPA): SA·noPA, SA·PA, TS·noPA, and TS·PA. Because all four variants share the LCFM corpus and every other hyperparameter, the differences reported in the main text reflect masking and attention-bias choices alone.

Each variant was evaluated on the full battery of frozen-embedding probes described above, which measure complementary facets of representation quality. Fragment-type and peak-type F1 measure how well a linear probe recovers ion class (b/y/neutral-loss/unannotated) from frozen embeddings; instrument macro F1 and PTM balanced accuracy measure technical and chemical decodability; precursor *m/z*/mass R² and spectrum confidence R² measure regression of continuous physical quantities. Recall@1 is retrieval fidelity — the fraction of query spectra whose nearest embedded neighbour is a true replicate of the same peptide. Cross-spectrum ion-identity AUROC measures whether the same fragment ion, observed in different spectra of the same peptide, embeds more similarly than a different ion at the same approximate *m/z*, testing that the model encodes ion identity rather than mere *m/z* position.

We additionally quantified attention specialisation on a per-head basis. A structurally-enriched (“structural”) head is a self-attention head whose attention between peaks in a known physical relationship — isotopic partners, complementary b/y ions, or neutral-loss pairs — exceeds a fixed multiple of a random- attention baseline, in which the same attention mass is redistributed uniformly over all peak pairs. The mean isotope-spacing enrichment reported for a variant is, averaged across its structural heads, the ratio of observed attention between peaks separated by one isotope spacing (1.003355*/z* Da) to the attention expected under that random baseline.

### Corpus-scale evaluation

To assess the effect of training-set scale and label confidence, we compared the deployed model — a 12-layer TS·noPA encoder trained for *∼*230,000 steps on the full LCFM corpus — against an MCFM baseline: a smaller 9-layer TS·noPA model trained for *∼*90,000 steps on the curated MCFM subset (*∼*10% of LCFM spectra). Both use identical Thompson-span masking with isotope co-masking and differ only in training corpus and capacity. MCFM is a strict subset of LCFM and the two tiers span essentially the same acquisition diversity, so the comparison varies training-set scale and label confidence while holding instrument and fragmentation breadth essentially fixed, rather than introducing a disjoint dataset. Confidence-based curation does itself shift the spectral distribution (MCFM is enriched for cleaner, higher-scoring spectra), so scale and label confidence covary here rather than being independently controlled. To remove the evaluation-set confound, both checkpoints were evaluated on the same held-out LCFM spectra under an identical frozen-probe protocol (Supplementary Note 6).

## Acknowledgements

K.K. is supported by a postdoctoral fellowship grant from the Independent Research Fund Denmark (grant no. 4257-00010B). We thank the EuroHPC JU for a resource allocation grant used in this work (EHPC- AI-2024A01-086), and the EuroHPC team for their feedback in deployment and training strategies in the MareNostrum v5 supercomputer environment. Models were trained on InstaDeep’s AIchor computing cluster (aichor.ai). We also thank Tine Claeys and Ralf Gabriels for their input on the format and structure of our dataset, and Duarte O. Carmo for discussions on model architecture and training.

## Author Contributions

K.K. and T.P.J. conceived the project. K.K. selected and annotated the PRIDE datasets. K.K. and J.D. designed and executed the data curation pipeline, and assembled the training corpus. M.N. and D.P. designed the model architecture and self-supervised training objective, and developed the model training infrastructure and conducted pretraining experiments. M.R., R.C., D.P. and J.D. contributed to data preprocessing, training, data analysis and results interpretation. R.C. and K.E. provided feedback on experiment design and model evaluation. M.N., R.C., D.P. and I.H. performed evaluation and downstream tasks. M.R., M.N., K.K., D.P., R.C., J.D. and I.H. drafted the manuscript with feedback from J.D., A.S., J.V.G and N.L.C. N.L.C., A.S., T.P.J., J.V.G. and K.K. supervised the project. All authors reviewed and approved the final manuscript.

## Declaration of Generative AI and AI-Assisted Technologies in the Writing Process

During the preparation of this work the authors used Claude Opus (4.6 and 4.8) in order to explore scientific ideas and directions, assist with software development, as well as improve the readability and language of the manuscript text. The authors reviewed and edited the content as needed, and take full responsibility for the content of the published article.

## Data Availability

All training data are publicly available through the PRIDE repository [15], with the exception of one dataset (MSV000090792) deposited in MassIVE [16]. Project accession numbers are listed in Supplementary Table S1. Training and evaluation data are publicly available on HuggingFace: https://huggingface.co/datasets/ InstaDeepAI/InstaNovo. Each of the three labelled confidence tiers is released in two forms: splits/ holds the quality-filtered, peptide-disjoint training, validation and test partitions the model was trained and evaluated on, and by project/ holds the tier before filtering and splitting, one directory per accession, from which alternative partitions can be derived. The unlabelled ACFM tier is approximately 5.7 TB and is not redistributed; because it comprises every MS/MS scan from the same raw files, it is reconstructible from the PRIDE accessions with the conversion pipeline deposited at Figshare, and is otherwise available from the authors on request. All results in this work were produced from release v0.1 of this dataset; passing revision=v0.1 to load dataset in the HuggingFace datasets library reproduces that exact snapshot. The same dataset also contains the central peptide registry of split assignments, so that the peptide-disjoint 80/10/10 partitions can be reproduced and extended with new data. The corpus was assembled in two stages, each released separately. The first converts vendor raw files into database search results. Its deposit at Figshare (DOI: 10.6084/m9.figshare.33368752) holds the dataset curation metadata, the protein sequence databases, the conversion and search scripts, and the FragPipe workflow configurations with a table mapping each to the datasets it was applied to. The second turns those search results into the trainable, peptide-disjoint splits released above — tier construction, quality filtering, splitting and shuffling — and is in the InstaNovo-FM code repository (see Code Availability). An interactive explorer for the frozen embedding space is hosted at https://instadeepai.github.io/InstaNovo-FM.

## Code Availability

The InstaNovo-FM foundational model repository, including training code, evaluation pipelines, the dataset-construction pipeline that builds the confidence tiers and peptide-disjoint splits, and Jupyter notebooks to reproduce the figures, is available at https://github.com/instadeepai/InstaNovo-FM. The InstaNovo code- base, including *de novo* sequencing inference and model evaluation utilities, is available at https://github.com/instadeepai/InstaNovo. Model weights are available from the Releases section of the InstaNovo-FM repository (https://github.com/instadeepai/InstaNovo-FM/releases). Each external benchmark model is run from its own container definition under docker/ in the InstaNovo-FM repository: Casanovo from package v5.1.2 on python:3.11-slim, loading the v5.0.0 release checkpoint, and XuanjiNovo built from the MassNet-DDA source at commit 84105ec on pytorch/pytorch:2.1.0-cuda12.1-cudnn8-runtime.

## Supplementary Information

InstaNovo-FM: A Self-Supervised Foundation Model for Proteomics Tandem Mass Spectrometry

### Supplementary Notes

#### Note 1: Model architecture details

InstaNovo-FM builds upon the spectrum encoder introduced in InstaNovo [28], extending it to large-scale, task-agnostic pretraining on heterogeneous proteomics datasets. The full architecture is described in Fig. S4. **Peak encoder.** Each (*m/z,* intensity) pair is embedded using a multi-scale sinusoidal encoder with learnable frequencies [54]. The *m/z* value is projected onto *K* = *d/*2 = 384 frequencies and expanded in both quadratures,

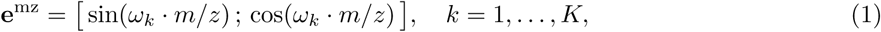

giving a vector of dimension *d* = 768. The frequencies *ω_k_* are initialised log-spaced, corresponding to periods of approximately 2.5–25 Da, and are trained with the rest of the model. This vector is passed through a two-layer MLP with ReLU activations, concatenated with the peak’s scalar intensity, and projected through a second two-layer MLP back to dimension *d* = 768.

##### Positional encoding

No positional encoding is applied to the peak sequence: peaks are processed as an unordered set, and neither rotary position embeddings nor additive positional signals are used. The only positional information available for a masked peak is a Gaussian-blurred *m/z* value (*σ* = 10 Da) combined with the peak’s original visible intensity, which supplies approximate spatial identity without revealing the *m/z* reconstruction target (Methods, section Thompson masking with isotope co-masking); no learnable mask token is used.

#### Note 2: Failure of hyperspherical uniformity at high dimensionality

Hyperspherical uniformity loss encourages embeddings to spread uniformly on the unit hypersphere:

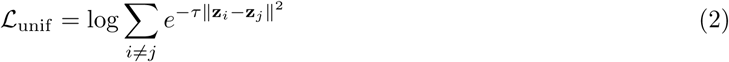

For a batch of *B* = 1024 random unit vectors in R^768^, pairwise cosine similarities concentrate near zero regardless of the embedding’s effective dimensionality *d*_eff_ (participation ratio). Monte Carlo simulation shows the loss difference between *d*_eff_ = 768 (full rank) and *d*_eff_ = 17 (severely collapsed) is only *≈* 0.008 — invisible to the optimizer at a loss scale of *∼* 3.0. This was confirmed empirically: three reproducibility runs each showed *L*_unif_ *≈* 0.06934 *≈* log(1024) = log *B*, the theoretical minimum, throughout training. The run achieving the lowest training loss (DNS-2984, loss 3.003) also produced the worst participation ratio (*d*_eff_ = 17.19 vs. 33.03 and 27.52 for the other runs) and worst retrieval (Recall@1 = 0.589 vs. 0.673 and 0.657), demonstrating that hyperspherical uniformity is mathematically unable to prevent dimensional collapse in high-dimensional embedding spaces.

#### Note 3: Binning strategy evaluation

##### Candidates

Twelve binnings were evaluated across three families: four fixed-width grids (0.01, 0.02, 0.05 and 0.1 Da), three fixed-ppm grids (10, 20 and 50 ppm), and five adaptive hyperbolic schedules 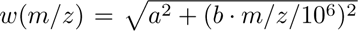, which behave like a fixed-width grid of width *a* at low *m/z* and like a fixed-ppm grid of slope *b* at high *m/z*. Per-strategy results are given in Supplementary Table S2.

##### Metrics

Each was scored on two quantities that trade off against bin width, measured over 200,000 LCFM spectra (27 million peaks). The jump rate is the fraction of peaks of the same chemical identity that land in different bins across spectra of the same peptide; it is the label noise the classification head inherits. The collision rate is the fraction of same-spectrum peaks sharing a bin; it is the ambiguity in the target. Narrow bins lower collisions but raise jumps, and wide bins do the reverse. Eight of the twelve strategies are Pareto-optimal on the two axes, and the combined score ^1^ (jump + collision) varies by less than half a percentage point across the four coarsest of them. The choice was therefore made on physical grounds. **Fixed-ppm.** Because width scales with *m/z*, these grids become unresolvably fine at the low end. At 20 ppm the width at *m/z* 100 is 0.002 Da, so the Gln/Lys difference of 0.036 Da is spread over 18 bins and the singly-charged isotope spacing over 502, spending head capacity on distinctions the training signal cannot support. **Adaptive.** Within the proteomic mass range the best-performing schedule is almost identical to a 0.2 Da fixed grid, varying only from 0.200 Da at *m/z* 100 to 0.204 Da at *m/z* 2000, so it adds vocabulary-shape complexity to the hierarchical head without changing behaviour. **Selected.** The deployed model uses a fixed 0.2 Da grid of 12,250 bins (jump 3.00%, collision 6.40%). This width is the one whose bin edges match the instrument: the fitted P95 envelope of fragment-ion mass error on LCFM is approximately *m/z*-independent at *≈* 0.09 Da, so a 0.2 Da bin holds a *≈* 1.1*×* margin against that tail, where 0.1 Da sits at *≈* 0.56*×* and finer widths fall below 0.3*×*. Going coarser than 0.2 Da starts to merge chemistry the model should resolve: single-residue differences, neutral losses, and isotope spacings at higher charge.

#### Note 4: Theoretical fragment ion annotation methodology

Fragment ion matching was performed in two stages. In stage 1, base fragment ions were matched to experimental peaks using a PPM-based greedy algorithm within 10 ppm tolerance, assigning peaks one-to-one in order of decreasing theoretical intensity. In stage 2, neutral losses (H_2_O, *−*18.011 Da; NH_3_, *−*17.027 Da; H_3_PO_4_, *−*97.977 Da) and isotopic peaks (spacing 1.003355*/z* Da) were generated and matched only for base ions confirmed in stage 1. This conditional approach reduces false positives by exploiting the physical dependence between parent ions and their associated neutral losses and isotope series.

#### Note 5: Integrated-gradients attribution and off-database peak matching

##### Integrated-gradients attribution

For a spectrum passing quality gates (*≥*33% backbone cleavage coverage, *≥*7 distinct fragment groups), we mask up to three fragment groups per spectrum and, for each, attribute the model’s reconstruction prediction back to the visible peaks using integrated gradients [56] (Captum [147], 50 integration steps). The baseline is the same spectrum with all peak intensities set to zero while *m/z* values are preserved, so attribution measures the contribution of each peak’s intensity to the masked prediction; the completeness (convergence) delta is recorded for every attribution. This yields a per-peak influence score and, per masked group, the rank of the ion-ladder neighbour (a peak displaced from the masked ion by a single amino-acid residue mass) among the most-attributed peaks. Across 413 evaluation spectra this gives 1,239 masked-group predictions.

##### Attribution concentration

Two summaries describe how attribution is distributed over the peaks of a spectrum. The first is the Pearson correlation between a peak’s attribution and its relative intensity, computed across all visible peaks and pooled over masked groups; a high value would indicate that attribution simply tracks peak height. The second is the Lorenz curve and its associated Gini coefficient, borrowed from the measurement of inequality. Peaks are ranked from least to most attributed and the cumulative share of total attribution is plotted against the cumulative share of peaks: a model spreading attribution evenly over all peaks traces the diagonal, whereas one concentrating it on a few peaks bows below. The Gini coefficient is twice the area between the curve and the diagonal, so 0 denotes perfectly even attribution and 1 denotes all attribution falling on a single peak.

##### Off-database peak matching

The standard annotation “database” comprises the backbone b/y ions and their isotopes and neutral losses (Note 4); peaks not assigned there are unannotated. For each peptide we generate theoretical *m/z* values for additional, chemically defined off-database species — internal fragments (contiguous subsequences up to six residues), side-chain d- and w-ions, side-chain neutral losses, and combined precursor losses — and record an unannotated peak as a *match* when its observed *m/z* lies within 10 ppm (with a 0.5 Da coarse gate) of a theoretical off-database *m/z*. The per-category hit rate is the fraction of unannotated peaks with at least one such match (4,806 of 42,949; Table S7).

##### Null models

To establish that matches are not coincidental we compare the observed match rate against three nulls: a scramble null that shuffles the peptide’s residue order and rebuilds the fragment library from the scrambled sequence, leaving the observed spectrum unchanged (5 trials), an *m/z*-shift null that rigidly displaces the spectrum by non-physical offsets (3.7, 7.3, 11.1 Da), and, because shuffling a sequence preserves its amino-acid composition, an absent-residue control that matches immonium-related ions built only from amino acids not present in the peptide. Enrichment is the observed match rate divided by the null match rate (2.1-fold over scramble, 18-fold over *m/z*-shift); no enrichment was detected against the absent-residue control (0.39-fold, i.e. depleted).

##### Confidence

Per-peak joint confidence is the product of the model’s predicted probabilities for the correct group bin and the correct offset bin, *P* (group) *× P* (offset), from the hierarchical reconstruction head.

#### Note 6: Linear-probe evaluation

##### Setup

All probes are trained on frozen mean-pooled peak-token embeddings; the encoder is never fine-tuned. Features are standardised (zero mean, unit variance) using statistics fit on the training split only. Probes use the dataset’s predefined train/validation/test splits (100,000 / 10,000 / 10,000 spectra) with no peptide sequence shared across splits and per-project sample capping (maximum 15% of any split from a single project), with splits not being project-disjoint. Spectra whose target value is missing are dropped per probe, so the number of scored test spectra is at most 10,000 and varies by target (10,000 for precursor charge, 8,967 for fragmentation method). For reference, a majority-class predictor on the same test splits reaches macro-F1 0.171 for fragmentation method (4 classes), 0.104 for precursor charge (7), 0.025 for instrument (13), 0.082 for modification class (7) and 0.417 for PTM presence (2), with macro-AUROC 0.500 in every case; a majority-class macro-F1 cannot exceed 1/K for K scored classes. For the regression targets a mean predictor gives *R*^2^ = 0 by construction, so the reported *R*^2^ values are already stated relative to that baseline. Classification targets (fragmentation method, instrument, precursor charge, PTM presence) are reported as macro-F1 / balanced accuracy, macro-averaged over the classes present in the test split: four fragmentation methods (CID, ETD, HCD, HCID) and seven charge states. A fifth fragmentation label (spectra with no recorded fragmentation method) and a rarer high charge state occur in the probe’s training split but have no test support, so they cannot be scored; including them would make the denominator depend on whether a given model happens to predict them. Instrument (13-class) and PTM-presence metrics are unaffected by this convention. And regression targets (precursor *m/z*, precursor mass, hydrophobicity, spectrum confidence) as *R*^2^. Macro-F1 is used for the classification targets because they are heavily class-imbalanced: it weights every class equally, so strong performance on a dominant class cannot mask poor performance on rare ones. It is averaged over the classes with support in the test split.

##### Models

Classification uses multinomial L2-regularised logistic regression with class-balanced weighting; regression uses Ridge regression. The regularisation strength (*C* for logistic regression, *α* for Ridge) is selected on the validation split from a logarithmic grid.

##### Implementation and reproducibility

Probes are fit with cuML (RAPIDS) on GPU; logistic regression uses the L-BFGS solver with max iter = 5000. We pin the cuML version, because cuML and scikit-learn differ in their solver and in how the regularisation strength is scaled, so the same nominal *C*/*α* grid yields different effective regularisation and different probe scores between the two backends; fixing the backend keeps probe comparisons across models internally consistent. Because the embeddings are low-rank relative to their nominal dimension, the logistic-regression objective is near-degenerate (the L-BFGS line search converges into a flat basin); probe scores should therefore be read as well-regularised lower bounds on linear decodability rather than exact optima.

#### Linear-probe evaluation on *Hela qc*

To eliminate bias due to dataset distribution alignment, we perform additional evaluations on a downstream biological dataset, *Hela qc*. This dataset does not contain overlapping spectra with LCFM, the training dataset of the InstaNovo-FM model. We split the dataset (*n* = 17683) into 80%-10%-10% subsets, ensuring no peptide sequence overlap between splits. We use these splits to train, validate and test the linear probe models, using the methods detailed above. The results are summarised in Supplementary Table S11. We see that IN v1.2 performs best on the precursor metrics (precursor charge, precursor *m/z*, precursor mass), while XuanjiNovo leads on the remaining three metrics (PTM presence, hydrophobicity, modification class). InstaNovo-FM closely follows the leading precursor metrics, trailing IN v1.2 by only 0.2% on precursor mass, 0.5% on precursor *m/z*, and 3.3% on precursor charge, while outperforming Casanovo on all three. Casanovo pulls ahead of InstaNovo-FM on the remaining peptide-property metrics. This experiment demonstrates the importance of dataset distribution - where InstaNovo-FM previously lead on the precursor metrics in Supplementary Table S10, we now see IN v1.2 pulls ahead, although it is still close. The *Hela qc* dataset did not contain search instrument or fragmentation type metadata and so we could not perform linear probes on those metrics.

## Supplementary Figures

**Figure S1:**
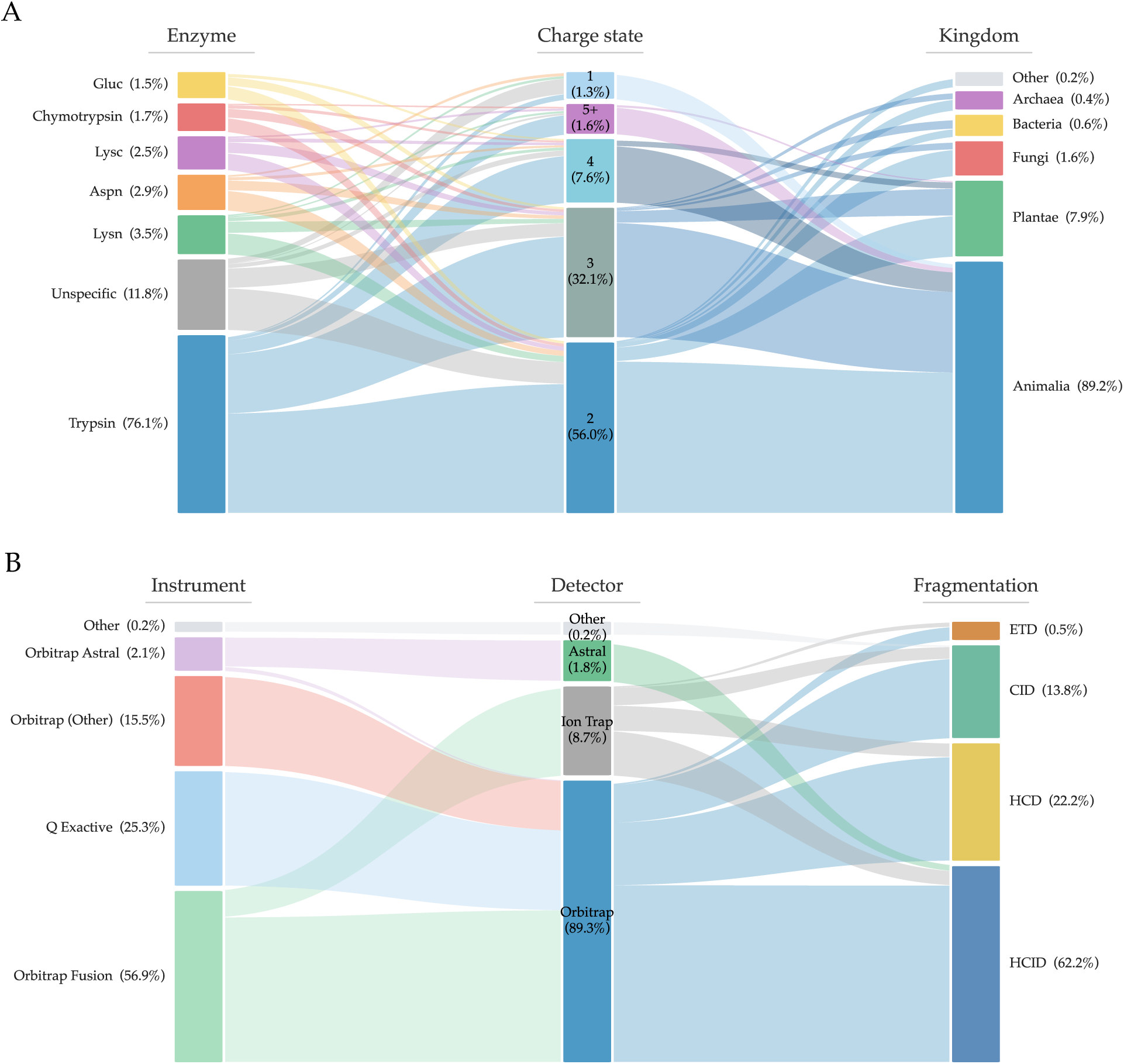
Corpus composition. **A.** Dataset biological characteristic composition as an enzyme *→* precursor charge state *→* taxonomic kingdom Sankey diagram. **B.** Technical characteristic composition as an instrument *→* detector *→* fragmentation-method Sankey diagram, showing the dominance of trypsin/Orbitrap/HCD alongside substantial long-tail diversity.

**Figure S2:**
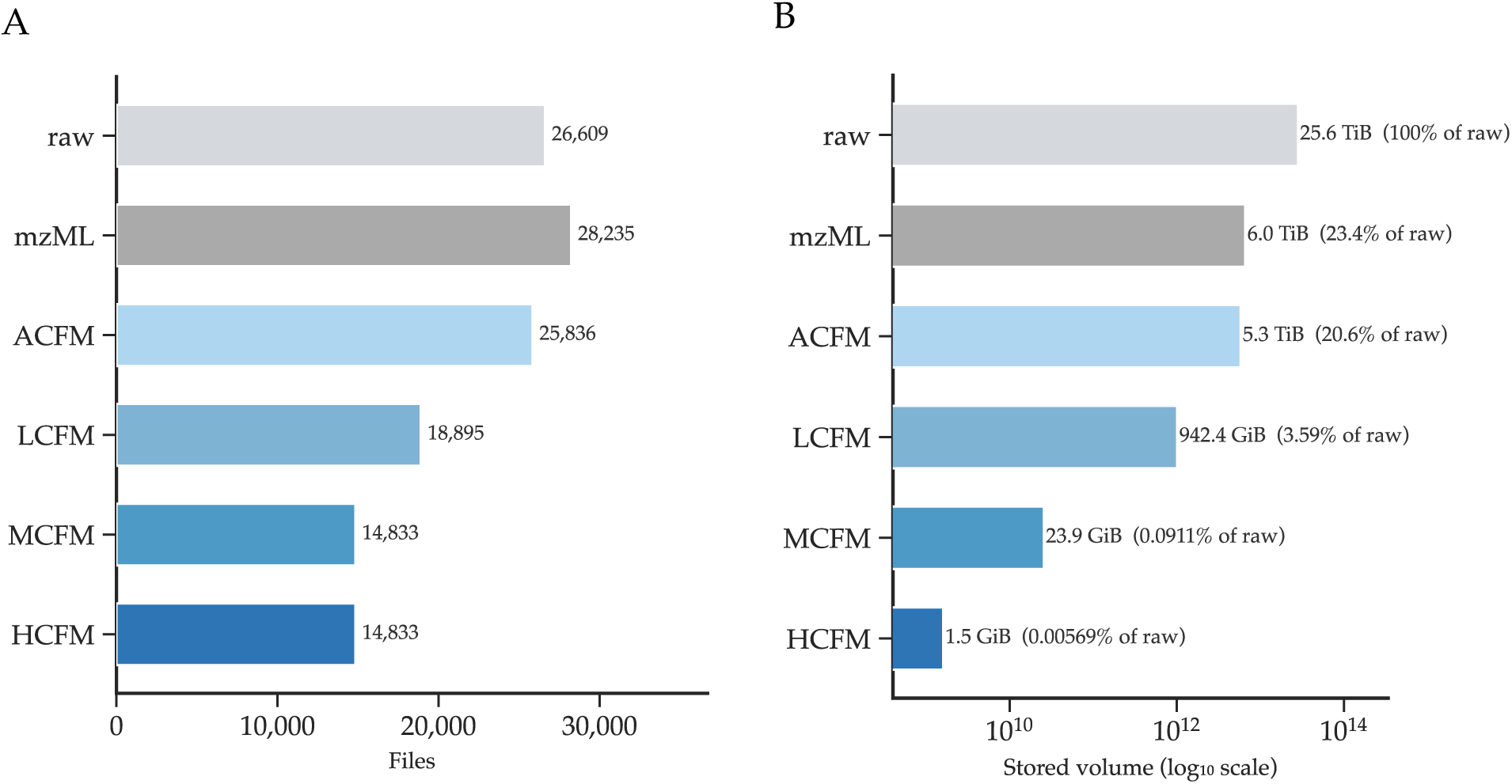
Data-volume funnel of the reprocessing pipeline. **A.** File counts and **B,** stored volume at each processing stage (raw *→* mzML *→* ACFM *→* LCFM *→* MCFM *→* HCFM), illustrating the compression achieved by the unified reprocessing and confidence-tiering pipeline.

**Figure S3:**
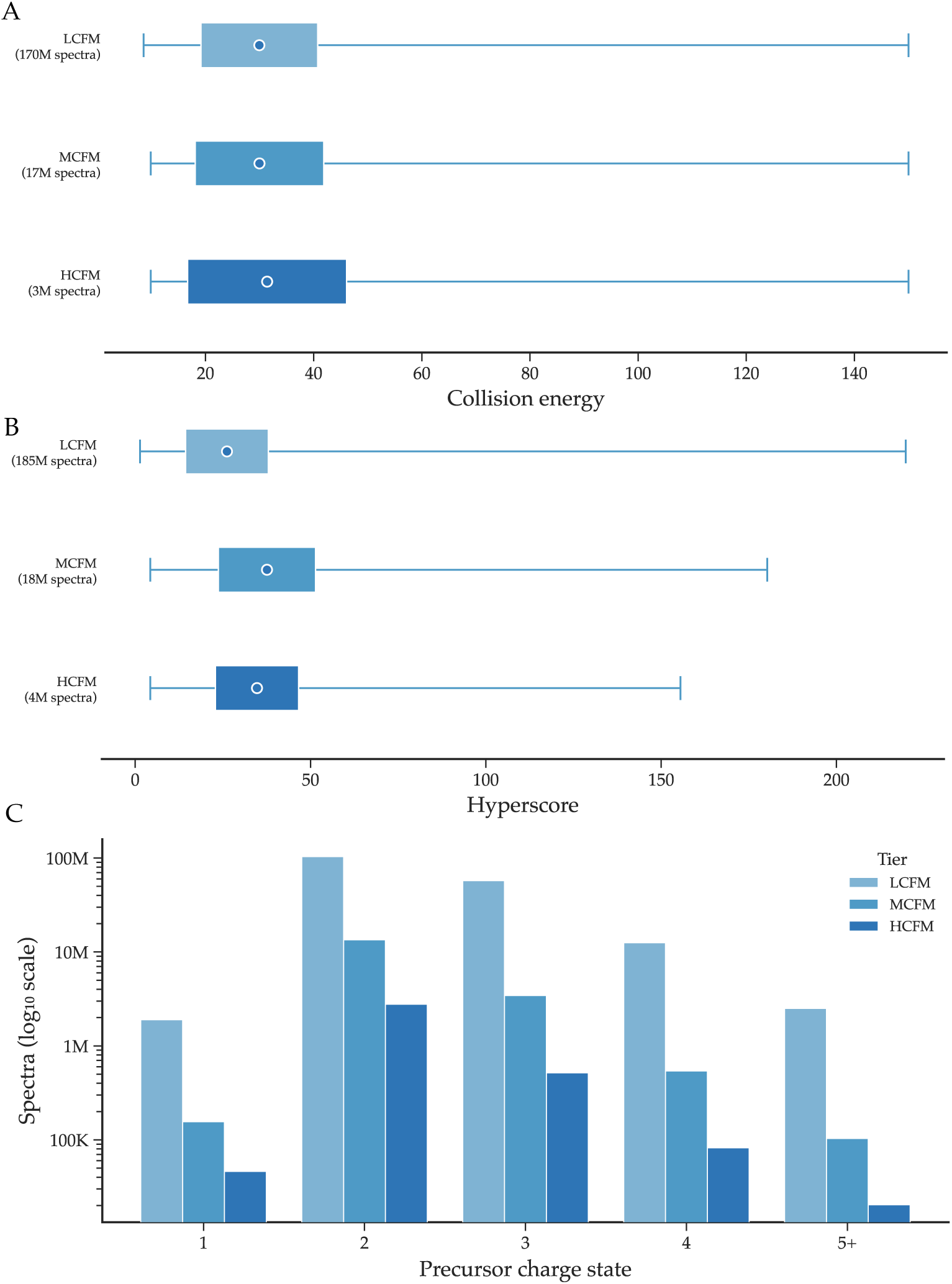
Acquisition-parameter distributions across confidence tiers. Distributions of **A,** collision energy, **B,** hyperscore, and **C,** precursor charge state for the LCFM, MCFM and HCFM tiers, showing that higher-confidence tiers preserve the overall parameter ranges. Extended numeric summaries across tiers are provided in Supplementary Table S6.

**Figure S4:**
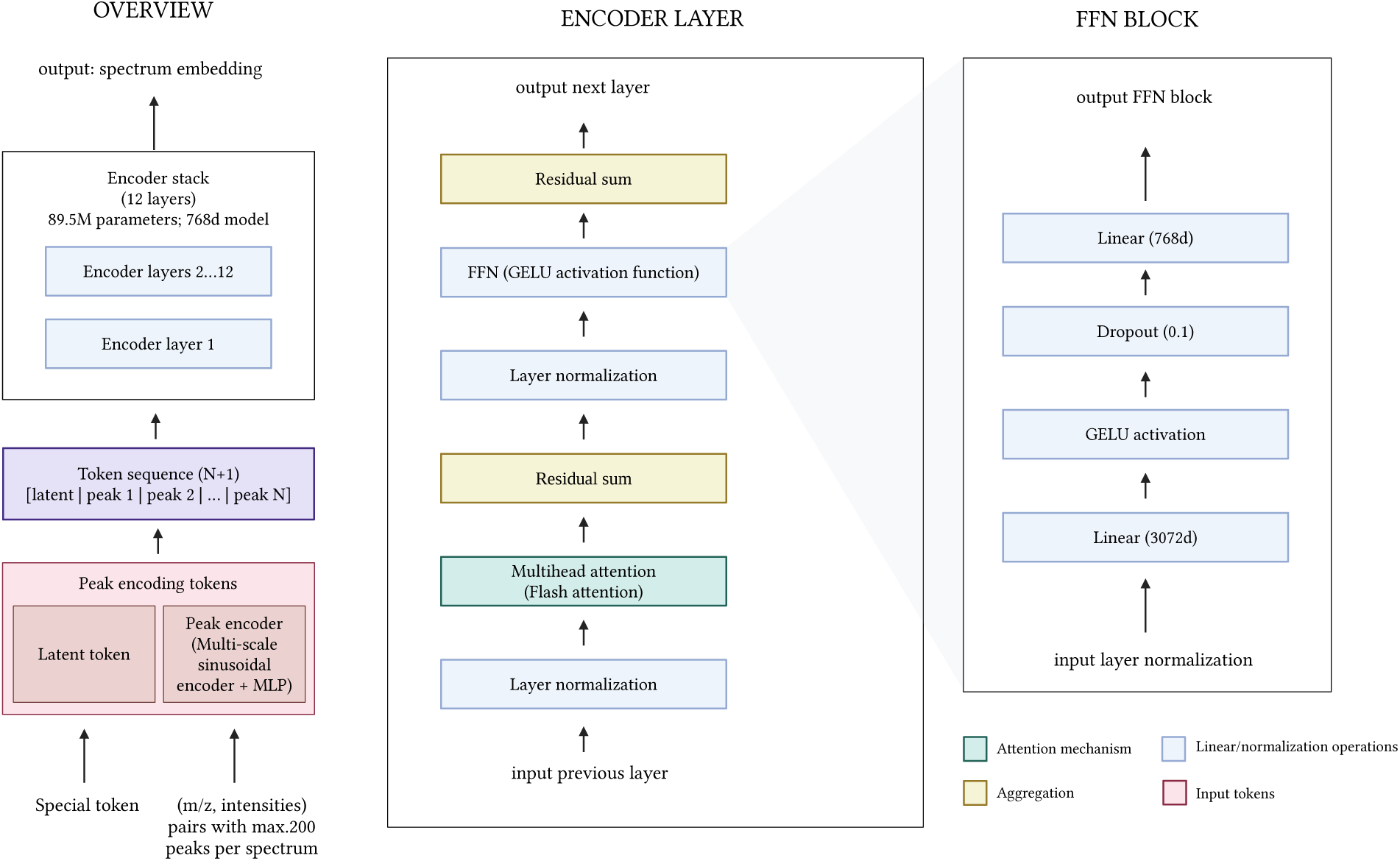
InstaNovo-FM architecture overview. **A.** End-to-end schematic: (*m/z,* intensity) peak sequences are embedded by a multi-scale sinusoidal peak encoder, prepended with a [CLS] latent token, and passed through the 12-layer transformer encoder (*≈*89.5M parameters, *d* = 768); the spectrum embedding is the mean of the final-layer peak-token states. **B.** Encoder-layer detail (pre-norm, multi-head attention with a Flash backend and no positional encoding, GELU feedforward, residual sums). **C**. Feedforward block detail.

**Figure S5:**
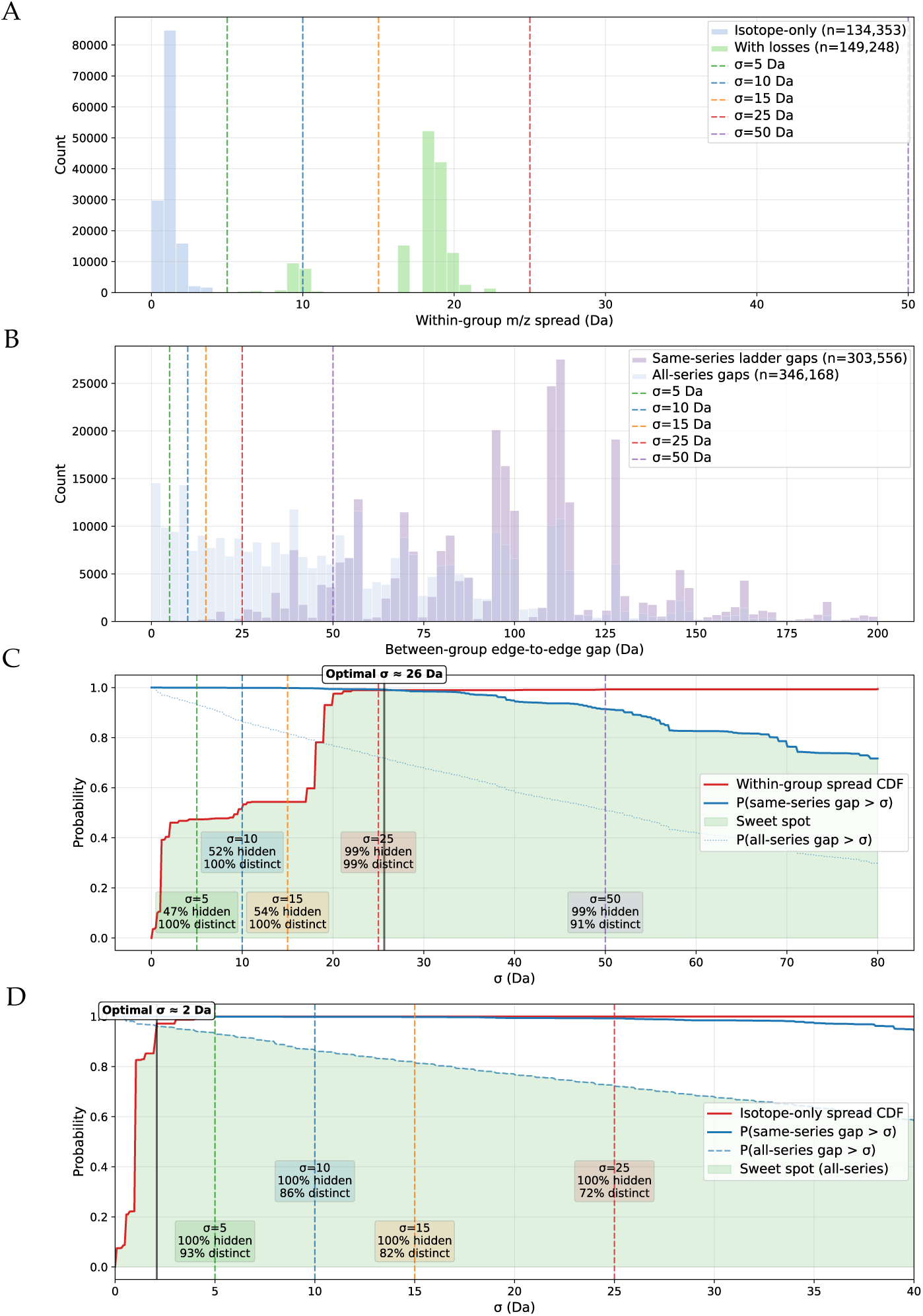
Calibration of the masked-peak blur width. *σ***. A.** Within-group *m/z* spread for isotope-only groups (*n* = 134,353) and groups that also include neutral losses (*n* = 149,248), with candidate *σ* thresholds (5–50 Da). **B.** Between-group edge-to-edge gaps for same-series ladder gaps (*n* = 303,556) and all-series gaps (*n* = 346,168). **C.** With-loss regime: the cumulative distribution function (CDF) of within-group *m/z* spread, i.e. the fraction of fragment groups lying entirely within *σ* and therefore fully co-masked, plotted against *P* (gap *> σ*); the “sweet spot” (all isotope/loss peaks co-masked while fragment groups remain separable) is maximised at *σ ≈* 26 Da. **D.** Isotope-only regime: the same trade-off, with dashed lines marking candidate widths and the fraction of groups fully blurred (“hidden”) and of between-group gaps still resolvable (“distinct”) at each. The deployed *σ* = 10 Da hides every isotope-only group while leaving 86% of gaps distinct, and hides 52% of loss-bearing groups with all gaps distinct; it is therefore the single width satisfying both regimes, where the per-regime optima marked on the panels each maximise one. Optimal for this regime alone is *σ ≈* 2 Da.

**Figure S6:**
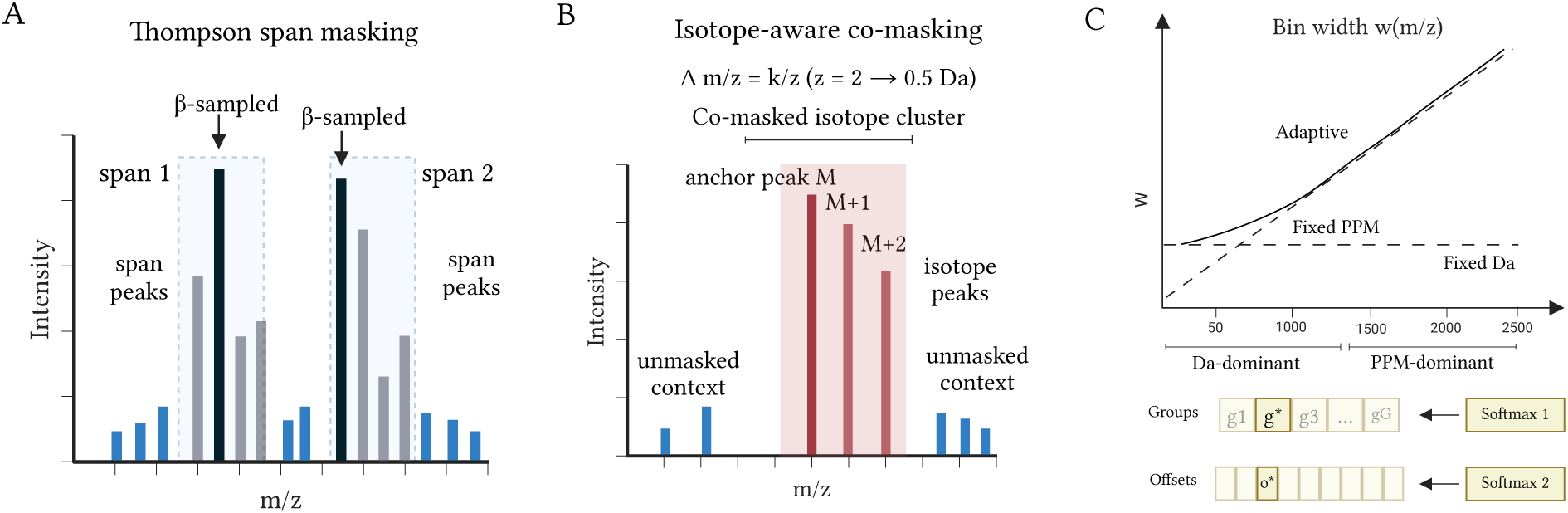
Self-supervised pretraining objective. **A.** Thompson-span masking: high-intensity anchor peaks are *β*-sampled (intensity-weighted) and expanded to contiguous *m/z*-ordered spans of peaks. **B.** Isotope- aware co-masking: each anchor peak *M* is co-masked together with its isotope cluster (*M* +1, *M* +2; spacing Δ*m/z* = *k/z*, where *k* = 1.003355 Da is the ^13^C–^12^C mass difference and *z* the fragment charge), keeping surrounding context unmasked. **C.** Adaptive bin-width schedule *w*(*m/z*) interpolating a Da-dominant and a PPM-dominant regime, and the hierarchical reconstruction head (group softmax followed by a within-group offset softmax).

**Figure S7:**
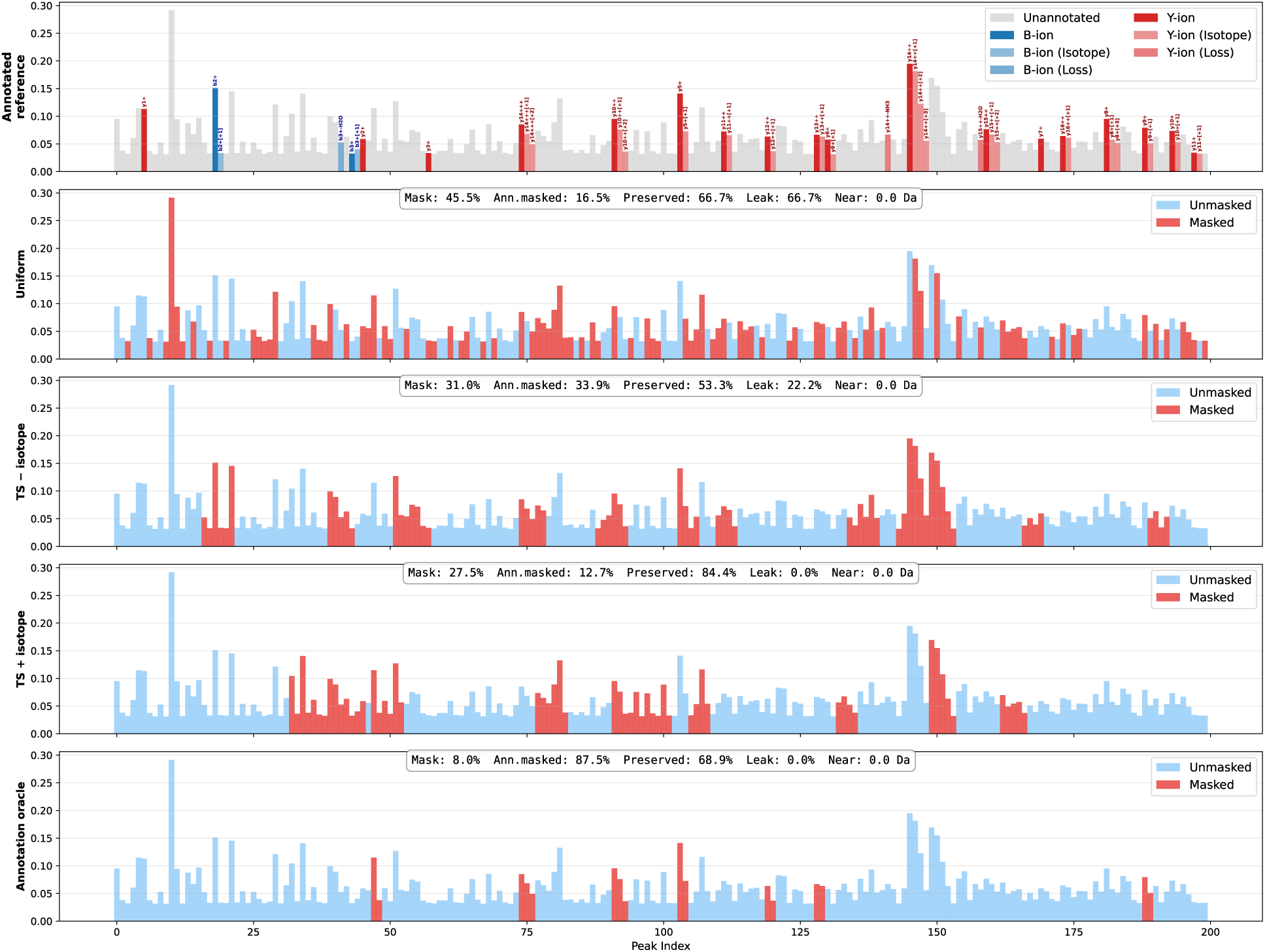
Masking strategies on a representative spectrum. Masking applied to one representative LCFM spectrum (VSAPGVLTAQDR). Top: annotated reference spectrum. Rows: masked peaks (red) under uniform masking, Thompson-span without isotope co-masking, Thompson-span with isotope co-masking, and the annotation-driven oracle, each annotated with per-strategy mask / annotated-masked / preserved / leakage rates. Aggregate metrics across all five strategies over 200,000 spectra are given in Supplementary Table S3.

**Figure S8:**
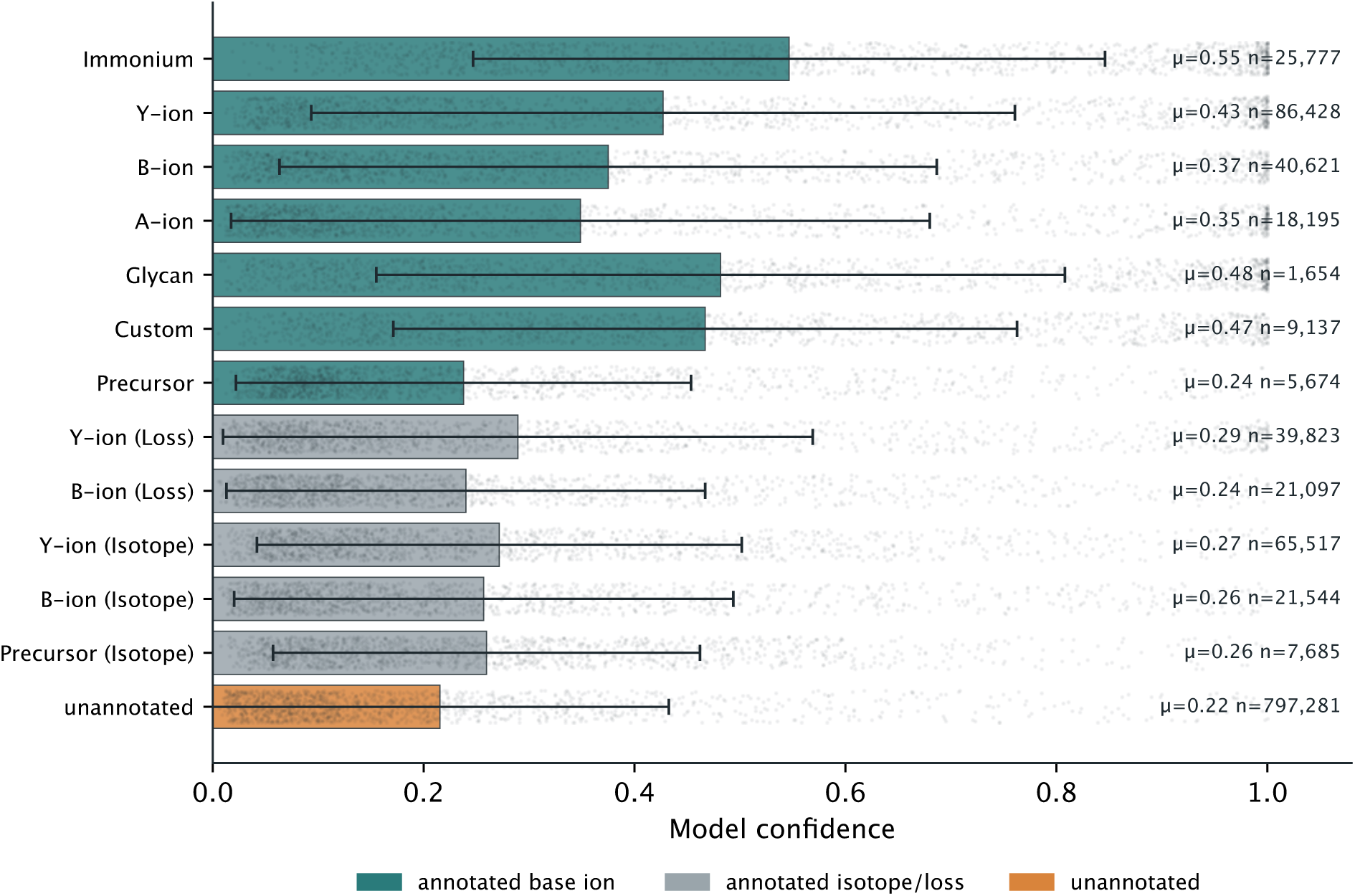
Per-peak model confidence by ion type. Per-peak reconstruction confidence by ion type on the LCFM test split (bars = mean, whiskers *±*1 SD, points = individual peaks, jittered and subsampled to *≤*1,500 per type). Annotated fragment ions (immonium, *y*, *b*) carry systematically higher confidence than unannotated peaks (annotated-vs-unannotated AUROC 0.640).

**Figure S9:**
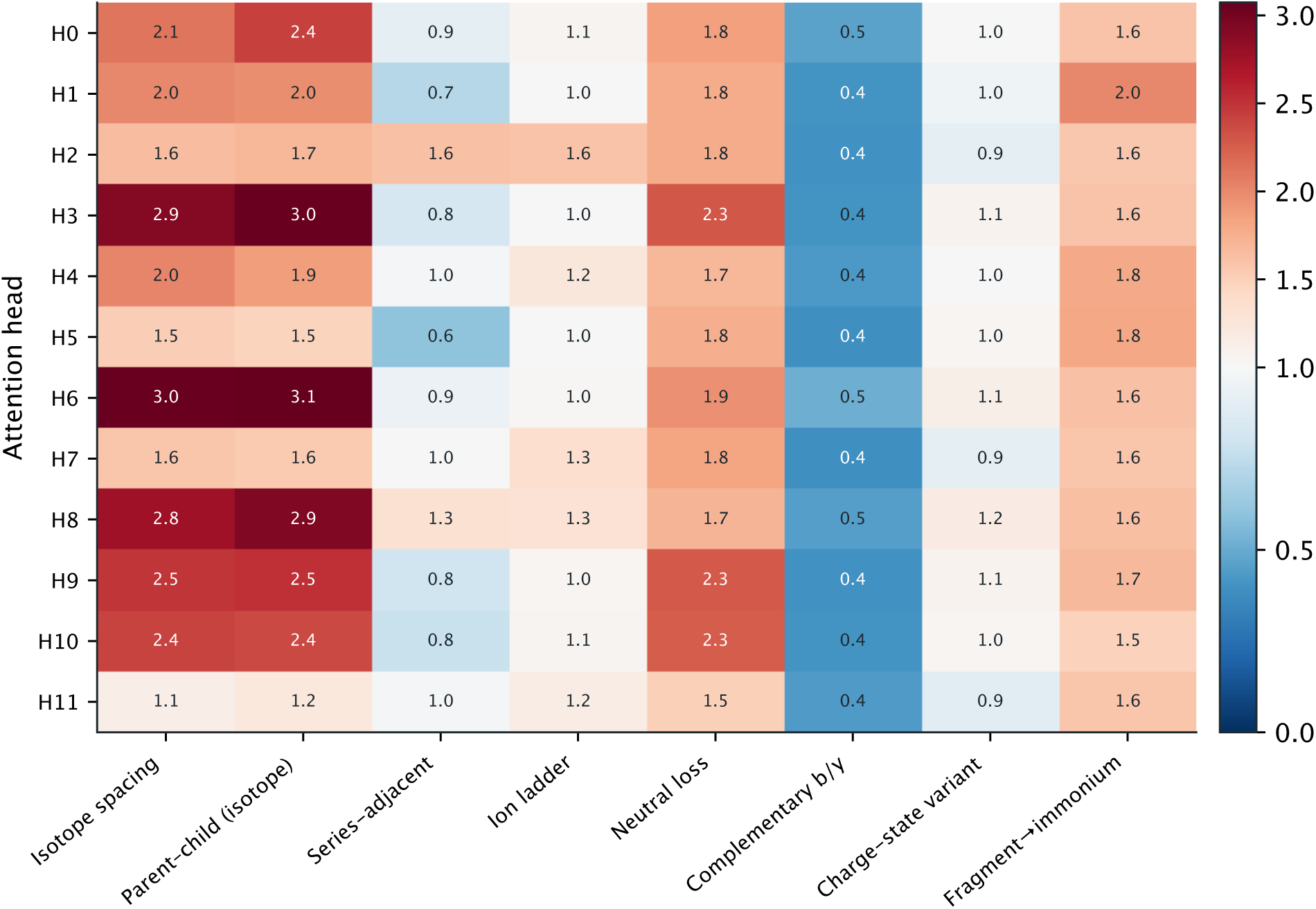
Per-head self-attention enrichment by physical relation. Per-head attention enrichment for each physical peak relation relative to a random peak-pair baseline (1.0) on the LCFM test split. 9 of 12 heads are structurally enriched; isotope-spacing and parent–child (isotope) relations dominate (up to *∼*3*×*), while complementary *b*/*y* attention stays at or below baseline.

**Figure S10:**
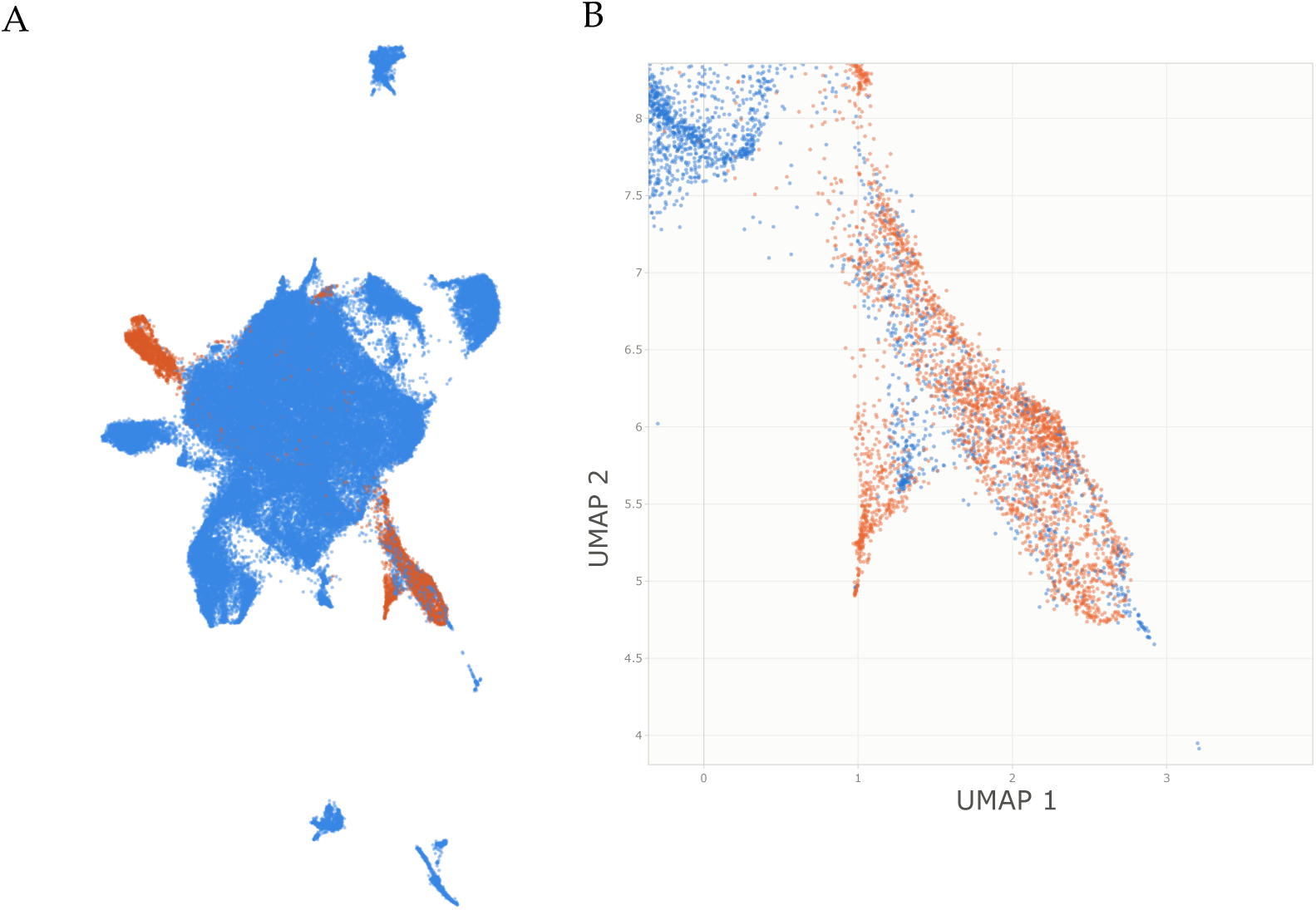
Spectrum-level embedding UMAP across DDA and DIA regimes. UMAP projections of InstaNovo-FM [CLS] spectrum embeddings on the 100k evaluation corpus coloured by acquisition mode (DDA vs. DIA). **A.** Full corpus. **B.** Zoomed subregion, showing that DDA and DIA spectra of the same peptides interleave rather than forming acquisition-driven clusters.

**Figure S11:**
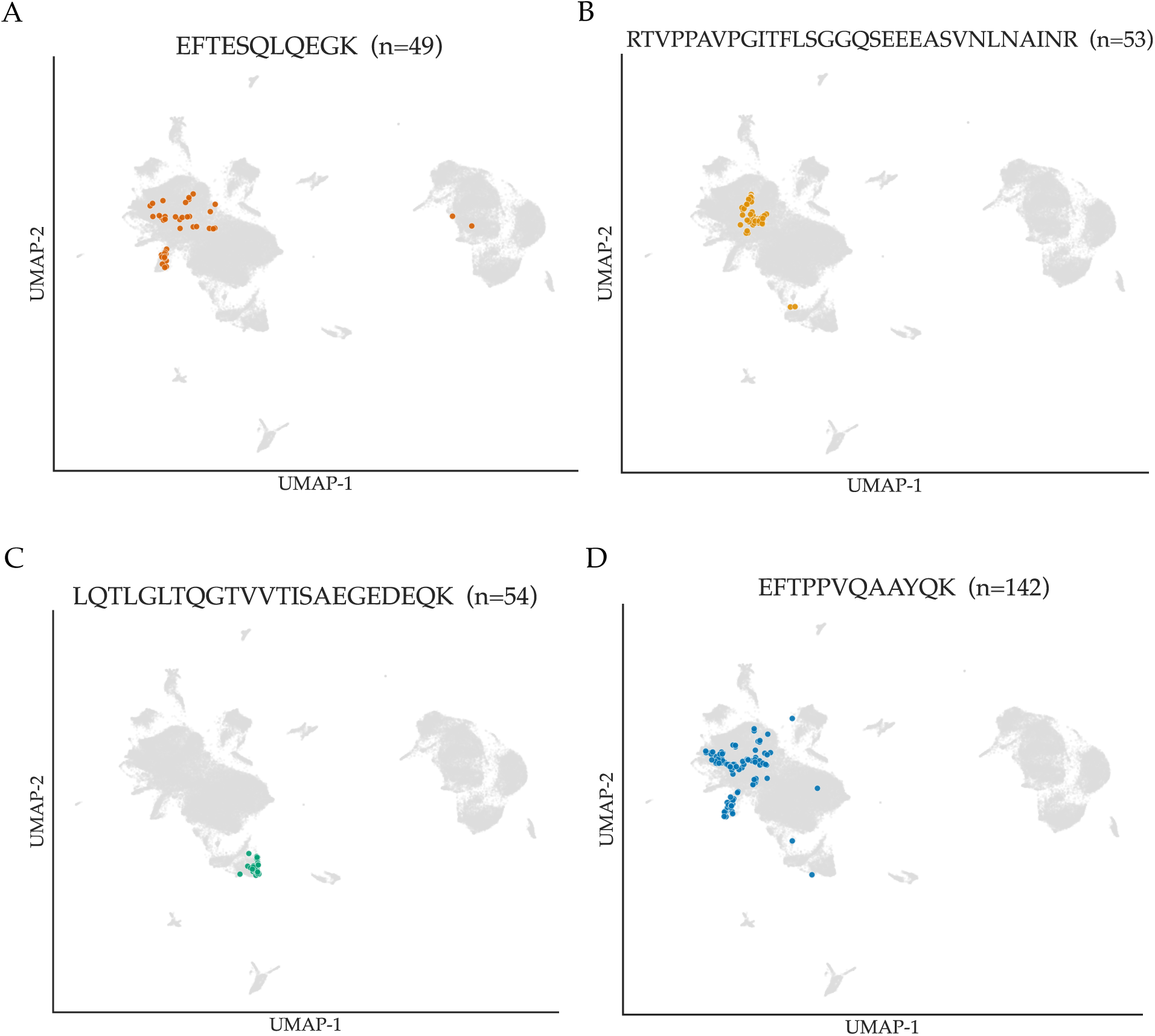
Query-peptide embedding projection for duplicate retrieval. UMAP projection of individual query peptides (coloured points) onto the InstaNovo-FM spectrum-embedding background (grey), for four example peptides of increasing occurrence count. Repeated observations of the same peptide land in a tight neighbourhood, illustrating the nearest-neighbour structure that underlies duplicate retrieval (benchmark values in Supplementary Table S10).

**Figure S12:**
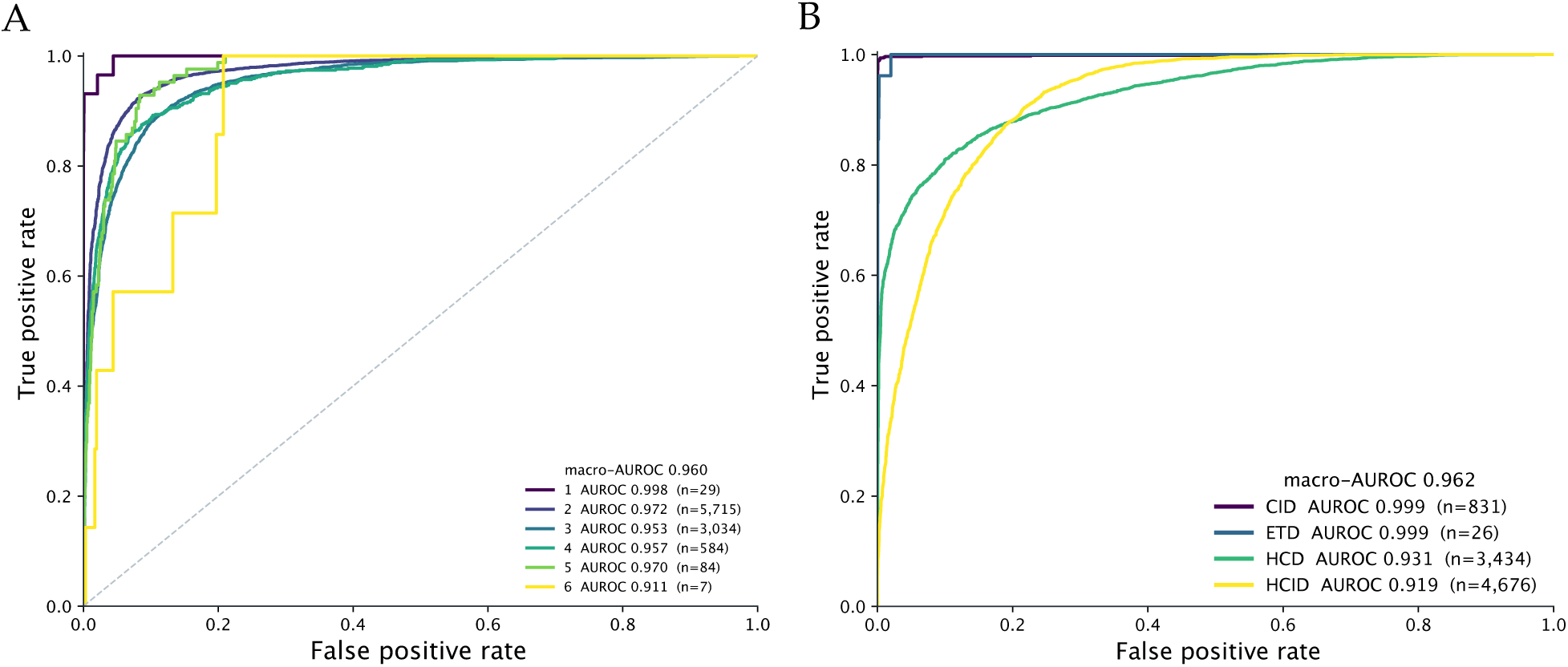
Per-class receiver operating characteristic (ROC) curves for frozen linear probes. One-vs-rest (OvR) ROC curves evaluated on frozen mean-pooled peak-token embeddings from the held-out LCFM test split. **A.** Precursor charge state (macro-AUROC 0.956). **B.** Fragmentation method (CID, ETD, HCD, HCID; macro-AUROC 0.962). Extended classification and regression benchmarks across architectures are detailed in Supplementary Table S10.

**Figure S13:**
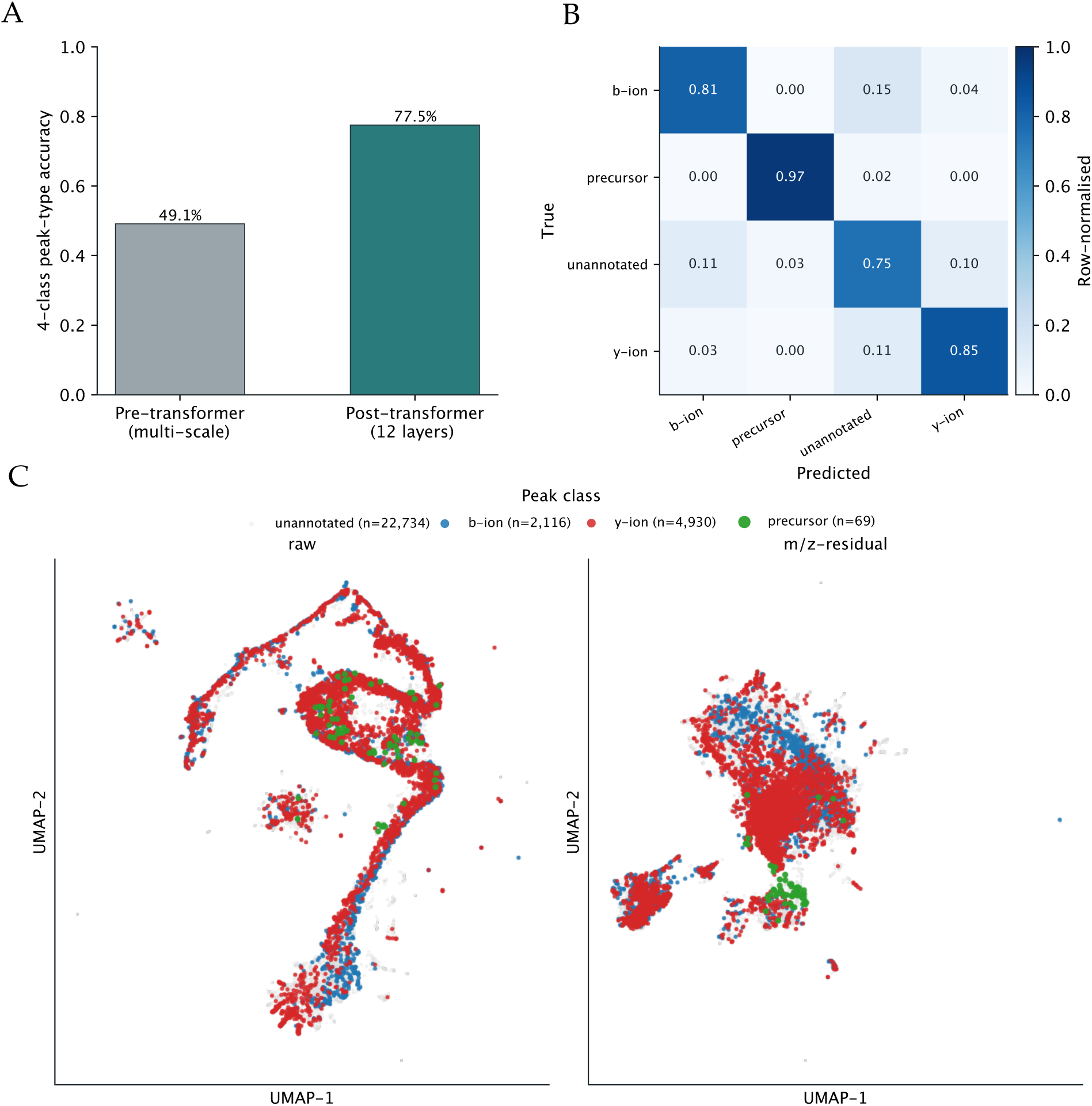
Peak-type structure emerges through the transformer. **A.** Four-class peak-type decoding accuracy from frozen peak-token embeddings: 49.1% before the transformer to 77.5% after it. **B.** Row-normalised confusion matrix of the post-transformer probe. **C.** UMAP of peak-token embeddings coloured by ion class (raw and *m/z*-residual); *b*/*y*-ions separate cleanly from unannotated peaks.

**Figure S14:**
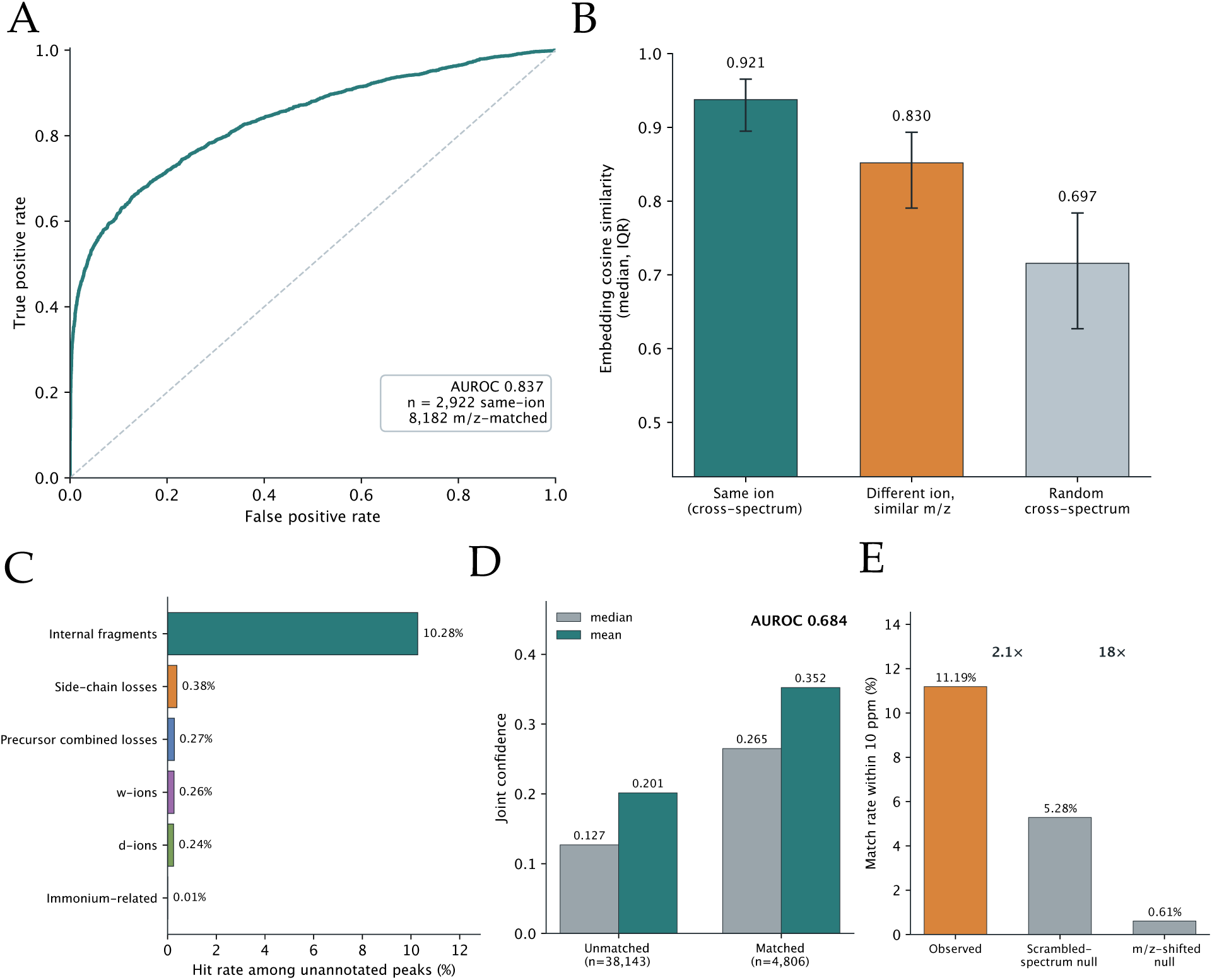
Cross-spectrum ion identity and off-database chemistry of unannotated peaks. **A.** ROC for distinguishing the same fragment ion observed in different spectra of the same peptide from *m/z*-matched cross-spectrum pairs (AUROC 0.837; *n* = 2,922 same-ion, 8,182 *m/z*-matched). **B.** Embedding cosine similarity: same ion cross-spectrum (0.921) vs. a different ion at similar *m/z* (0.830) vs. random cross-spectrum pairs (0.697). **C.** Per-category hit rate among unannotated peaks matching off-database fragment species within 10 ppm (internal fragments dominate at 10.3%, with contributions from side-chain losses, precursor combined losses, and *d*/*w*-ions). **D.** Matched off-database peaks receive higher joint confidence (median 0.265) than unmatched ones (0.127; AUROC 0.684). **E.** Observed match rate (11.2%) versus scrambled-spectrum (2.1-fold) and *m/z*-shifted (18-fold) null models (Supplementary Table S7).

**Figure S15:**
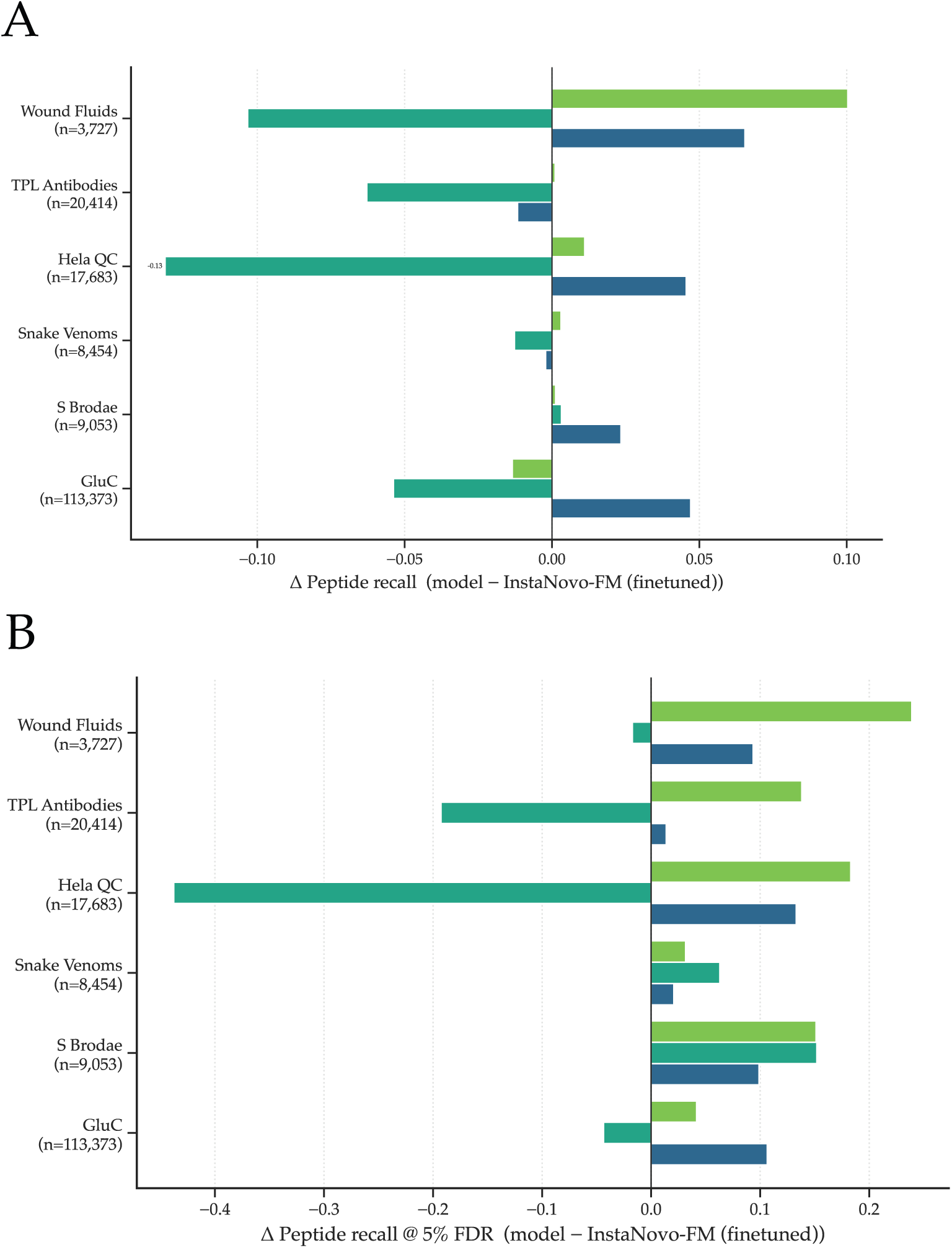
*De novo* peptide recall across biological datasets. **A.** Absolute recall improvement (Δ) over InstaNovo-FM (fine-tuned). **B.** Recall Δ at 5% FDR.

**Figure S16:**
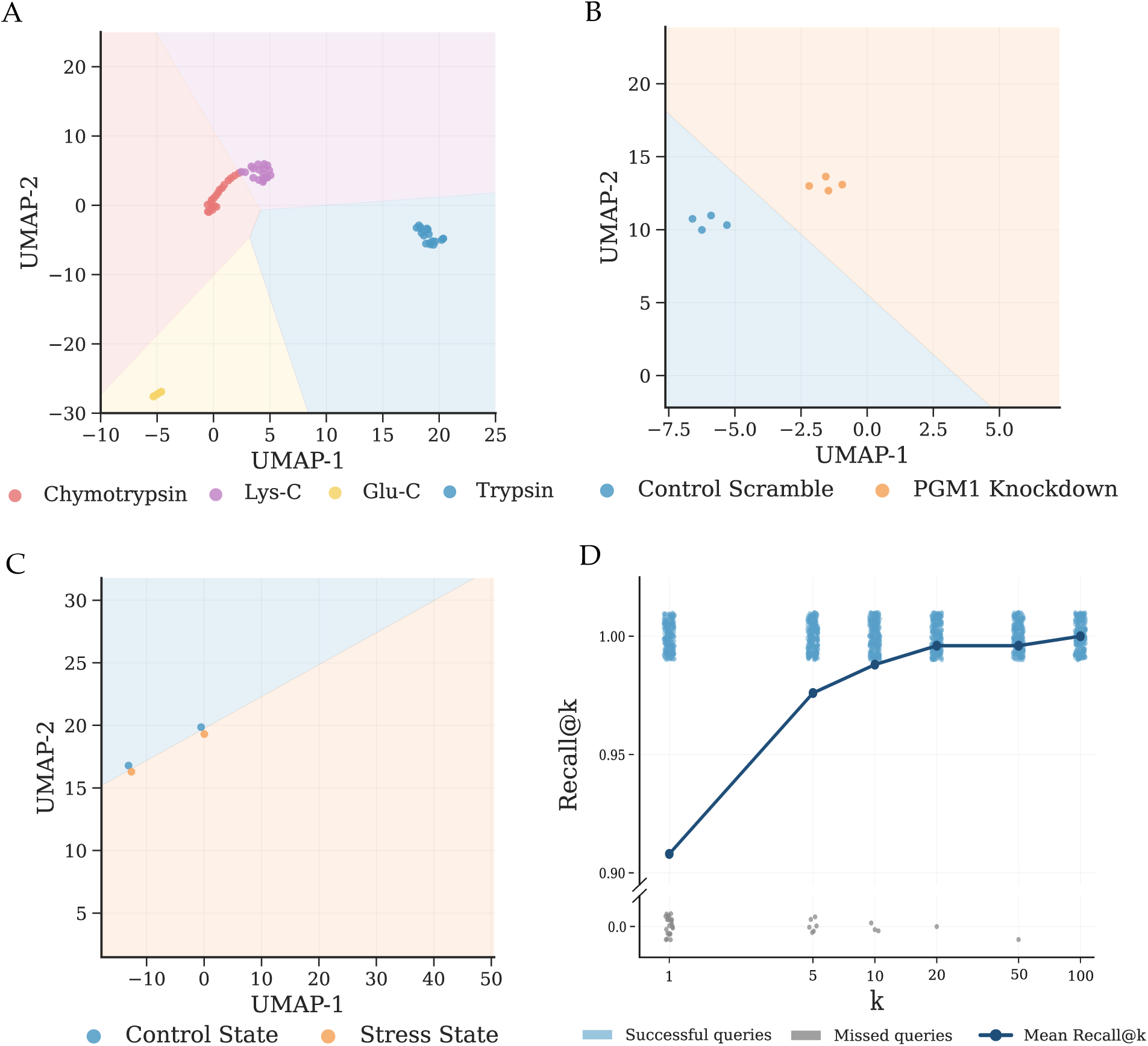
Run-condition classification and duplicate-spectrum retrieval. A–C. UMAP projections of InstaNovo-FM spectrum embeddings, coloured by known experimental condition, with linear-probe decision boundaries fit by logistic regression; in each cohort, conditions occupy distinct, well-separated embedding regions. **A.** Digestion enzyme (chymotrypsin, Lys-C, Glu-C, trypsin). **B.** Genetic perturbation (control-scramble vs. PGM1 knockdown). **C.** Biological stress state (control vs. stress). **D.** Recall@*k* for query–reference retrieval on the spectral rescue task (per-query points and mean), reaching *∼*1.0 by *k* = 20.

**Figure S17:**
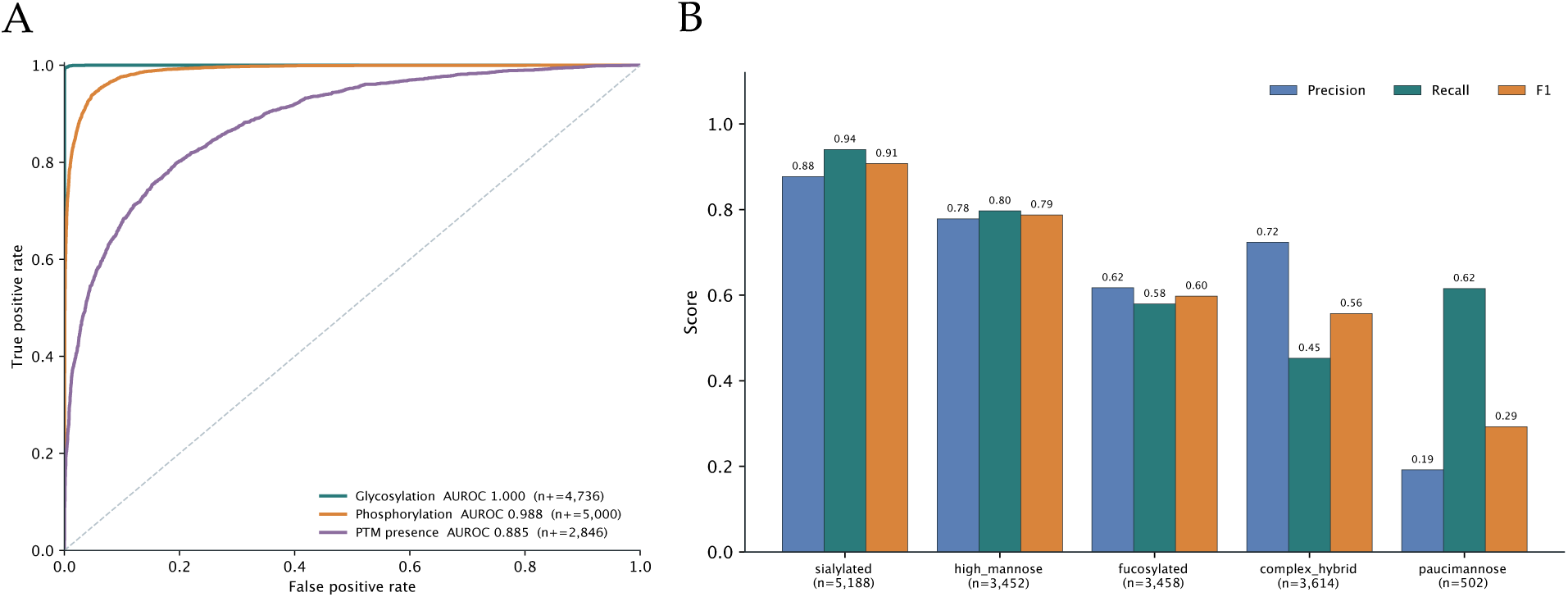
Modification detectability and. *N* **-glycan family identification. A.** Modification-detection ROC from frozen-embedding linear probes: glycosylation (AUROC 1.000, *n*_+_ = 4,736), phosphorylation (0.988, *n*_+_ = 5,000) and overall PTM presence (0.885, *n*_+_ = 2,846), illustrating that detectability scales with mass-shift magnitude. **B.** Five-way *N* -glycan family identification from frozen InstaNovo-FM embeddings: per-class precision, recall and F1 for sialylated, high-mannose, fucosylated, complex/hybrid and paucimannose classes (macro-F1 0.628, balanced accuracy 0.714 vs. 0.320 majority baseline), recovering sialylated and high-mannose classes with high fidelity.

**Figure S18:**
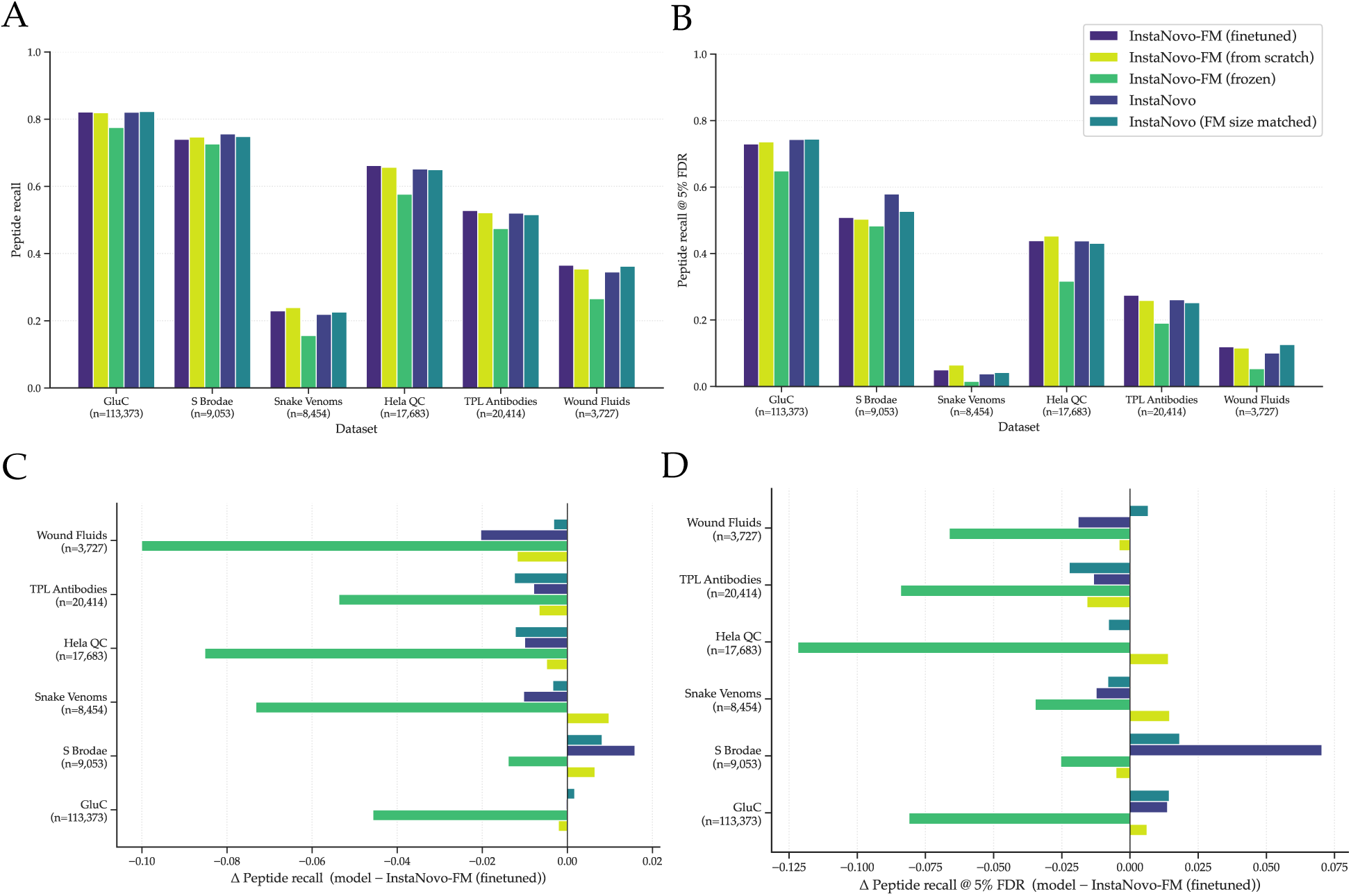
*De novo* sequencing initialisation and model-variant comparison. Peptide recall across the six biological validation datasets for InstaNovo-FM-initialised fine-tuning vs. from-scratch vs. frozen-encoder vs. standard InstaNovo and its FM-size-matched variant. **A.**, Full peptide recall. Numeric values in Supplementary Table S13. **B.** Peptide recall at 5% FDR. **C.** Recall Δ relative to InstaNovo-FM (fine-tuned). **D.** Recall Δ relative to InstaNovo-FM (fine-tuned) at 5% FDR.

## Supplementary Tables

**Table S1:** Dataset accessions used in the pretraining corpus. All 92 accessions comprising the assembled corpus, grouped by the selection category described in the Methods.

| Accession | Dataset title | Reference |
| --- | --- | --- |
| **Large-scale studies selected by automated annotation** |  |  |
| MSV000090792 | Enhancing single-cell proteomics through tailored Data-Independent Acquisition and micropillar array-based chromatography | [65] |
| PXD002052 | Machine Learning Based Classification of Diffuse Large B-cell Lymphoma Patients by their Protein Expression Profiles | [66] |
| PXD003977 | Expanding proteome coverage with CHarge Ordered Parallel Ion aNalysis (CHOPIN) combined with broad specificity proteolysis | [67] |
| PXD005573 | Optimization of Experimental Parameters in Data-Independent Mass Spectrometry Significantly Increases Depth and Reproducibility of Results | [68] |
| PXD006201 | Characterisation of protein ubiquitination using UbiSite technology | [69] |
| PXD014525 | Rapid and site-specific deep phosphoproteome profiling by data-independent acquisition (DIA) without the need for spectral libraries | [10] |
| PXD023064 | Systematic HLA Epitope Ranking Pan Algorithm (SHERPA) | [70] |
| PXD037009 | Global, site-resolved analysis of ubiquitylation occupancy and turnover rate reveals systems properties | [71] |
| PXD043989 | Immunopeptidomics-based identification of naturally presented non-canonical circRNA-derived peptides | [72] |
| PXD044135 | Quantitative differences in rumen epithelium proteins in lambs fed wheat, perennial wheat or perennial wheat plus lucerne. | [73] |
| PXD044169 | Thermal proteome profiling of sponge whole-body movement | [74] |
| PXD044301 | Welfare of Rainbow Trout at Slaughter: Integrating Behavioural, Physiological, Proteomic and Quality Indicators | [75] |
| PXD044303 | Reconstruction of Metabolic Pathways, Protein Expression, and Homeostasis Machineries across Maize Bundle Sheath and Mesophyll Chloroplasts: Large-Scale Quantitative Proteomics Using the First Maize Genome Assembly | [76] |
| PXD044325 | Effect of an immune challenge and feed supplements on broiler chicken individual breast muscle protein synthesis | [77] |
| PXD044360 | AMPK activator 991 specifically activates SnRK1 and thereby affects seed germination in rice | [78] |
| PXD044445 | Remodeling of the human skeletal muscle proteome found after long-term endurance training but not after strength training | [79] |
| PXD044830 | The phosphorylation landscape of infection-related development by the rice blast fungus | [80] |
| PXD045299 | SARS-CoV-2 Envelope TurboID LFQ using internal tagging | [81] |
| PXD045457 | 15 cm Aurora Gen3 column benchmarking on an Exploris 480 | [82] |
| PXD045471 | Human breast adipose tissue extracellular vesicles | [83] |
| PXD045500 | Benchmarking CHIMERYS using Wide Window Acquisition and Classical Narrow Isolation Window Data Dependent Acquisition | [82] |
| PXD045650 | Quantitative profiling of proteome extracted from zebrafish embryos exposed to different concentrations of the fungicide difenoconazole against untreated control groups. | [84] |
| PXD045662 | Ubiquiton – an inducible, linkage-specific polyubiquitylation tool | [85] |
| PXD045779 | Mapping the substrate landscape of protein phosphatase 2A catalytic subunit PPP2CA | [86] |
| PXD046182 | Cervical Cancer Tissue Immunopeptidomics | [87] |
| PXD046211 | Benchmarking of single cell proteomics data analysis workflows | [88] |
| PXD046286 | DCAF1-based PROTACs with activity against clinically validated targets overcoming intrinsic- and acquired-degrader resistance | [89] |
| PXD046451 | iTRAQ-based proteomics reveals potential markers and treatment pathways for acute Achilles tendon rupture | [90] |
| PXD046453 | Ultra-fast label-free quantification and comprehensive proteome coverage with narrow-window data-independent acquisition. DDA vs DIA comparison | [91] |
| PXD046460 | Pure and mixed crude extracts of *Thaurea aromatica* K172 and *Pseudomonas putida* KT2440 for benchmarking GroEL-proteotyping | [92] |
| PXD046802 | DIA-based phosphoproteomics identifies early phosphorylation events in response to EGTA and mannitol in Arabidopsis | [93] |
| PXD046911 | Candida auris proteomic profiling | [94] |

Table S1 continued from previous page
| Accession | Dataset title | Reference |
| --- | --- | --- |
| PXD047134 | Large-scale proteomics analysis of five brain regions from Parkinson's disease patients with a GBA1 mutation | [95] |
| PXD047226 | VAP spatially stabilizes dendritic mitochondria to locally support synaptic plasticity | [96] |
| PXD047641 | NeoDisc: A neoantigen discovery pipeline for the development of personalized cancer immunotherapy | [97] |
| PXD047705 | Proteins of anti-sigma70 and anti-RNAP immunoprecipitates in bacteria | [98] |
| PXD047761 | Proteomic data of the Trypanosoma cruzi RNA-binding protein UBP1 induction and knockdown in insect-dwelling epimastigotes | [148] |
| PXD047762 | An automated and fast sample preparation workflow for laser microdissection guided ultrasensitive proteomics | [99] |
| PXD047782 | Ubiquitinome of Plasmodium yoelii lines NSS and NSR | [100] |
| PXD047839 | timsTOF HT improves protein identification and quantitative reproducibility for deep unbiased plasma protein biomarker discovery (Cancer cohort dataset) | [101] |
| PXD047873 | A CTL-inspired Killing System Reveals Ultralow-dose Chemotherapy is Sufficient for Inducing a Pyroptosis-mediated Antitumour Immune Function | [102] |
| PXD048219 | The natural diversity of the yeast proteome links aneuploidy tolerance to protein turnover | [103] |
| PXD048453 | The Secretome of Macrophages Has a Differential Impact on Spinal Cord Injury Recovery According to the Polarization Protocol | [104] |
| PXD048617 | The post-biotic sodium butyrate synergizes the antiproliferative effects of dexamethasone against the AGS gastric adenocarcinoma cell | [105] |
| PXD049028 | The one hour human proteome: Orbitrap Astral Mass Spectrometer | [106] |
| PXD057992 | Wound healing degradomics | [149] |
| **Landmark studies** |  |  |
| PXD000561 | A draft map of the human proteome | [107] |
| PXD000865 | Mass spectrometry based draft of the human proteome | [108] |
| PXD004452 | HeLa proteome of 12,250 protein-coding genes | [109] |
| PXD013868 | Arabidopsis proteomic tissue atlas | [110] |
| PXD014877 | The Proteome Landscape of the Kingdoms of Life | [12] |
| PXD019483 | The Proteome Landscape of the Kingdoms of Life | [12] |
| PXD024364 | Deep Human Proteome Sequencing Enables Global Detection of Mutations and Alternative Splicing | [150] |
| PXD029360 | Generation of ENSEMBL-based proteogenomics databases boost the identification of novel peptides | [111] |
| **ProteomeTools** |  |  |
| PXD004732 | ProteomeTools - Building ProteomeTools based on a complete synthetic human proteome | [112] |
| PXD009449 | Systematic characterization of 21 post-translational modification using synthetic peptides | [113] |
| PXD010595 | ProteomeTools – Part II - Prosit: proteome-wide prediction of peptide tandem mass spectra by deep learning | [23] |
| PXD021013 | ProteomeTools – Part III - HLA Class I & II & non-tryptic peptides | [112] |
| PXD056559 | ProteomeTools Part VI - Citrullinated peptides | [114] |
| **Nine-species benchmark** |  |  |
| PXD003868 | Quantitative global proteomics of yeast PBP1 deletion mutants and their stress responses reveals glucose metabolism, translation and stress granule changes | [116] |
| PXD004325 | Combination of bottom up 2D-LC-MS and intact protein separation enhances coverage of proteome and low molecular weight short open reading frame encoded peptides of the archeon Methanosarcina mazei | [117] |
| PXD004424 | Proteomic and bioinformatic characterization of extracellular vesicles released from human primary macrophages upon influenza A infection | [118] |
| PXD004467 | Proteome analysis of the hemolymph, mushroom body, and antenna of honeybee resistance against Varroa infestation | [119] |
| PXD004536 | Proteomics of Candidatus Thiodiazotropha endoloripes, the chemosynthetic symbiont of the lucinid clam Loripes lucinalis from the Bay of Fetovaia | [120] |
| PXD004565 | Mini-Bacillus - Large-scale reduction of the Bacillus subtilis genome: Consequences for the transcriptional network, resource allocation, and metabolism | [121] |
| PXD004947 | Characterization of the tomato fruit pericarp proteome (2) | [122] |
| PXD004948 | Cystinosin interactomics - Impact of cystinosin glycosylation on protein stability by differential dynamic SILAC | [123] |

Table S1 continued from previous page
| Accession | Dataset title | Reference |
| --- | --- | --- |
| PXD005025 | Label-free Proteomics Reveals that Cowpea Severe Mosaic Virus Transiently Suppresses the Host Leaf Protein Accumulation During the Compatible Interaction with Cowpea (*Vigna unguiculata* [L.] Walp.) | [124] |
| **Complementary studies targeting metadata gaps)** |  |  |
| PXD000900 | Confetti: A Multi-protease Map of the HeLa Proteome for Comprehensive Proteomics | [125] |
| PXD001258 | Correction To - Confetti: A Multi-protease Map of the HeLa Proteome for Comprehensive Proteomics | [125] |
| PXD006939 | High-throughput MS-based immunopeptidomics | [126] |
| PXD009935 | Determinants of the cellular immunopeptidome -cellular peptidome | [151] |
| PXD010154 | A deep proteome and transcriptome abundance atlas of 29 healthy human tissues | [127] |
| PXD014017 | Immunopeptidomics of colorectal cancer organoids | [128] |
| PXD025859 | Structural Interpretation of Glycosite-specific N-Glycans Using Tandem Mass Spectrometry | [129] |
| PXD026629 | Precision N-glycoproteomics of murine peritoneal macrophage | [130] |
| PXD026649 | Precision mapping of glycosite-specific glycans reveals distinctive N-glycosylation on human spermatozoa | [131] |
| PXD031025 | Human serum glycoproteome - Glyco-Decipher | [132] |
| PXD031032 | Re-analysis of glycoproteomics data with Glyco-Decipher | [132] |
| PXD035158 | Glycoproteomic analysis of Fut8 knock out/wild type mouse brain, Human IgG and Human Serum. | [133] |
| PXD036161 | High-throughput proteomics of the 26 medically most important elapids and vipers from sub-Saharan Africa | [134] |
| PXD044641 | Site- and structure-specific glycosylation signatures of bovine, caprine, porcine and human milk-derived extracellular vesicles | [135] |
| PXD047450 | Structure and mechanism of the Zorya anti-phage defense system | [136] |
| PXD047898 | Maximizing glycoproteomics identification depth in complex mixtures on the timsTOF Pro platform | [137] |
| PXD055983 | Differential protein expression in new1 knock-out yeast | [138] |
| PXD058160 | Expanding the Landscape of Aging via Orbitrap Astral Mass Spectrometry and Tandem Mass Tag (TMT) Integration | [139] |
| PXD058913 | Comprehensive analysis of the proteome of *S. cerevisiae* wild-type and pdr5<sup>-</sup> cells in response to Bisphenol A (BPA) exposure | [140] |
| PXD062877 | Proteomic analysis of innate immune responses in Polg D257A macrophages challenged with *Pseudomonas aeruginosa* | [141] |
| PXD063058 | Proteome profiling of cerebrospinal fluid and machine learning reveal biomarkers of two forms of Alzheimer's disease characterized by increased or not altered levels of tau | [142] |
| PXD065790 | AuxB interacts directly with GpsB and PknB to coordinate cell envelope processes that contribute to intrinsic antibiotic resistance in *Staphylococcus aureus* | [143] |
| PXD068040 | Differential interaction proteomics and phosphoproteomics of STK17B on an Orbitrap Astral, Nikiforov | [152] |
| PXD068926 | Mycobacterium tuberculosis Quantitative proteomics | [144] |

**Table S2:** Binning strategy comparison. Jump rate, collision rate, combined score, vocabulary size, and effective information (bits) for all 12 evaluated strategies, with the deployed fixed 0.2 Da grid shown beneath for reference. Note that adaptive ultra coarse is numerically identical to a fixed 0.5 Da grid, and adaptive baseline is all but indistinguishable from the deployed 0.2 Da grid (12,124 versus 12,250 bins; combined score 4.72% versus 4.70%), which is why the adaptive family was not carried forward.

| Strategy | $N_{\text{bins}}$ | Jump rate | Collision rate | Combined | Eff. info (bits) |
| --- | --- | --- | --- | --- | --- |
| fixed_da_0.1 | 24,500 | 5.4% | 3.71% | 4.54% | 11.62 |
| fixed_da_0.05 | 49,000 | 8.5% | 2.19% | 5.34% | 11.98 |
| fixed_da_0.02 | 122,500 | 14.3% | 0.57% | 7.46% | 12.21 |
| fixed_da_0.01 | 245,000 | 21.4% | 0.16% | 10.76% | 11.88 |
| fixed_ppm_50 | 78,243 | 10.6% | 0.08% | 5.35% | 12.42 |
| fixed_ppm_20 | 195,604 | 17.4% | 0.01% | 8.70% | 12.41 |
| fixed_ppm_10 | 391,205 | 27.7% | 0.00% | 13.87% | 11.49 |
| adaptive_high_res | 79,863 | 12.2% | 0.58% | 6.37% | 12.33 |
| adaptive_med_res | 50,371 | 9.6% | 0.90% | 5.23% | 12.26 |
| adaptive_low_res | 32,623 | 7.5% | 1.88% | 4.69% | 12.03 |
| adaptive_baseline | 12,124 | 2.8% | 6.63% | 4.72% | 11.15 |
| adaptive_ultra_coarse | 4,900 | 1.1% | 8.84% | 4.99% | 10.58 |
| **fixed_da_0.2 (deployed)** | **12,250** | **3.0%** | **6.40%** | **4.70%** | **11.18** |

**Table S3:** Masking strategy comparison. Five of the six evaluated masking configurations, measured on 200,000 LCFM spectra; the sixth is a second Thompson-span-with-isotope run at a 25% fragment-group budget, omitted here because the deployed model uses the 30% budget shown. Mask ratio is the achieved fraction of peaks masked, which exceeds the configured budget for the uniform and Thompson strategies. Signal efficiency is the fraction of masked peaks that are annotated fragment ions. Leakage is the pooled information leakage, the fraction of masked fragments whose complementary or neutral-loss partner survives elsewhere in the spectrum. Partial groups is the fraction of fragment groups left partly masked, expressed over the mean number of groups per spectrum. Mean gap is the mean *m/z* distance from a masked peak to its nearest surviving neighbour.

| Strategy | Mask ratio | Signal eff. | Leakage | Partial groups | Mean gap (Da) |
| --- | --- | --- | --- | --- | --- |
| uniform | 45.5% | 31.5% | 46.6% | 30.0% | 6.8 |
| thompson | 44.0% | 35.0% | 52.6% | 29.9% | 6.4 |
| thompson_span | 30.2% | 35.2% | 34.8% | 7.1% | 14.1 |
| thompson_span_isotope | 26.6% | 35.9% | 29.1% | 1.6% | 16.0 |
| signal_aware_fragment | 15.1% | 75.7% | 8.5% | 1.6% | 17.4 |

**Table S4:** Masking-strategy and PA-bias factorial evaluation. 2*×*2 factorial across masking strategy (signal-aware fragment masking, SA, vs. Thompson-span masking with isotope co-masking, TS) and pairwise- attention bias (PA vs. noPA) on LCFM-trained checkpoints (*∼*230,000 steps), evaluated on the LCFM validation split (the ablation split; deployed-model results reported elsewhere are on the held-out test split). Fragment-type F1 is averaged over the four classes with support in the test split; the deployed-model tables report 0.855 for this checkpoint rather than 0.865 because spectra with unrecorded fragmentation are additionally excluded from probe training there. Bold indicates the best value per metric row.

| Metric | SA·noPA | SA·PA | TS·noPA | TS·PA |
| --- | --- | --- | --- | --- |
| Fragment type F1 (4-class) | 0.793 | 0.804 | **0.865** | 0.757 |
| Instrument macro F1 | 0.777 | 0.772 | **0.807** | 0.675 |
| Spectrum conf. $R^2$ | 0.927 | 0.937 | **0.973** | 0.971 |
| PTM bal. accuracy | 0.777 | 0.795 | **0.802** | 0.708 |
| Mod. class macro F1 | 0.582 | **0.631** | 0.620 | 0.505 |
| Precursor $m/z$ $R^2$ | 0.937 | **0.940** | 0.929 | 0.884 |
| Cross-spectrum AUROC | **0.874** | 0.870 | 0.857 | 0.759 |
| Recall@1 overall | **0.406** | 0.375 | 0.400 | 0.324 |
| Recall@1 CID | 0.455 | **0.473** | 0.426 | 0.331 |
| Structural heads / 12 | 1 | 1 | **8** | 4 |
| Mean isotope enrichment | 1.39 | 0.54 | **1.99** | 0.66 |
| Peak type Macro F1 | 0.559 | 0.562 | **0.575** | 0.468 |

**Table S5:** Representation quality of the curated-subset MCFM baseline (9-layer, *∼*90,000 steps) versus the full-corpus deployed model (12-layer, *∼*230,000 steps); both use identical Thompson-span (TS·noPA) masking and are evaluated on the *same* held-out LCFM spectra with identical frozen-probe protocols. Bold marks the stronger value where the gap exceeds probe-level noise; confidence is effectively tied. Macro-F1 is averaged over the classes present in the test split.

| Frozen-probe metric | MCFM baseline (9L, 90K) | Deployed LCFM (12L, 230K) |
| --- | --- | --- |
| Fragment type macro F1 (4-class) | 0.781 | **0.855** |
| Instrument macro F1 (13-class) | 0.697 | **0.804** |
| PTM presence bal. accuracy | 0.751 | **0.802** |
| Hydrophobicity $R^2$ | 0.518 | **0.605** |
| Precursor mass $R^2$ | 0.698 | **0.732** |
| Precursor $m/z$ $R^2$ | 0.896 | **0.929** |
| Precursor charge macro F1 (7-class) | 0.602 | **0.650** |
| Spectrum confidence $R^2$ | **0.978** | 0.973 |
| Duplicate retrieval Recall@1 | **0.307** | 0.215 |
| Duplicate retrieval mAP@20 | **0.126** | 0.076 |

**Table S6:** Dataset tier quality metrics. Spectral annotation quality, charge distribution, and masking performance across HCFM, MCFM, and LCFM tiers (200,000 spectra each).

| Metric | HCFM | MCFM | LCFM |
| --- | --- | --- | --- |
| Mean peaks/spectrum | 134.0 | 135.3 | 135.0 |
| Mean fragment groups/spectrum | 13.6 | 15.9 | 12.3 |
| Backbone coverage | 63.5% | 67.3% | 58.3% |
| Mean matched peaks | 32.4 | 38.7 | 28.8 |
| Quality gate rejection | 19.4% | 14.8% | 15.6% |
| SAF training signal efficiency | 60.4% | 64.1% | 62.6% |
| Mean precursor charge | 2.16 | 2.23 | 2.43 |
| b-ion median abs. PPM | 5.1 | 3.9 | 4.1 |
| y-ion median abs. PPM | 3.5 | 3.0 | 3.2 |

**Table S7:** Off-database chemistry of high-attention unannotated peaks (TS·noPA). Of *∼*42,900 unannotated peaks examined by integrated-gradients attribution, 11.2% match a defined off-database fragment species within 10 ppm — a 2.1-fold enrichment over a sequence-scrambled decoy library and 18-fold over *m/z*-shifted nulls. The hit-rate column is the overall per-category match rate; remaining columns give matched-peak counts stratified by fragmentation method. No immonium-ion enrichment was detected (0.39-fold, i.e. depleted relative to the null). Matched peaks receive higher model confidence than unmatched unannotated peaks (median joint confidence 0.265 vs. 0.127; AUROC 0.684). Source: deployed InstaNovo-FM TS·noPA checkp<u>oint, LCFM test split.</u>

| Off-database species | Hit rate | HCD | CID | HCD/CID | ETD |
| --- | --- | --- | --- | --- | --- |
| Internal fragment | 10.3% | 4198 | 182 | 34 | — |
| Side-chain loss | 0.38% | 95 | 68 | — | — |
| Precursor combined loss | 0.27% | 71 | 36 | 11 | — |
| w-ion | 0.26% | 93 | 17 | 1 | 1 |
| d-ion | 0.24% | 74 | 27 | 3 | — |
| Immonium-related | 0.01% | — | 5 | — | — |

**Table S8:** InstaNovo-FM training hyperparameters.

| Hyperparameter | Value |
| --- | --- |
| Model dimension ($d$) | 768 |
| Attention heads | 12 (head dim 64) |
| Encoder layers | 12 |
| Feedforward dimension | 3072 |
| Dropout | 0.1 |
| Peak encoder | Multi-scale sinusoidal (learned frequencies) |
| Positional encoding | None (peaks processed as unordered set) |
| Masked-peak encoding | Gaussian-blurred $m/z$ ($\sigma = 10$ Da) + visible intensity |
| Pairwise attention bias | None |
| Attention backend | Flash (PyTorch SDPA) |
| Total parameters | $\approx 89.5$M |
| Masking strategy | Thompson span + isotope co-masking |
| Fragment-group mask budget | 30% ($\approx 26.6\%$ of peaks on LCFM) |
| Span size | 3–4 peaks |
| Isotope co-masking spacing | $1.003355/z$ Da |
| Binning | Fixed resolution, 0.2 Da |
| $m/z$ range | [50, 2500] Da |
| Number of bins | 12,250 (245 groups × 50 offsets) |
| Loss weights ($w_{\text{group}}, w_{\text{offset}}$) | 0.5, 0.5 |
| Intensity head | 2-layer MLP, Huber ($\delta = 0.05$), $\lambda_{\text{intensity}} = 0.2$ |
| Peak learning rate | $10^{-4}$ |
| LR schedule | Cosine warmup-hold-decay |
| Warmup fraction | 5% |
| Hold fraction | 30% |
| Final learning rate | $10^{-5}$ |
| Batch size | 1,024 |
| Training steps | $\sim 230,000$ |
| Precision | fp16 |
| Gradient clip | 1.0 |
| Gradient checkpointing | Enabled |
| Checkpoint metric | median_ae_ppm (min) |
| Checkpoint interval | 10,000 steps (top-3 retained) |

**Table S9:** Data subsets used for each evaluation. Summary of corpus, split, and sample size across all evaluation experiments. LCFM splits are peptide-disjoint 80/10/10.

| Evaluation | Corpus / split | Sample size |
| --- | --- | --- |
| Factorial ablation (frozen-embedding probes) | LCFM / validation | same probe protocol (below) |
| Linear-probe battery (incl. external-benchmark comparison) | LCFM / train-val-test | 100,000 / 10,000 / 10,000 |
| Duplicate retrieval | LCFM / test | 20,000 groups (200,000-spectrum pool) |
| Duplicate rescue | PXD074343 / run 477-1 | 30,404 spectra |
| Peak-type, cross-spectrum ion identity, IG attribution | LCFM / test | 10,000 spectra (∼411,000 peaks) |
| UMAP visualisation | LCFM / test | 100,000 (140,000-spectrum pool) |
| Signal composition / theoretical fragment annotation | HCFM / validation | 200 spectra |
| *Hela qc* probe comparison | *Hela qc* / 80-10-10 | $n = 17,683$ |

**Table S10:** Linear-probe and duplicate-retrieval performance across models on the held-out LCFM dataset. We exclude precursor metrics for XuanjiNovo since precursor information is fed to their encoder.

| Metric | InstaNovo-FM | IN v1.2 | Casanovo | XuanjiNovo |
| --- | --- | --- | --- | --- |
| Precursor charge macro F1 | **0.650** | 0.551 | 0.461 | – |
| Precursor $m/z$ $R^2$ | **0.929** | 0.915 | 0.837 | – |
| Precursor mass $R^2$ | **0.732** | 0.672 | 0.573 | – |
| Hydrophobicity $R^2$ | 0.605 | 0.614 | 0.525 | **0.704** |
| PTM presence bal. accuracy | 0.802 | **0.810** | 0.737 | 0.793 |
| Modification class macro F1 | 0.622 | **0.688** | 0.532 | 0.619 |
| Search instrument macro F1 | **0.804** | 0.690 | 0.538 | 0.516 |
| Fragmentation type macro F1 | **0.855** | 0.809 | 0.725 | 0.736 |
| Duplicate retrieval Recall@1 | 0.215 | 0.337 | 0.238 | **0.378** |
| Duplicate retrieval mAP@20 | 0.076 | 0.162 | 0.096 | **0.171** |

**Table S11:** Linear-probe performance across models on *Hela qc* dataset. We exclude precursor metrics for XuanjiNovo since precursor information is fed to their encoder.

| Metric | InstaNovo-FM | IN v1.2 | Casanovo | XuanjiNovo |
| --- | --- | --- | --- | --- |
| Precursor charge macro F1 | 0.577 | **0.610** | 0.560 | – |
| Precursor $m/z$ $R^2$ | 0.951 | **0.956** | 0.920 | – |
| Precursor mass $R^2$ | 0.887 | **0.889** | 0.837 | – |
| Hydrophobicity $R^2$ | 0.568 | 0.670 | 0.601 | **0.727** |
| PTM presence bal. accuracy | 0.761 | 0.811 | 0.782 | **0.816** |
| Modification class macro F1 | 0.545 | 0.603 | 0.641 | **0.669** |

**Table S12:** Peptide recall across the biological validation datasets for the de novo sequencing benchmark. The best model per dataset is in bold.

| Dataset | IN-FM (fine-tuned) | IN v1.2 | Casanovo | XuanjiNovo |
| --- | --- | --- | --- | --- |
| GluC | 0.821 | **0.868** | 0.767 | 0.808 |
| S Brodae | 0.740 | **0.764** | 0.743 | 0.741 |
| Snake Venoms | 0.230 | 0.227 | 0.217 | **0.233** |
| Hela QC | 0.662 | **0.708** | 0.531 | 0.673 |
| TPL Antibodies | 0.528 | 0.517 | 0.465 | **0.529** |
| Wound Fluids | 0.366 | 0.431 | 0.262 | **0.466** |

**Table S13:** Peptide recall across the biological validation datasets for the de novo sequencing InstaNovo-FM and InstaNovo model variant benchmarks. The best model per dataset is in bold. We abbreviate “InstaNovo (FM size matched)” to “IN (FM-SM)” to conserve page space.

| Dataset | IN-FM (fine-tuned) | IN-FM (from scratch) | IN-FM (frozen) | IN | IN (FM-SM) |
| --- | --- | --- | --- | --- | --- |
| GluC | 0.821 | 0.819 | 0.775 | 0.821 | **0.823** |
| S Brodae | 0.740 | 0.747 | 0.726 | **0.756** | 0.748 |
| Snake Venoms | 0.230 | **0.239** | 0.156 | 0.219 | 0.226 |
| Hela QC | **0.662** | 0.657 | 0.577 | 0.652 | 0.650 |
| TPL Antibodies | **0.528** | 0.522 | 0.474 | 0.520 | 0.516 |
| Wound Fluids | **0.366** | 0.354 | 0.266 | 0.345 | 0.362 |

**Table S14:** Quality-filter attrition per confidence tier. PSMs removed from each unsplit tier by each retention criterion. Each PSM is counted once for every criterion it fails, so a PSM failing two criteria contributes to both counts. Two criteria remove nothing at all: no PSM in the corpus carries a precursor charge above 7 or a precursor *m/z* above 2,000 Da, either before or after filtering. The unresolved-modification criterion likewise removes nothing from MCFM or HCFM, because those PSMs are already excluded during tier construction. The per-criterion counts therefore sum exactly to the number removed for those two tiers, while LCFM contains 101,009 PSMs that fail more than one criterion.

| Criterion | LCFM | MCFM | HCFM |
| --- | --- | --- | --- |
| Unsplit tier PSMs | 184,607,213 | 18,422,236 | 3,684,448 |
| Retention time > 10,800 s | 2,537,627 | 166,138 | 14,052 |
| Lower isolation offset > 300 Da | 8,147 | 833 | 283 |
| Unresolved modification annotation | 384,857 | 0 | 0 |
| Precursor charge > 7 | 0 | 0 | 0 |
| Precursor $m/z$ > 2,000 Da | 0 | 0 | 0 |
| Retained (training partition) | 181,777,591 | 18,255,265 | 3,670,113 |
| Retained fraction | 98.47% | 99.09% | 99.61% |

**Table S15:** Precursor charge by acquisition mode in the LCFM training partition. A precursor charge of 0 is recorded only for DIA spectra, where the precursor is an isolation window rather than a selected ion and no single charge state is assigned. Every spectrum is labelled DDA or DIA; the acquisition field is never absent.

| Precursor charge | DDA | DIA | Total |
| --- | --- | --- | --- |
| 0 | 0 | 6,521,868 | 6,521,868 |
| 1 | 1,897,086 | 0 | 1,897,086 |
| 2 | 102,537,678 | 0 | 102,537,678 |
| 3 | 56,236,913 | 0 | 56,236,913 |
| 4 | 12,173,162 | 0 | 12,173,162 |
| 5 | 2,070,002 | 0 | 2,070,002 |
| 6 | 310,127 | 0 | 310,127 |
| 7 | 30,755 | 0 | 30,755 |
| Total | 175,255,723 | 6,521,868 | 181,777,591 |
| Share of partition | 96.4% | 3.6% | 100% |

**Table S16:**
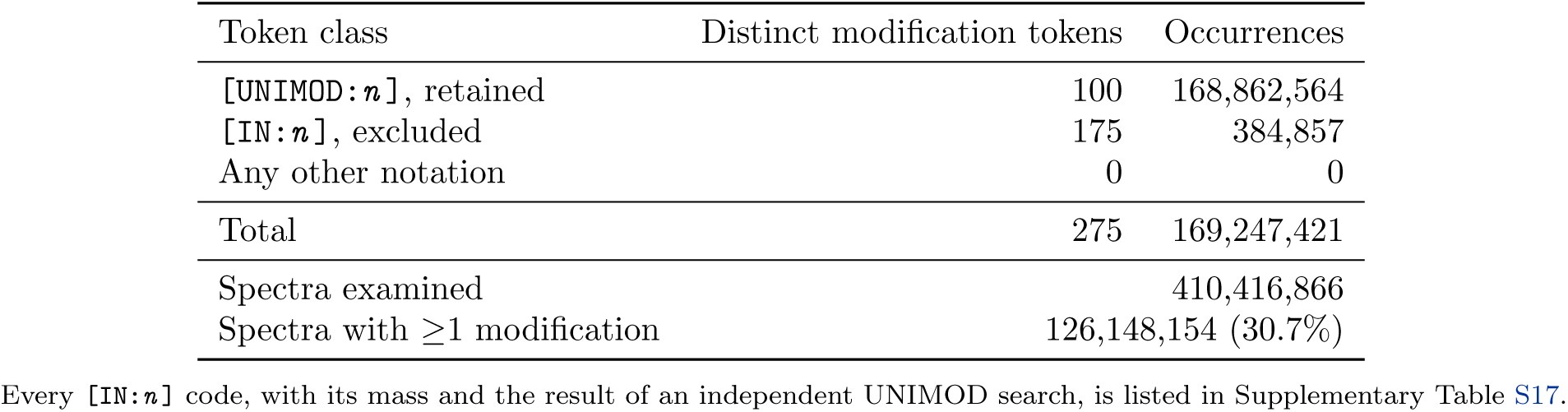
Modification vocabulary of the released corpus. Every bracketed modification token in the sequence column, across all six released datasets. Standard amino acid residues are not counted; a token is counted per residue it occurs on, so one modification seen on two residues counts twice. [UNIMOD:*n*] denotes a modification resolved against UNIMOD and retained; [IN:*n*] is an internal namespace for compositions absent from UNIMOD, which the retention filter excludes. No token in any other notation occurs anywhere, so every modification in the corpus is either resolved against UNIMOD or explicitly marked as unresolved. Glycopeptides are not excluded as a class: those whose glycan carries a UNIMOD identifier are retained.

**Table S17:** Unresolved modification codes in full. All 175 [IN:*n*] codes observed in the corpus, the notation used for a modification that does not resolve to a UNIMOD identifier. Spectra carrying one are excluded by the retention filter, so this is the complete vocabulary of what the quality filters drop on modification grounds. Δ mass is the monoisotopic mass delta in Da. *Nearest* is the distance in Da to the closest UNIMOD record at the same residue, obtained by searching the UNIMOD database distributed with pyOpenMS; a large distance is evidence that no UNIMOD identifier exists for that composition. 165 of the 175 codes lie further than 5 Da from any record, and the median distance is 123 Da. UNIMOD records are identified by elemental composition rather than by mass, and isobaric compositions exist, so a close match is a candidate for re-annotation rather than proof of identity.

| Code | $\Delta$ mass | Spectra | Nearest | Code | $\Delta$ mass | Spectra | Nearest |
| --- | --- | --- | --- | --- | --- | --- | --- |
| N[IN:3000] | 2864.0361 | 1,272 | 3.0360 | N[IN:3088] | 2571.9455 | 2,931 | 219.0995 |
| N[IN:3001] | 2042.7197 | 11,018 | 17.0152 | N[IN:3089] | 2425.8876 | 3,097 | 73.0416 |
| N[IN:3002] | 2334.7977 | 1,205 | 16.0327 | N[IN:3090] | 3210.1374 | 1,382 | 349.1373 |
| N[IN:3003] | 3138.1163 | 939 | 277.1162 | N[IN:3091] | 2174.7983 | 1,715 | 29.9741 |
| N[IN:3004] | 3300.1691 | 778 | 439.1690 | N[IN:3092] | 2570.9251 | 1,078 | 218.0791 |
| N[IN:3005] | 2026.6870 | 9,494 | 28.9888 | N[IN:3093] | 2215.8248 | 582 | 11.0524 |
| N[IN:3006] | 1694.6028 | 7,679 | 0.0001 | N[IN:3094] | 2717.9782 | 2,040 | 143.0219 |
| N[IN:3007] | 2221.7878 | 11,113 | 17.0154 | N[IN:3095] | 2786.9633 | 1,328 | 74.0368 |
| N[IN:3008] | 2075.7299 | 14,628 | 15.9950 | N[IN:3096] | 2569.9047 | 10,974 | 217.0587 |
| N[IN:3009] | 2237.7827 | 9,206 | 33.0103 | N[IN:3097] | 2539.9305 | 511 | 187.0845 |
| N[IN:3010] | 2674.9361 | 1,071 | 186.0640 | N[IN:3098] | 2553.9098 | 2,304 | 201.0638 |
| N[IN:3011] | 2278.8093 | 7,971 | 72.0211 | N[IN:3099] | 3007.0580 | 9,574 | 146.0579 |
| N[IN:3012] | 2643.9415 | 2,359 | 217.0586 | N[IN:3100] | 4176.4914 | 165 | 1315.4913 |
| N[IN:3013] | 2457.8774 | 1,263 | 105.0314 | N[IN:3101] | 4307.5497 | 167 | 1446.5496 |
| N[IN:3014] | 2366.8253 | 11,374 | 13.9793 | N[IN:3102] | 2621.9836 | 375 | 239.0165 |
| N[IN:3015] | 1873.6709 | 1,995 | 1.0204 | N[IN:3103] | 3374.2059 | 1,458 | 513.2058 |
| N[IN:3016] | 2141.7881 | 657 | 49.0429 | N[IN:3104] | 2053.7720 | 391 | 5.9629 |
| N[IN:3017] | 2190.7932 | 1,536 | 13.9792 | N[IN:3105] | 2270.7929 | 807 | 66.0205 |
| N[IN:3018] | 2172.7449 | 826 | 32.0275 | N[IN:3106] | 3147.1152 | 484 | 286.1151 |
| N[IN:3019] | 2076.7503 | 4,320 | 15.9949 | N[IN:3107] | 2528.8781 | 1,018 | 176.0321 |
| N[IN:3020] | 3048.0846 | 603 | 187.0845 | N[IN:3108] | 4103.4750 | 126 | 1242.4749 |
| N[IN:3021] | 2966.0315 | 306 | 105.0314 | N[IN:3109] | 2814.0106 | 889 | 46.9895 |
| N[IN:3022] | 2313.8729 | 399 | 36.9575 | N[IN:3110] | 3284.1742 | 868 | 423.1741 |
| N[IN:3023] | 2586.9200 | 3,024 | 234.0740 | N[IN:3111] | 2919.0420 | 3,090 | 58.0419 |
| N[IN:3024] | 2498.9039 | 1,925 | 146.0579 | N[IN:3112] | 2393.8726 | 895 | 41.0266 |
| N[IN:3025] | 3405.2005 | 1,068 | 544.2004 | N[IN:3113] | 2772.9841 | 2,988 | 88.0160 |
| N[IN:3026] | 2254.7980 | 2,440 | 50.0256 | N[IN:3114] | 3083.1104 | 1,713 | 222.1103 |
| N[IN:3027] | 2133.7717 | 5,606 | 41.0265 | N[IN:3115] | 2012.7455 | 1,706 | 15.0473 |
| N[IN:3028] | 2108.7401 | 1,452 | 15.9949 | N[IN:3116] | 2871.0572 | 282 | 10.0571 |
| N[IN:3029] | 2789.9994 | 1,632 | 71.0007 | N[IN:3117] | 2010.6921 | 618 | 12.9939 |
| N[IN:3030] | 2424.8672 | 7,054 | 72.0212 | N[IN:3118] | 3720.3071 | 256 | 859.3070 |
| N[IN:3031] | 1711.6181 | 3,104 | 1.0204 | N[IN:3119] | 2448.8784 | 2,184 | 96.0324 |
| N[IN:3032] | 3081.0948 | 3,877 | 220.0947 | N[IN:3120] | 3866.3650 | 140 | 1005.3649 |
| N[IN:3033] | 2823.0096 | 774 | 37.9905 | N[IN:3121] | 2263.8347 | 5,438 | 59.0623 |
| N[IN:3034] | 2775.0248 | 1,053 | 85.9753 | N[IN:3122] | 3738.3428 | 794 | 877.3427 |
| N[IN:3035] | 2555.9254 | 842 | 203.0794 | N[IN:3123] | 3663.2857 | 2,440 | 802.2856 |
| N[IN:3036] | 2069.7669 | 834 | 10.0320 | N[IN:3124] | 4101.4594 | 301 | 1240.4593 |
| N[IN:3037] | 3372.1902 | 3,699 | 511.1901 | N[IN:3125] | 2287.8460 | 693 | 62.9844 |
| N[IN:3038] | 2110.7935 | 564 | 18.0483 | N[IN:3126] | 2545.8934 | 428 | 193.0474 |
| N[IN:3039] | 2407.8519 | 3,307 | 55.0059 | N[IN:3127] | 3591.2645 | 798 | 730.2644 |
| N[IN:3040] | 3704.3122 | 337 | 843.3121 | N[IN:3128] | 2158.8034 | 1,685 | 45.9690 |
| N[IN:3041] | 3503.2485 | 289 | 642.2484 | N[IN:3129] | 2279.8297 | 3,172 | 71.0007 |
| N[IN:3042] | 2935.0369 | 5,224 | 74.0368 | N[IN:3130] | 3228.1731 | 329 | 367.1730 |
| N[IN:3043] | 2222.8082 | 1,340 | 18.0358 | N[IN:3131] | 3886.4164 | 256 | 1025.4163 |
| N[IN:3044] | 2060.7554 | 15,933 | 1.0205 | N[IN:3132] | 3809.3436 | 1,234 | 948.3435 |
| N[IN:3045] | 2463.8403 | 836 | 110.9943 | N[IN:3133] | 4272.5238 | 67 | 1411.5237 |
| N[IN:3046] | 2336.8511 | 1,948 | 13.9793 | N[IN:3134] | 4118.4496 | 221 | 1257.4495 |
| N[IN:3047] | 2968.0471 | 521 | 107.0470 | N[IN:3135] | 3594.3006 | 292 | 733.3005 |
| N[IN:3048] | 2205.7929 | 3,916 | 1.0205 | N[IN:3136] | 2465.8937 | 2,596 | 113.0477 |
| N[IN:3049] | 2117.7768 | 3,941 | 25.0316 | N[IN:3137] | 3665.3013 | 963 | 804.3012 |
| N[IN:3050] | 3448.2426 | 762 | 587.2425 | N[IN:3138] | 4483.5818 | 71 | 1622.5817 |
| N[IN:3051] | 2434.8991 | 1,143 | 82.0531 | N[IN:3139] | 4468.6072 | 246 | 1607.6071 |
| N[IN:3052] | 2231.8198 | 1,017 | 27.0474 | N[IN:3140] | 4409.5450 | 233 | 1548.5449 |
| N[IN:3053] | 2256.8514 | 408 | 52.0790 | N[IN:3141] | 2798.0157 | 408 | 62.9844 |
| N[IN:3054] | 2475.9257 | 273 | 123.0797 | N[IN:3142] | 4247.5173 | 118 | 1386.5172 |
| N[IN:3055] | 2245.7990 | 3,521 | 41.0266 | N[IN:3143] | 2701.9833 | 659 | 159.0168 |
| N[IN:3056] | 2116.7564 | 4,672 | 24.0112 | N[IN:3144] | 6441.3465 | 20 | 3580.3464 |
| N[IN:3057] | 3009.0737 | 1,955 | 148.0736 | N[IN:3145] | 2246.8194 | 188 | 42.0470 |
| N[IN:3058] | 2880.0311 | 1,128 | 19.0310 | N[IN:3146] | 3882.3600 | 307 | 1021.3599 |
| N[IN:3059] | 2715.9626 | 8,713 | 145.0375 | N[IN:3147] | 4475.6032 | 99 | 1614.6031 |
| N[IN:3060] | 2188.7398 | 1,537 | 16.0326 | N[IN:3148] | 3827.3541 | 452 | 966.3540 |
| N[IN:3061] | 2391.8569 | 1,405 | 39.0109 | N[IN:3149] | 3520.2638 | 469 | 659.2637 |
| N[IN:3062] | 3234.1487 | 278 | 373.1486 | N[IN:3150] | 3519.2685 | 86 | 658.2684 |
| N[IN:3063] | 4028.4179 | 253 | 1167.4178 | N[IN:3151] | 2142.8085 | 555 | 50.0633 |
| N[IN:3064] | 2018.7084 | 2,732 | 21.0102 | N[IN:3152] | 3302.2099 | 217 | 441.2098 |
| N[IN:3065] | 3517.2278 | 3,526 | 656.2277 | N[IN:3153] | 4412.6187 | 99 | 1551.6186 |
| N[IN:3066] | 2878.0154 | 3,100 | 17.0153 | N[IN:3154] | 1857.6760 | 418 | 1.0204 |
| N[IN:3067] | 2157.7830 | 1,663 | 46.9894 | N[IN:3155] | 4833.7394 | 74 | 1972.7393 |
| N[IN:3068] | 2320.8562 | 1,964 | 29.9742 | N[IN:3156] | 3521.2842 | 85 | 660.2841 |
| N[IN:3069] | 3155.1316 | 523 | 294.1315 | N[IN:3157] | 3140.1570 | 265 | 279.1569 |
| N[IN:3070] | 3153.1159 | 1,738 | 292.1158 | N[IN:3158] | 3959.4328 | 178 | 1098.4327 |
| N[IN:3071] | 2481.8886 | 1,756 | 129.0426 | N[IN:3159] | 5563.0086 | 38 | 2702.0085 |
| N[IN:3072] | 2262.8143 | 9,751 | 58.0419 | N[IN:3160] | 3589.2489 | 127 | 728.2488 |
| N[IN:3073] | 2028.7404 | 1,246 | 30.9945 | N[IN:3161] | 3812.3796 | 137 | 951.3795 |
| N[IN:3074] | 2100.7615 | 4,061 | 8.0163 | N[IN:3162] | 2716.9830 | 486 | 144.0171 |
| N[IN:3075] | 2627.9465 | 3,130 | 233.0536 | N[IN:3163] | 2791.0198 | 729 | 69.9803 |
| N[IN:3076] | 2432.8835 | 4,357 | 80.0375 | N[IN:3164] | 4393.5501 | 132 | 1532.5500 |
| N[IN:3077] | 3226.1323 | 4,801 | 365.1322 | N[IN:3165] | 3156.1520 | 76 | 295.1519 |
| N[IN:3078] | 3229.1683 | 495 | 368.1682 | N[IN:3166] | 3010.1192 | 268 | 149.1191 |
| N[IN:3079] | 3407.2412 | 139 | 546.2411 | N[IN:3167] | 2644.9619 | 449 | 216.0382 |
| N[IN:3080] | 2896.0511 | 684 | 35.0510 | N[IN:3168] | 3447.2474 | 127 | 586.2473 |
| N[IN:3081] | 1695.6232 | 828 | 1.0205 | N[IN:3169] | 2043.7401 | 58 | 15.9948 |
| N[IN:3082] | 2101.7819 | 947 | 9.0367 | N[IN:3170] | 6586.3840 | 18 | 3725.3839 |
| N[IN:3083] | 3064.0795 | 1,498 | 203.0794 | N[IN:3171] | 2408.8723 | 741 | 56.0263 |
| N[IN:3084] | 2580.9570 | 346 | 228.1110 | N[IN:3172] | 6732.4420 | 26 | 3871.4419 |
| N[IN:3085] | 2937.0777 | 807 | 76.0776 | N[IN:3173] | 2350.7927 | 22,760 | 0.0377 |
| N[IN:3086] | 2206.8133 | 4,238 | 2.0409 | K[IN:3174] | 1431.8157 | 13,333 | 107.1849 |
| N[IN:3087] | 2459.9308 | 315 | 107.0848 |  |  |  |  |

